# Reconstituting Mouse Embryogenesis Ex Utero from Gastrulation to Fetal Development Reveals Maternally Independent Metabolic Programs

**DOI:** 10.64898/2026.03.17.710314

**Authors:** Dmitry Lokshtanov, Shihong Max Gao, Weizhi Xu, Avraham Kosman, Francesco Roncato, Nancy De la Cruz, Nawal A. Khan, Andrea Woods, Ian Campbell, Andrew Woehler, Christina Christoforou, Lang Ding, Amy Hu, Monique Copeland, Lihua Wang, Xiushuai Yang, Castle Raley, Kym M. Delventhal, Antonio Herrera, Alessandro Valente, Sylvia Wright, Elidet Gomez-Cesar, Raanan Shlomo, Sergey Golenchenko, Bernardo Oldak, Alperen Yilmaz, Gulben Gurhan-Sebinc, Mehmet-Yunus Comar, Sergey Viukov, Noa Novershtern, Hua Zhang, Thao Duong, Lingjun Li, Nizar Khatib, Reli Rachel Kakun, Isabel Espinosa-Medina, Maria E. Florian-Rodriguez, Gioele LaManno, Paul W. Tillberg, Meng C. Wang, Itay Maza, Sanjay Srivatsan, Ashley Solmonson, Jacob H. Hanna, Alejandro Aguilera-Castrejon

## Abstract

Mammalian development takes place inside the maternal uterus, creating technological constraints that make difficult the study of embryogenesis in live developing embryos. A central challenge for understanding the role of metabolism in mammalian development is discriminating placental and uterine-regulated signals from embryo-intrinsic processes independent of maternal influence, a process that until now has remained inseparable during gastrulation and organogenesis^1–3^. Ex utero culture systems allowing continuous growth of embryos during pre-gastrulation to organogenesis^4,5^ offer a promising solution to this challenge. Here, we present optimized ex utero culture platforms that support faithful development of mouse embryos from gastrulation (embryonic day 6.5/7.5) through the fetal period (embryonic day ∼12.5) and harnessed these platforms for dissecting metabolic transitions in vivo during embryogenesis independently of uterus and placenta. We characterized the metabolome of in utero and ex utero whole embryos, fetal organs and culture medium between embryonic days E6.5 and E12.5 by liquid chromatography mass-spectrometry (LC-MS) metabolomics, isotope tracing, and single cell transcriptomics. These datasets present a comprehensive overview of the dynamic embryonic metabolism during gastrulation and organogenesis in utero and ex utero. This analysis revealed that the midgestational metabolic switch occurring at E10.5-E11.5 is faithfully recapitulated ex utero, indicating that this transition is intrinsically programmed in embryonic tissues and does not require direct maternal or placental cues. Notably, oxygen availability modulated the extent of this transition, but elevated oxygen was insufficient to induce it prematurely, demonstrating that the switch is developmentally timed and only partially environmental-responsive. We further harnessed the ex utero platform for identifying and perturbing a mitochondrial redox shift at E7.5-E8.5 that is critical for developmental progress after gastrulation. These findings uncover the remarkable metabolic plasticity of the mammalian embryo, demonstrating its capacity to sustain growth independently of maternal inputs from the establishment of the body plan through the onset of the fetal period. Moreover, they highlight the use of long-term ex utero culture as a unique framework for dissecting the mechanisms that shape embryogenesis under physiological and experimentally perturbed conditions, while functionally uncoupling embryonic programs from maternal and placental influences.

## Introduction

The interplay of signals regulating the emergence of a conserved body plan with complex tissues and organs from a mass of seemingly identical cells during animal development remains poorly characterized^6,7^. Understanding this process has been particularly challenging for mammals, where the confinement and inaccessibility of the developing embryo inside the uterus has limited the study of the post-implanted embryo, restricting research on these stages to developmental snapshots that do not permit following the live progress of the process^8–11^. The ability to grow mammalian embryos independently of the uterus for multiple days represents an accessible method for visualizing and manipulating living embryos, as well as exploring embryogenesis independently of fetal-maternal interactions. A variety of culture techniques have been employed in the last century for culturing mouse and rat post-implantation embryos ex utero^12–18^. The use of these platforms has proven instrumental for expanding our knowledge on mammalian development after implantation^19–22^, even though these culture platforms were not highly efficient and did not cover the period from before gastrulation until organ formation^8,23–26^. Recently, culture systems capable of supporting prolonged embryogenesis independently of the uterus from before gastrulation (E5.5/E6.5) to advanced organogenesis (E11.0) were established by incorporating the roller culture on a drum with an electronic gas mixture control and flow/pressure regulation system, combined with defined growth conditions and medium (termed Ex Utero Culture Medium (EUCM))^4,5^.

A major advantage of the ex utero culture platforms is the ability to perform perturbations whose outcomes can be followed over multiple days of development. This opens new possibilities for functional and mechanistic interrogation of mammalian post-implantation development with unprecedented flexibility. Metabolism has emerged as a critical regulator of embryogenesis, helping direct tissue and organ formation beyond its basic role in energy supply for cell function and growth^27–30^. Metabolic switches play a crucial role during mammalian pre- and post-implantation development, as cells differentiate and specialize in nutrient and metabolic requirements to support new tissue functions in response to dynamic physiological and environmental conditions^1,2,27,31^. After implantation, due to the physiologically hypoxic environment in utero, the developing embryo relies primarily on glycolysis to support its growth^32,33^. As the embryo grows, and before the establishment of blood flow through the placenta around E10.5, the yolk sac serves as the major site of nutrient absorption from the maternal environment^34,35^. Previous analyses of mammalian embryonic metabolism at midgestation have relied on genetic loss-of-function experiments, in vitro models, or intrauterine metabolomic measurements^1–3,36^ with few studies able to measure metabolic activities in tissues in vivo. In utero metabolomic analysis coupled to stable isotope tracing has identified embryonic day 10.5–11.5 as a period of metabolic transition for the embryo, characterized by increased purine metabolism and a gradual increase in glucose oxidation in the mitochondria, which have been hypothesized to be driven by the establishment of placental nutrient transfer to the embryo^2^. In studies performed in vivo, it has been difficult to determine if metabolic transitions observed in the embryo arise due to a cue from maternal or placental compartments or if they are intrinsic to embryonic development. There are also difficulties in isolating the response of a metabolic perturbation to a single compartment at specific developmental times due to the dynamic metabolic interplay among maternal, placental, and fetal tissues. Here, we devised an extended ex utero culture system that supports growth of mouse embryos from the onset of gastrulation (E6.5/E7.5 days post-coitum (dpc)) towards the fetal period (E12.5), which allows characterizing and perturbing metabolic transitions occurring during mammalian gastrulation and organogenesis independently of maternal and placental metabolism. Our study reveals metabolic switches that are inherently programmed to the embryo proper during development, triggered independently of placental function or maternal metabolic input.

## Results

### Extended ex utero culture platforms from late gastrulation to the end of organogenesis

To establish an extended mouse embryo culture system, we hypothesized that the limit of embryo growth achieved previously using a maximum value of 21% O_2_ was restricted by oxygen delivery due to tissue size, so, increasing oxygen levels beyond 21% could enhance oxygen availability to internal tissues. Accordingly, we modified our previously described electronic gas regulation culture system integrated with the roller culture on a drum^4,37^ to allow not only precise control of O_2_ and CO_2_ levels between 5 to 20% and gas flow/pressure control, but also oxygen levels beyond atmospheric values (hyperoxia) by incorporating an oxygen gas inlet **(Fig. 1A, Extended data Fig. 1A)**. To facilitate the implementation of our ex utero embryo culture protocol, we additionally tested whether commercial electronic gas mixers with gas flow and/or pressure control (Biospherix OxyStreamer, Eppendorf DASGIP MX4/4, and Okolab tri-gas mixer) could be adapted to support embryo growth as our in-house designed device (Arad Technologies Model 2) **(Fig. 1B, Extended data Fig. 1A-D)**. After testing multiple growth settings using these four systems, we established conditions that support growth of E7.5 embryos (neural plate and early headfold-stage^38^) until the forelimb digit formation stage (∼E12.5, Theiler stage 20) with consistent and reproducible efficiency **(Fig. 1A, Extended data Fig. 1E, 3)**. Importantly, by measuring the gas flowing in the rotating drum, gas relative humidity, as well as the differential pressure inside the drum while connected to the gas mixer module, we found that maintaining a stable gas flow of 30-50 mL per minute, a relative humidity of approximately ∼85-90%, and an internal pressure oscillations of ∼4-6 mbar during the culture supports normal and efficient development in all four gas regulators **(Extended data Fig. 2)**.

**Figure. 1.**
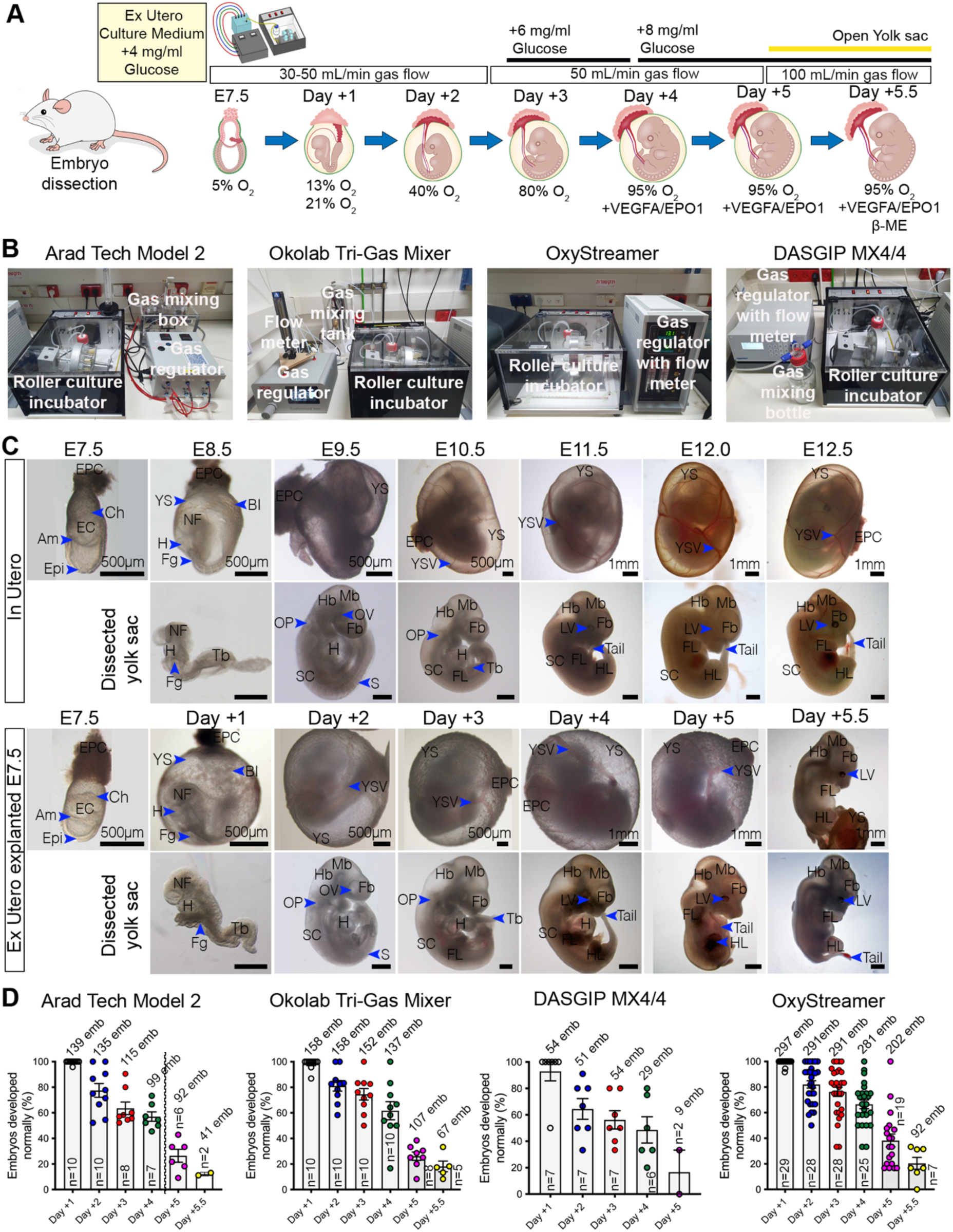
Devising ex utero growth platforms for culturing mouse gastrulating embryos to fetal development. **A,** Schematic overview of the mouse embryo ex utero culture conditions from E7.5 to Day +5.5. Embryos were placed in the roller culture system at E7.5 and sequentially exposed to specific gas compositions, glucose concentrations, and supplemental factors (VEGF-A, EPO, β-ME) at each developmental day. Gas flow rates and oxygen percentages used across the culture period are indicated. **B,** Representative configurations of the electronic gas-mixing modules adapted for the rotating culture incubator. Systems shown include the Arad Tech Model 2, Okolab Tri-Gas Mixer, OxyStreamer, and DASGIP MX4/4. **C,** Bright-field images of embryos developing in utero (top panels) and embryos explanted at E7.5 and cultured ex utero (bottom panels), shown at matched developmental stages from E7.5 to ∼E12.5/Day +5.5. For each stage, embryos are displayed both intact within the yolk sac and after dissection. Scale bars are indicated in each panel. **D,** Percentage of normally developing embryos (see Methods) at each day of ex utero culture per gas regulation system. “emb” denotes the number of embryos assessed; “n” denotes the number of independent experiments. Data represent mean ± s.e.m. Discontinuous line in Arad Tech Model 2 represents embryos transferred to an alternative system due to the 80% O_2_ limit of this setup. Am, amnion; BI, blood islands; Ch, chorion; EC, exocoelomic cavity; Epi, epiblast; EPC, ectoplacental cone; EPO, erythropoietin; Fb, forebrain; Fg, foregut pocket; FL, forelimb; H, heart; Hb, hindbrain; HL, hindlimb; LV, lens vesicle; Mb, midbrain; NF, neural folds; OP, otic pit; OV, optic vesicle; S, somites; SC, spinal cord; Tb, tail bud; VEGF-A, Vascular Endothelial Growth Factor A; YS, yolk sac; YSV, yolk sac vessel.

We employed our previously described ex utero culture medium (EUCM), which supports embryo growth with higher efficiency to E12.5 than DMEM with rat serum alone **(Extended data Fig. 1E)**. Given that glucose represents one of the main metabolic sources following implantation^28,39^ and during mid-gestation^40,41^, we supplemented EUCM with 4 mg/mL Glucose for the first 3 days of culture, followed by an increase to 6 mg/mL at E10.5, and 8 mg/mL after E11.5, which promotes higher embryo survival (**Extended data Fig. 1E**). Similar to previous results^4^, sequential increase in the levels of oxygen, starting from a hypoxic atmosphere of 5% O_2_ at E7.5, 13% at E8.25, and rising to normoxia (21% O_2_) at late E8.75 until E9.5 was most optimal **(Fig. 1A, C)**. We previously found that an environment of 21% O_2_ sustained a maximum growth until E11.0 (Theiler stage 18) after 3.5 days^4^. This is likely due to increased embryo size and the lack of placental connection to the maternal blood supply for facilitating nutrient and oxygen delivery. We found that a gradually increasing hyperoxic environment (40% O_2_ at E9.5, 80% O_2_ at E10.5 and 95% O_2_ at E11.5) can support proper embryo growth until a developmental stage equivalent to ∼E12.0 after 5 days, with the embryo growing inside the yolk sac, and without inducing any visible oxygen damage **(Fig. 1A, C, Extended data Fig. 1E, Fig. 2, Supplementary Video 1)**. In parallel, supplementation with Erythropoietin (Epo) and VEGFA after culture day 4 (∼E11.5), known for their role stimulating differentiation of mature erythrocytes^42^ and blood vessel formation^43^ promoted embryo survival **(Extended data Fig. 1E)**. Embryos with intact yolk sac did not develop beyond day 5 of ex utero culture **(Extended data Fig. 1E)**. Previous studies^44^ have demonstrated that exteriorization of rat embryos from the yolk sac allows short-term embryo culture after 11 gestation days, thus, we tested whether the culture time can be extended beyond 5 days by opening the yolk sac. Careful dissection of the yolk sac to avoid rupture of major blood vessels while maintaining the vascular continuity of the embryo with the yolk sac through the umbilical cord allowed further embryo growth for ∼12 hours, reaching a developmental stage equivalent to E12.5 dpc (Theiler stage 20). **(Fig. 1A, C, Extended data Fig. 1E, Fig. S3A-C, Supplementary Video 2)**.

**Figure 2.**
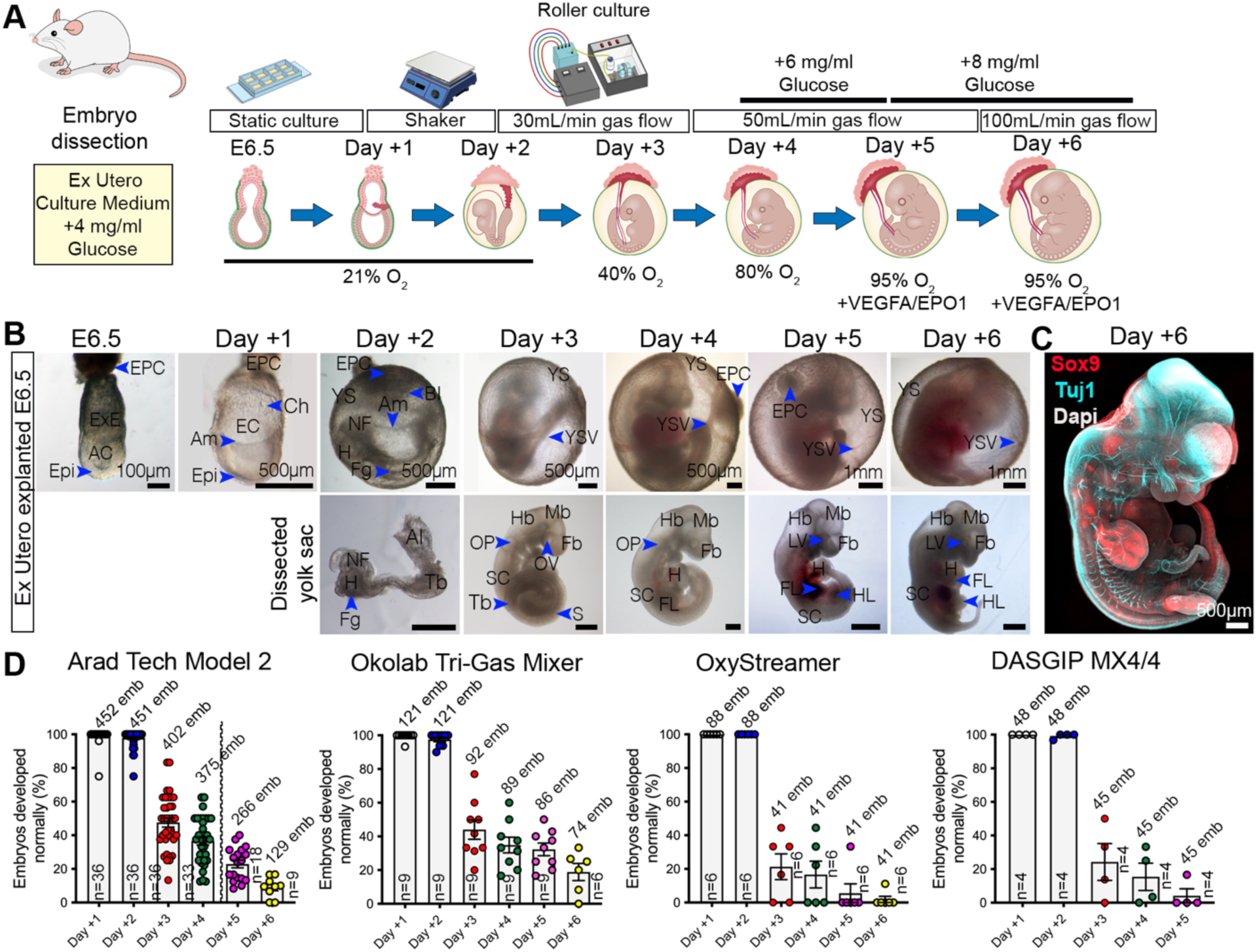
Extended ex utero culture supports development of mouse pre-gastrulation embryos through terminal organogenesis. **A,** Schematic of the six-day sequential culture regimen used to grow embryos ex utero from E6.5. Embryos were initially maintained in static culture for 24 hours, then moved to a rotating shaker on day 1, and subsequently transferred to the roller bottle culture integrated with electronic gas-regulation systems. Oxygen levels and glucose supplementation were progressively adjusted during development as indicated. **B,** Bright-field images of E6.5 embryos grown ex utero from day +1 to +6. Embryos cultured beyond day two are shown both intact within the yolk sac (top panels) and after dissection (bottom panels). Scale bars are indicated on each panel. **C,** Representative whole-mount immunostaining of a day +6 ex utero-grown embryo stained for Sox9 (red), Tuj1 (cyan), and DAPI (white) illustrating proper spatial organization of skeletal progenitors and peripheral/central nervous system structures. Scale bar = 500 µm. **D,** Percentage of embryos displaying morphologically normal development (see Methods) when cultured from E6.5 across four gas-regulation platforms. “emb” denotes total number of embryos assessed. “n” indicates number of independent experiments. Data represent mean ± s.e.m. Discontinuous line in Arad Tech Model 2 represents embryos transferred to an alternative system due to the 80% O_2_ limit of this setup. Images represent a minimum of 3 biological replicates. AC, amniotic cavity; Am, amnion; BI, blood islands; Ch, chorion; EC, exocoelomic cavity; Epi, epiblast; EPC, ectoplacental cone; EPO, erythropoietin; ExE, extraembryonic ectoderm; Fb, forebrain; Fg, foregut pocket; FL, forelimb; H, heart; Hb, hindbrain; HL, hindlimb; LV, lens vesicle; Mb, midbrain; NF, neural folds; OP, otic pit; OV, optic vesicle; S, somites; SC, spinal cord; Tb, tail bud; VEGF-A, Vascular Endothelial Growth Factor A; YS, yolk sac; YSV, yolk sac vessel.

To assess embryonic stage and efficiency of appropriate embryo development ex utero we employed previously defined embryo scoring systems based on morphological traits^45–48^ **(Fig. 1C, Supplementary Video 1, see methods)**. These features included: somite number, embryo size, yolk sac circulation, allantois development, axial turning of the embryo, heart development, neural tube closure, brain regionalization, development of optic and olfactory systems as well as branchial arches, forelimbs and hindlimbs. Embryos between E8.0 and E10.5 were predominantly staged by somite counting, while for E11.5 and afterwards we additionally employed hindlimb staging. During the first 3 days of ex utero development (E7.5-E10.5), there were not visible differences between embryos cultured in previously reported conditions under normoxia^4^ and hyperoxic conditions **(Fig. 1C)**, which indicates tolerance of embryos to variable oxygen levels. At the fourth day of culture, embryos reached a stage equivalent to E11.2/E11.5 (42-45 somites, Theiler stage 19), while after 5 days of ex utero growth inside the yolk sac the embryos matched their ∼E12.0 in utero counterparts **(Fig. 1C, Extended data Fig. 3A-C)**. Maximum embryo growth is reached after 5.5 days of culture, equivalent to ∼E12.5 (∼48-49 somites, Theiler stage 20), when the earliest signs of digits in the forelimb are visible **(Fig. 1C, Extended data Fig. 3A, B)**. These culture platforms yield ∼60-75% proper embryo development after 3 days of culture, ∼50-60% after 4 days, 15-30% after 5 days, and 10-20% at 5.5 days of ex utero development in different mouse strains **(Fig. 1D, Extended data Fig. 3D, E)**. Culture efficiency is significantly higher in outbred strains CD1 and CFW **(Extended data Fig. 3D)**, which corresponds to their superior fertility observed in utero^49^. The crown-rump length of the cultured embryos was slightly reduced although comparable to matched in utero embryos (**Extended data Fig. E**). The efficiency and reproducibility of the ex utero culture protocol was validated in an independent laboratory **(Extended data Fig. 3F)**. As previously observed in short-term cultures^44,50^, embryos after E11.5 (culture day 4 in our setting) grow at a slower rate compared to in utero controls, suggesting that speed of development is influenced by maternal and chorioallantoic placenta nutrition. After 5.5 days the embryos show evident abnormalities, mainly enlarged heart, pericardial edema, internal hemorrhage, and necrosis of internal tissues due to lack of oxygen and nutrient delivered to the internal parts of the body.

### Embryo culture platforms support extended ex utero development from pre-gastrulation stages

Next, we evaluated whether the newly adapted gas regulators and extended culture protocol support growth of embryos from pre-gastrulation (E6.5 dpc). We employed the EUCM as described above combined with the accurate control of gas flow and oxygen concentration of the gas control systems described for E7.5 culture. Consistent with previous findings^4^, only a combination of static and roller culture protocols in a 21% oxygen atmosphere during the static period allowed proper development of embryos beyond E8.5 **(Fig. 2A, Extended data Fig. 4A)**. Embryos are first cultured in static plates for a day, followed by static culture on a rotation shaker from E7.5 to E8.5, which we found helps supporting growth of E8.5 embryos for longer times (1-2 somite pairs) compared to non-shaking culture. At day 2, early somite-stage embryos (4-9 somites) were relocated from shaking culture to the rotating bottles culture for testing further growth in each one of the four electronic gas regulator systems **(Fig. 2A)**. Transfer of E6.5 explanted embryos to rotating conditions at E6.5 or E7.5 failed to support adequate growth beyond 2 days **(Extended data Fig. 4A)**. The in-house designed Arad Technologies system and the Okolab Tri-Gas Mixer supported robust embryo growth ex utero culture for up to six days (two days of static culture and four additional culture days in rotating system), which may be due to the fact that under our configurations these two systems regulate gas pressure and flow, while the OxyStreamer and Eppendorf DASGIP systems mostly control gas flow. We did not find significant differences in media pH or osmolality which could affect embryo development due to an accumulation of waste products in static culture compared to rotating bottles system or between gas regulation systems **(Extended data Fig. 4F-G)**. Embryos cultured from E6.5 using the established protocol reached a maximum developmental stage comparable to ∼E11.75 in utero developed embryos (Theiler stage 19) **(Fig. 2A, B, Supplementary Video 3)**. During the first two days of static culture our ex utero protocol yield adequate embryo growth with 95-100% efficiency, while after transfer to the rotating culture system (Arad Tech/Okolab) culture efficiency ranges from 40-50% at day 3, 35-40% at day 4, 20-30% at day 5 and 12-18% at the last day of culture **(Fig. 2C)**. Despite a slight developmental delay of 2-4 somite pairs at day 3 and 4 of culture, which was more pronounced after 5 and 6 days (4-6 somites), embryogenesis proceeded similarly to in utero controls **(Fig. 2B)**. The latter observation suggests that longer culture periods (and the earlier the embryos are explanted) correlate with the slower developmental tempo. Whole-embryo immunostaining for the globally expressed genes SOX9 (bone/cartilage marker) and TUJ1 (pan-neuronal marker) showed a similar pattern to in utero development, demonstrating formation of the digit domains in the anterior footplate **(Fig. 2B)**. Besides, only embryos obtained from outbred mouse strains (CD1 (ICR), CFW) or mixed backgrounds (CD1 x BDF1) are able to grow properly beyond two days of culture if explanted at E6.5 **(Fig. 2B, C, Extended data Fig. 4B, C)** which relates with the fact that inbreeding depression leads to poor fecundity^51^. In contrast to cultures starting at E7.5 in which human adult serum (HAS) and human umbilical cord blood serum (HCS) showed no evident differences in growth efficiency, embryos cultured from E6.5 showed a higher percentage of properly grown embryos after three days of culture when supplemented with HCS **(Extended data Fig. 4D)**. Comparison of the crown-rump length of ex utero and in utero embryos showed a parallel increase in size between cultured embryos (explanted at E7.5 or E6.5) and controls developed in utero, reaching a maximum size of ∼8-9 mm at the final stage of growth **(Extended data Fig. 4E)**. Embryos cultured ex utero from E6.5 showed significantly smaller size (ranging from 300 to 500µm) to in utero embryos **(Extended data Fig. 4E)**, which could be explained given the delay observed in speed of developmental events when embryos are grown from pre-gastrulation.

### Ex utero embryogenesis recapitulates natural 3D tissue architecture of developing organs and tissue-wide single cell transcriptome

We next compared the development of major internal organs at the molecular and architectural level in ex utero and in utero developed embryos from E11.5 to E12.5 (day 4 to 5.5 of ex utero culture from E7.5) by whole-mount immunostaining. We focused on developmental markers representing tissues derived from the three embryonic germ layers. At E11.5/E12.5, SOX9 is widely expressed in the mesenchyme condensations of the skull, the branchial arches and their subsequent derivatives, the developing ear, and in the paraxial mesoderm within the developing vertebrae and ribs, genital ridges, and in the limbs, where it validates proper initiation of digit formation in the forelimb and hindlimb, although hindlimb development in ex utero-cultured embryos was slightly delayed relatively to E12.5 in utero controls **(Fig. 3A, B)**. SOX9 is also expressed in the pancreatic progenitors, revealing differentiation of pancreatic buds in both ex utero and in utero embryos **(Fig. 3A, B)**. Developing neurons in the peripheral and central nervous system were marked by Tuj1, showing proper innervation of organs and limbs ex utero **(Fig. 3A, B)**. In both in utero controls and ex utero embryos, SOX2 was observed in the brain, spinal cord, foregut and stomach **(Fig. 3C, D)**, while NKX2-1 labeled the trachea, lung primordia, as well as the regions of the telencephalon and hypothalamus **(Fig. 3C, D)**. PAX6 labeling demonstrates adequate development of eyes, forebrain, hindbrain and spinal cord ex utero **(Fig. 3E, F)**. Embryonic heart and muscle bundles along the vertebrae follow similar patterns in control embryos and ex utero embryos, evidenced by MHC-II staining. The distribution of neural crest-derived tissues (SOX10/ISL1^+^) across ex utero embryos, as well as endothelial cells lining the blood vessels (identified by CD31), were analogous to their in utero counterparts **(Extended data Fig. 5)**. Whole-embryo 3D reconstructions of immunostained samples using light-sheet microscopy showed that, despite a slight developmental delay, the 3D tissue architecture of the ex utero-grown embryos is comparable embryos developed in the womb, with only MHC-II showing differences in the orientation of muscle bundles and a slightly enlarged heart **(Supplementary Video 4)**. Taken together, our molecular and 3D histological analyses suggest that ex utero cultured embryos consistently recapitulate spatiotemporal gene expression patterns observed in embryos developed in utero until approximately Theiler stage 20 (48-49 somites stage), similar to E12.5 days of development.

**Figure 3.**
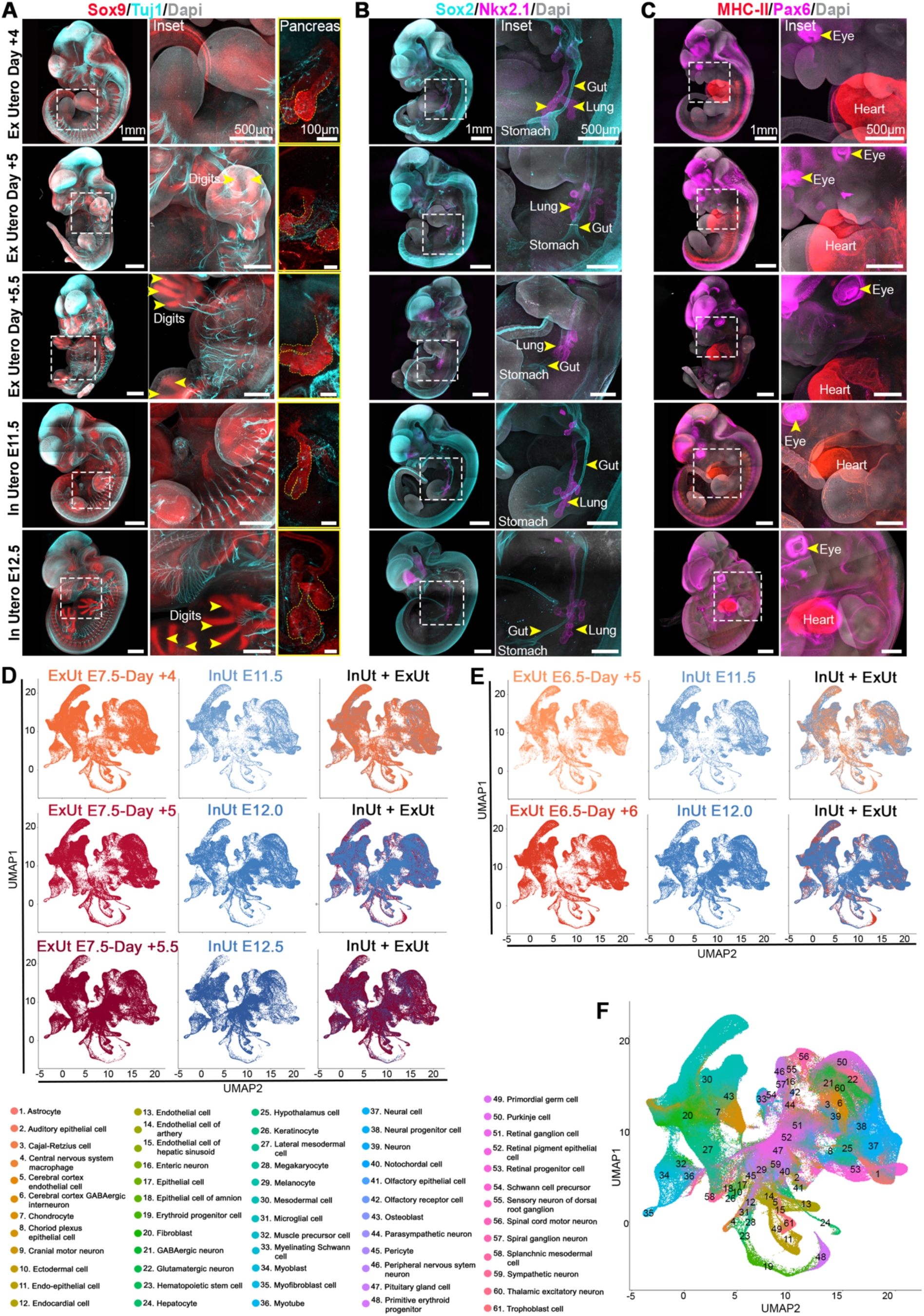
Tissue architecture and spatio-temporal expression profiles of developing organs are recapitulated in ex utero embryos. **A,** Representative whole-mount immunostaining images of embryos developed ex utero and in utero time-matched controls showing body-wide neurons (Tuj1, cyan) and cartilage/bone progenitors (Sox9, red). Insets highlight limb digit patterning and pancreatic bud morphology (right panels). **B,** Whole-embryo maximum intensity projections of ex utero and in utero embryos stained for Sox2 (cyan), identifying the developing nervous and digestive systems, and Nkx2.1 (magenta), labeling the lung primordia, telencephalon and hypothalamus. **C**, Whole-embryo immunostaining for Pax6 (magenta) and MHC-II (Myosin Heavy Chain-II; red) in ex utero and in utero embryos at the indicated stages. Pax6 demarcates the embryonic forebrain, spinal cord, and eye, while MHC-II labels the developing heart and muscle tissues. White, DAPI. Insets are enlargements of the dashed boxes. Yellow arrows denote highlighted developed tissues. Scale bars are indicated on each panel. Images represent a minimum of 3 biological replicates. **D-F,** Single nucleus RNA-transcriptomics characterization of ex utero culture embryos explanted at E6.5 or E7.5. Embryos extracted at E7.5 or E6.5 were cultured for multiple days (day +4, +5, +5.5 or +5, +6) to reach developmental endpoints, then subjected to nuclei isolation followed by single-cell combinatorial indexing (sci-seq3). **D,** UMAP embeddings displaying individual cells isolated from embryos cultured ex utero starting at E7.5 and age-matched in utero controls. Left panels: ex utero samples at culture day +4 (orange), day +5 (dark red), and day +5.5 (maroon). Middle panels: time-matched in utero controls at E11.5, E12.0, and E12.5 (blue). Right panels: merged ex utero and in utero samples showing overall similarity in cellular composition. n = 164,224 cells from three day +4 ex utero embryos; n = 129,569 cells from three day +5 ex utero embryos; n = 205,016 cells from four day +5.5 ex utero embryos; n = 39,313 cells from E11.5 in utero embryos; n = 80,128 cells from E12.0 in utero embryos; n = 80,921 cells from E12.5 in utero embryos. **E,** UMAP embeddings of ex utero cultured embryos starting at E6.5 and stage-matched controls. Left panels: ex utero samples at culture day +5 (orange) and day +6 (red). Middle panels: in utero controls at E11.5 and E12.0 (blue). Right panels: UMAP displaying merged ex utero and in utero samples. n = 48,737 cells from three day +5 ex utero embryos; n = 98,206 cells from three day +6 ex utero embryos; n = 39,313 cells from E11.5 in utero embryos; n = 80,128 cells from E12.0 in utero embryos. **F,** UMAP colored by cell type annotation via scANVI transfer learning from reference atlas^58^. Each color represents a distinct cell type with legend showing 61 identified cell types. UMAP plots show all cells considered in the analysis (n = 645,752 ex utero; n = 200,362 in utero).

We then analyzed the histological composition of placental tissues developing ex utero. The murine placenta is composed of three distinct cellular regions: the labyrinth, the junctional zone and the decidua^52^. Histological examination of coronal sections in whole embryos plus extraembryonic tissues (including yolk sac and ectoplacental cone) revealed that ex utero cultured embryos develop a small portion of the labyrinth placenta, with a network of blood vessels identified by CD31 immunostaining, although the junctional zone and decidua are mostly absent, given the lack of connection with the maternal uterus **(Extended data Fig. 5C)**. Further, as a functional test for ex utero developed tissues, we harnessed in vitro gametogenesis protocols^53^ to analyzed whether gonads isolated from cultured embryos preserve the capacity to properly generate postnatal oocytes **(Extended data Fig. 5D)**. Isolated female gonads and mesonephros from ex utero samples resemble gonads from in utero embryos, both morphologically and molecularly, as identified by GATA4 and WT1 **(Extended data Fig. 5E)**. After 17 days in culture, developmental chronological age of in vitro cultured ovaries corresponds to postnatal day (P)10 pups^54^; at this stage, ovaries were dissociated into single secondary follicles, which are comparable to those derived from in utero isolated gonads, both being surrounded by cumulus cells and displaying the characteristic cytoplasmic expression of STELLA **(Extended data Fig. 5F).** These results demonstrate that ex utero embryos can develop functional gonads that give rise to postnatal oocytes.

To comprehensively characterize the variety of cell types constituting the whole embryo in a quantitative and unbiased manner, we analyzed the transcriptional profile of single cells isolated from ex utero grown embryos and stage-matched in utero controls by single cell RNA sequencing **(Fig 3D-G, Supplementary Video 5)**. Considering that the estimated number of cells per embryo at E11.5 is ∼2.6 million and ∼6 million at E12.5^55^, we employed combinatorial cellular indexing, or ‘sci-RNA-seq3’, which uses specimen multiplexing to allow profiling the transcriptome of millions of nuclei, thus enabling organism-scale single-cell analyses^55,56^. We collected individual embryos grown ex utero at five different timepoints, spanning from embryos explanted at E7.5 and cultured during 4, 5 and 5.5 days, or embryos cultured from E6.5 till day 5 and 6 of ex utero development, as well as in utero controls at E11.5, E12.0, E12.5, and E13.5 dpc as reference controls. Low quality cells (<400 genes) were removed from the analysis, and those with high mitochondrial content were excluded using the miQC package. After quality control our dataset included ∼0.85 million single nuclei, 645,752 nuclei from sixteen ex utero embryos, and 200,362 from five in utero controls, with a median of 2029 UMIs (833 genes) per cell **(Extended data Fig. 6)**.

Samples across all developmental stages were integrated using Concord^57^, a dimensionality reduction contrastive learning-method designed for improving the resolution of cell states and mitigating batch artifacts in single-cell analysis. Uniform Manifold Approximation and Projection (UMAP) plots show a complete overlap in the profile of cell states between embryos developing ex utero from E7.5 through organogenesis until fetal development (E7.5 + 4, 5 and 5.5 days) and equivalent in vivo counterparts **(Fig 3D)**. Remarkably, ex utero embryos grown for 5.5 days starting from E7.5 (mid-gastrulation) are transcriptionally indistinguishable from E12.5 in utero embryos, which demonstrates that the complexity of cell lineage differentiation is faithfully recapitulated at the single-cell level in the ex utero conditions **(Fig 3D)**. Similarly, the UMAP-based transcriptional profile of cell types found in embryos cultured from E6.5 displayed high correspondence to in utero controls, with embryos grown for 5 days resembling E11.5 embryos, whereas those grown for 6 days approached the transcriptional state of E12.0 embryos. Nonetheless, embryos cultured ex utero from pre-gastrulation stages for six days exhibited some noticeable deviations in cell type composition compared to their in utero counterparts **(Fig 3E)**. Cells were annotated using transfer learning from a comprehensive mouse development reference atlas^58^ via scANVI^59^ (which annotates cell types independently of their assigned cluster). We selected data corresponding to E11.5 to E12.5 (a total of 1.5 million cells) from the reference dataset for annotation transfer to our query samples. The annotated cell clusters represent 61 distinct cell types comprising differentiated cells across tissues derived from all three germ layers, including specialized neurons, olfactory and auditory sensory cells, immune and epithelial lineages, as well as diverse muscle and endothelial cell types, consistent with the advanced fetal stage of the embryos **(Fig 3F, G, Extended data Fig. 7)**. All annotated cell types present in the in utero control embryos were also detected in the ex utero cultured samples **(Fig 3F, G)**. Quantitative comparison of the proportion of cell types in the embryos revealed no significant differences between ex utero and in utero conditions across most annotated clusters, with only two cell types (mesodermal cells and myotube) showing a higher representation ex utero, whereas four lineages (thalamic and sympathetic neurons, pituitary gland cells, and retinal pigment epithelial cells) displayed a reduction in abundance compared to intrauterine development **(Extended data Fig. 8)**. Notably, in embryos explanted at E7.5, these populations reached levels comparable to their E12.5 in utero counterparts by day 5.5, suggesting that the observed deficit likely reflects a transient developmental delay rather than a failure to specify these lineages. **(Extended data Fig. 8B)**. Correlation analysis of differentially expressed genes showed very high correlation (R>0.9) in most cell lineages across ex utero embryos cultured either from E7.5 or E6.5 when compared to E12.5 or E12.0 natural embryos, respectively **(Extended data Fig. 9-10)**. Only five cell clusters displayed correlation values between 0.9 and 0.82 for embryos explanted at E7.5 cultured for 5.5 days, while 18 cell clusters showed slightly higher variability (R=0.76-0.9) for embryos grown for six days from E6.5. Finally, to evaluate cell lineage allocation and tissue morphogenesis at the transcriptomic level, we performed cellular-resolution spatial transcriptomics using both Stereo-seq^60^ and Curio trekker^61^ platforms **(Extended data Fig. 11)**. Single mid-sagittal sections of E7.5 embryos cultured ex utero till day 5.5 were compared to E12.5 embryos developed in utero **(Extended data Fig. 11A)**. All major annotated cell lineages and gene markers were similarly allocated between in utero and ex utero embryos, despite technical differences in the anatomic section sampled **(Extended data Fig. 11B, C)**. Taken together, these findings demonstrate that embryos developing ex utero using the culture systems described herein recapitulate the formation of all major tissues and organs from gastrulation through the completion of organogenesis, mirroring in utero development despite the absence of maternal interaction.

### Metabolomic profiling of mammalian ex utero development identifies conserved metabolic transitions

The long-term ex utero embryo culture systems described herein provide the opportunity to uncouple maternal and placental metabolism from the embryo proper during gastrulation through the end of organogenesis. First, to understand the differences in nutrient availability in the ex utero environment, we analyzed the metabolite composition of complete fresh EUCM and individual sera compartments (rat serum (RS), human adult serum (HAS), and human umbilical cord blood serum (HCS)) by liquid chromatography mass-spectrometry (LC-MS) metabolomics. This analysis revealed distinct nutrients contributed by rat and human serum **(Fig. 4A)**. Rat serum was enriched for metabolites associated with the urea cycle (argininosuccinate, citrulline), pyrimidine nucleosides (cytidine, 5-methylcytidine, deoxycytidine) and amino acid metabolism (aspartate, glutamate), which suggest support of rapid growth and nitrogen balance related to nucleotide synthesis. In contrast, human serum displayed higher levels of amino acid catabolic intermediates (ketoleucine, 2-hydroxybutyrate) some of which are derived from the gut microbiome (phenylacetylglutamine, indole-3-lactate) suggesting there may be underexplored relationships between maternal gut metabolism and fetal development **(Fig. 4A)**. Although there are also likely additional differences in cytokines and hormones, the differences in metabolite composition of both sera compartments may provide a plausible explanation for the improved developmental outcome observed in culture with EUCM compared to rat serum only **(Extended data Fig. 1E)**.

**Figure 4.**
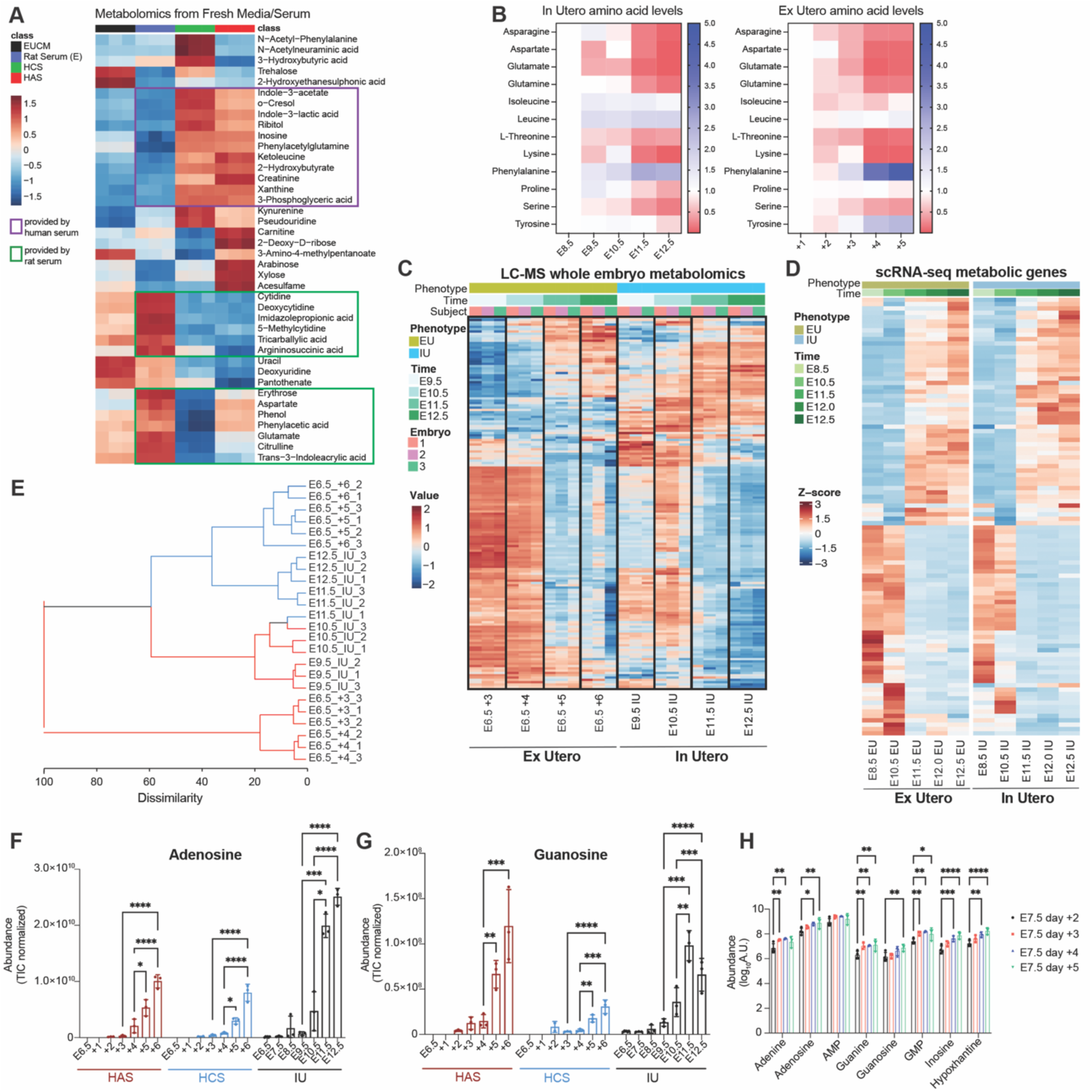
Metabolomic analysis of nutrient availability and metabolic transitions during ex utero embryogenesis. **A,** Heatmap of top 40 metabolites significantly different between complete EUCM and its individual serum components. Distinct nutrient contributions from each serum fraction are indicated in the colored boxes. **B**, Metabolomic analysis of amino acid changes in ex utero cultured embryos from E7.5 though five culture days compared to in utero time-matched controls. In utero controls (left panel) and ex utero embryos grown in HCS (middle panel) are shown. Values are fold change normalized to culture day 1 (E8.5). **C**, Whole-embryo metabolomic profiling demonstrates that the E10.5-E11.5 metabolic transition is recapitulated ex utero. Embryos explanted at E6.5 and cultured for six days were analyzed by LC-MS alongside in utero controls. Heatmap displays 160 evaluated metabolites normalized to total ion content. **D**, Heatmap of metabolic enzyme genes with the highest gene-metabolite correlation (top 100, |*r*| ≥ 0.7) reveals a corresponding transcriptional transition at E10.5-E11.5 in both in utero and ex utero embryos. Genes were mapped to metabolites using KEGG pathway annotations and selected based on Pearson correlation between gene expression and metabolite levels across matched developmental stages. Hierarchical clustering (Euclidean distance, complete linkage) was applied to rows. Z-score normalized expression values are shown. Heatmap incorporates datasets from E8.5, E10.5, E11.5, E12.0, and E12.5; E8.5 datasets were obtained from a prior study^4^. EU, Ex Utero; IU, In Utero. **E**, Hierarchical clustering of global metabolomic signatures showing that ex utero embryos grown in HCS cluster by developmental stage similarly to in utero embryos. **F-G**, Relative abundance of adenosine (**F**) and guanosine (**G**) through development in ex utero cultured embryos (explanted at E6.5 and cultured in HCS or HAS) compared to in utero time-matched controls. Data are mean ± S.D. **H**, Relative increase in abundance of eight purine-derived metabolites through culture time in ex utero embryos cultured from E7.5. Data represent mean ± S.D. * = p<0.05, ** = p<0.01; *** = p<0.001; **** = p<0.0001.

Next, to evaluate the extent in which embryonic metabolic programs observed in utero^1,2^ are preserved independently of the maternal environment, we performed untargeted metabolomics characterization of whole ex utero-grown embryos and their corresponding culture media by LC-MS, which allow simultaneous assessment of the embryo metabolic state and nutrient utilization dynamics. Embryos explanted at either E6.5 or E7.5 were subjected to ex utero culture, with whole embryos and culture media collected every 24 hours and compared to corresponding controls developing in the maternal uterus. Given the differences in embryo growth efficiency observed for embryos explanted at E6.5 when grown of using HAS or HCS supplementation, we characterized embryos in both conditions.

Comparison of amino acid levels in the embryo during organogenesis from E8.5 to E12.5 showed a parallel progressive and stage-dependent decrease of amino acids in cultured embryos mirroring in utero controls **(Fig. 4B)**. While embryos cultured in HCS closely recapitulated the amino acid signatures observed in utero, amino acid levels of those embryos grown in HAS showed minor deviations in amino acid abundance over time **(Extended data Fig. 12A)**. These differences cannot be attributed to lower amino acid availability in HAS, which presents overall higher amino acid concentration compared to HCS **(Extended data Fig. 12B)**. Interestingly, analysis of amino acid abundance in consumed media through a 24 hour time course on E10.5 revealed that consumption of some amino acids (i.e., Glutamate) increases over time, while others (i.e., Aspartate) are increasingly excreted to the culture media as organogenesis proceeds, despite their high abundance in culture media **(Extended data Fig. 12C)**. Glutamate and aspartate are significantly increased in HAS compared to HCS **(Extended data Fig. 12B)** and this corresponds to changes in these amino acids in embryos grown in each media, with glutamate and aspartate increasing in HAS embryos but decreasing in HCS embryos from E8.5 **(Fig. 4B, Extended data Fig. 12B)**. These data are consistent with embryos grown in HCS being more metabolically similar to in utero embryos compared with HAS **(Extended data Fig. 12D)**. Overall, these results indicate that ex utero cultured embryos accurately recapitulate physiological amino acid dynamics during organogenesis and that media composition can influence metabolic activities, providing an opportunity to interrogate how the embryo responds directly to changes in nutrient abundance, an approach that cannot be done in vivo.

To investigate the potential metabolic origin of defects observed in ex utero cultured embryos, we profiled the metabolome of abnormal embryos at E9.5 (E6.5 day +3) showing a range of morphological defects. Analysis of the top 75 differential metabolites revealed a marked metabolic divergence between normally developed embryos and embryos exhibiting morphological defects **(Extended data Fig. 12E)**. Properly developed embryos at E9.5 displayed a metabolomic profile characterized by high abundance of glycolytic intermediates, glycerophospholipids, and nucleotide-related metabolites, consistent with active anabolic growth **(Extended data Fig. 12E)**. Defective embryos showed a broad depletion of central carbon metabolites, multiple amino acids, NAD⁺, UDP-sugars, and acylcarnitines, suggesting impaired glycolytic activity, compromised amino-acid homeostasis, diminished mitochondrial oxidation, and defects in lipid-derived energy metabolism **(Extended data Fig. 12E)**. Conversely, a subset of breakdown products and stress-associated metabolites accumulated specifically in defective embryos, pointing to increased catabolic activity and metabolic dysregulation **(Extended data Fig. 12E)**. Together, these data reveal that successfully developed embryos maintain a coordinated metabolic program that supports nucleotide synthesis, redox balance, and biosynthetic flux, whereas morphologically defective embryos exhibit global metabolic collapse marked by reduced carbohydrate utilization, impaired amino-acid metabolism, and perturbed mitochondrial function.

Previous studies^2^ have described fundamental transitions in metabolic profiles of the embryo and placenta occurring between E10.5 and E11.5, with both tissues undergoing simultaneous but distinct changes. The synchronized timing of these transitions was largely attributed to increases of placental function after E10.5, resulting in increased transfer of nutrients and oxygen from the maternal circulation as the placental vasculature matures and becomes functional^2,52^. To determine whether global metabolic transitions characteristic of midgestation (E10.5-E11.5) are preserved ex utero, we performed whole-embryo metabolomic profiling of embryos explanted at E6.5 and cultured for six days, alongside in utero controls at matched stages. Remarkably, despite the absence of maternal and placental input, ex utero embryos underwent a pronounced metabolic transition between E10.5 and E11.5, closely mimicking the metabolic switch observed in utero **(Fig. 4C)**. In both in utero and ex utero conditions, this transition was characterized by coordinated increase in purine metabolites (adenosine, inosine, guanosine, urate), TCA-cycle intermediates (citrate/isocitrate, malate, succinate), amino acid metabolism (glutamate, aspartate, and arginine–urea cycle intermediates), and redox-associated metabolites **(Fig. 4C)**. Consistent with these findings, hierarchical clustering of global metabolomic signatures demonstrated that ex utero embryos cultured in HCS clustered closely to in utero developed embryos by developmental stage **(Fig. 4E),** highlighting the metabolic similarities of ex utero cultured embryos to intrauterine development despite of absence of maternal nutrition. The metabolic changes observed during this transition are indicative of an embryo-wide shift toward increased mitochondrial oxidative metabolism, consistent with previously identified features of the E10.5-E11.5 metabolic transition in utero^2^. Moreover, since elevated purine metabolism represents a defining feature of this metabolic transition^1,2^, we directly compared purine metabolite dynamics between in utero embryos and those cultured ex utero from E6.5 in either media supplemented with HCS or HAS. In both ex utero conditions, a significant increase in purine derivatives (adenosine, guanosine, etc.) was detected from E10.5, closely mirroring the increase observed in utero **(Fig. 4F-H)**. Consistent with a rise in purine biosynthesis during the E10.5-E11.5 transition, [U-¹³C]glucose tracing revealed increased incorporation of glucose-derived carbon into AMP and GMP ex utero over a 12-hour period, resembling what was previously reported in vivo^4^ **(Extended data Fig. 13A)**. Although the E10.5-E11.5 transition is maintained in embryos grown from E6.5 in both HCS and HAS, the overall metabolomic signature of embryos grown in HCS is more similar to in utero than those embryos cultured in HAS, which are grouped more strongly by culture environment than by developmental stage **(Extended data Fig. 12D)**. While the global structure of the transition was preserved ex utero, cultured embryos exhibited increased accumulation of purine degradation products (e.g., urate), suggesting that the placenta may play a significant role in purine metabolite clearance in vivo **(Extended data Fig. 12F, 13B**).

Finally, to assess whether the metabolic transition was accompanied by coordinated transcriptional remodeling, we integrated metabolomic data with gene expression profiles of metabolic enzymes by KEGG mapping^62^. To this aim, we incorporated the sci-RNA-seq3 dataset obtained in this study, covering E11.5, E12.0 and E12.5 mouse embryos, with our previous dataset of ex utero and in utero embryos at E8.5 and E10.5^4^. Analysis of the top 100 enzyme-coding genes whose expression levels were strongly correlated with associated metabolite abundance (|r| ≥ 0.7) revealed a sharp transcriptional transition at E10.5-E11.5 in both in utero and ex utero embryos **(Fig. 4D)**. The same transition was also observed when all 370 genes identified by KEGG were considered in the analysis **(Extended data Fig. 14A)**. The concordance between metabolic and transcriptional profile changes supports the existence of a tightly regulated developmental metabolic program that is retained ex utero. Collectively, these results demonstrate that embryos developing ex utero recapitulate intrauterine metabolic programs and establish that the E10.5-E11.5 metabolic transition represents an embryo-intrinsic developmentally programmed shift rather than a response to maternal or placental nutrient inputs at both transcriptomic and metabolomic level.

### Tissue-level characterization of embryonic metabolome at midgestation

We next aimed to incorporate tissue-level spatial information to our whole-embryo metabolomics analyses dataset by applying tissue expansion mass-spectrometry imaging (TEMI)-based spatial metabolomics^63^. Mid-sagittal sections from day 5.5 ex utero embryos and E12.5 in utero controls were expanded approximately 3.5-fold and analyzed by TEMI, which provides higher resolution compared to unexpanded samples **(Extended data Fig. 15)**. Mass spectrometry imaging of 39 metabolites and lipids spanning m/z values from 184.07 to 810.59 revealed that most molecules analyzed present comparable organ-scale distribution patterns between ex utero and in utero embryos, despite differences in the precise sagittal plane sampled **(Fig. 5A, Extended data Fig. 15)**. Putative molecular annotation based on m/z values enabled the identification of 19 metabolites, including Phosphorylcholine, Carnitine, and a variety of Lysophosphatidylcholines (LPCs), Phosphoethanolamines (PEs), Sphingomyelins (SM), and Phosphatidylcholines (PCs), which exhibit differential distributions across embryonic tissues **(Fig. 5A, Extended data Fig. 15)**.

**Figure 5.**
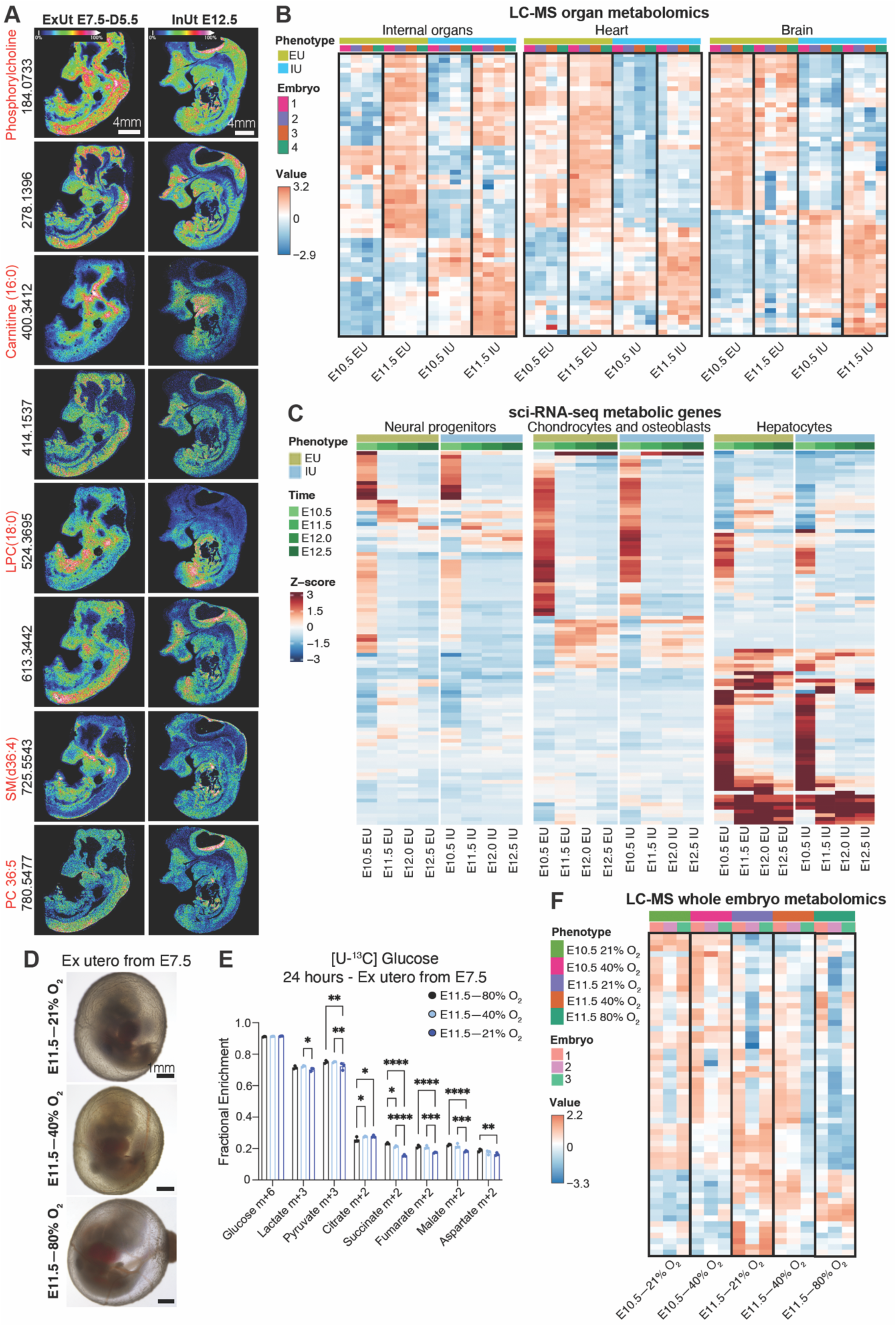
Tissue-level characterization of ex utero embryo metabolism and perturbations of the midgestation metabolic transition. **A**, Mass spectrometry images of ∼3.5 fold expanded (TEMI) embryos grown ex utero for 5.5 days starting from E7.5, compared to E12.5 in utero control embryos. Eight selected metabolites/lipids with *m/z* values ranging from 184.07 to 780.54 are shown. Putative metabolite identities are indicated in red above the corresponding image. All the MSI images were obtained using a 50-µm laser raster scanning with a mass error tolerance of 10 ppm. Scale bars, 4 mm. Color bar indicates the ion intensity scale ranging from 0 to 100%. **B,** Heatmaps displaying organ-level LC-MS metabolomic profiles for 58 metabolites evaluated on isolated internal organs, heart and brain dissected at E10.5 and E11.5 in ex utero and in utero embryos. Metabolites were normalized to total ion content. EU, Ex Utero; IU, In Utero. **C,** Cell type-specific heatmap showing normalized expression values of top 100 metabolic enzyme genes (|*r*| ≥ 0.7) identified by KEGG pathway mapping for neural progenitors, osteoblast/chondrocytes, and hepatocytes. Rows indicate metabolism-related genes identified by KEGG pathway mapping. Color scale represents normalized z-score expression values. Hierarchical clustering of genes was performed using Euclidean distance and complete linkage. EU, Ex Utero; IU, In Utero. Heatmap incorporates datasets from E10.5, E11.5, E12.0, and E12.5 due to differences in lineages present at early stages; E10.5 datasets were obtained from a prior study^4^. **D,** Representative bright-field images of embryos explanted at E7.5 and cultured ex utero under different oxygen concentrations (21%, 40%, 80%) during the period of the midgestation metabolic switch at E10.5 to E11.5. Scale bars, 1mm. **E,** [U-^13^C]-Glucose tracing for 24 hours on embryos cultured ex utero under 21%, 40% or 80% O₂ from E10.5 to E11.5. Data are mean ± S.D. of 3 biological replicates. * = p<0.05, ** = p<0.01; *** = p<0.001; **** = p<0.0001. **F,** Whole embryo metabolomic profiling of the midgestation metabolic transition under variable oxygen tensions. Heatmaps of embryos cultured ex utero and exposed to 21%, 40% or 80% O₂ atmospheres at E9.5-E10.5 or E10.5-E11.5 and analyzed by LC-MS. Heatmap displays 56 metabolites evaluated normalized to total ion content. All data represent a minimum of 3 independent experiments, except TEMI (n = 2 ex utero, 2 in utero).

Moreover, to determine whether the E10.5-E11.5 metabolic switch represents a coordinated embryo-wide process or tissue-restricted metabolic remodeling, we dissected internal organs (including lungs, intestine, liver, gonads, pancreas), heart and brain from ex utero and in utero embryos at these respective stages and subjected each compartment to LC-MS profiling **(Extended data Fig. 13C)**. This organ-resolved metabolomics dataset revealed a pronounced metabolic transition in the internal organs at E11.5, whereas the embryonic heart and brain displayed comparatively more modest but evident shifts in global metabolic profiles, with stage-dependent changes more evident in utero **(Fig. 5B)**. To verify these results within specific cell types rather than within heterogeneous tissues, we leveraged our single-cell RNA-sequencing dataset, which previously identified a transcriptional correlate of the metabolic switch at the whole-embryo level. Strikingly, KEGG pathway mapping across annotated cell types present from E10.5 to E12.5 demonstrated a synchronous metabolic reprogramming in all examined cell types (including neural progenitors, chondrocytes and osteoblasts, epithelial cells, erythroid lineage, hepatocytes, myocytes, white blood cells, cholinergic neurons, and Schwann cell precursors) both ex utero and in utero **(Fig. 5C, Extended data Fig. 14B)**. These results corroborate that the metabolic transition at E10.5-E11.5 represents a global synchronized shift in embryonic metabolic state coordinated by transcriptional changes across most embryonic tissues rather than localized lineage-restricted changes.

The use of embryo ex utero culture systems allows for applying environmental perturbations in live developing embryos. Thus, we tested whether altered oxygen concentrations can accelerate or abolish the midgestation metabolic transition. Given that and increase in oxygen due to placental function at E10.5 was hypothesized to control this metabolic switch, we cultured embryos under different oxygen concentrations from E9.5 to E11.5 and profiled their metabolome by LC-MS and [U-¹³C]glucose tracing **(Fig. 5D-F)**. Morphologically, only embryos cultured in 80% oxygen seemed viable and developed properly to E11.5, although overall development progressed also in lower oxygen conditions **(Fig. 5D)**. Glucose tracing revealed that only embryos grown in a 21% oxygen atmosphere showed significantly lower levels of glucose carbon enrichment in TCA cycle intermediates (**Fig. 5E**), with higher oxygen percentages leading to enrichment in citrate m+4 indicating multiple turns of the TCA cycle and more robust mitochondrial metabolism **(Extended data Fig. 15A)**. These changes in mitochondrial metabolism were coupled to decreased glucose carbon enrichment in purine metabolites (**Extended data Fig. 15B**) suggesting that increased purine synthesis during the transition is linked to oxygen availability. The metabolic profile of embryos cultured in 40% oxygen from E9.5 to E10.5 showed no changes compared to those grown under 21% oxygen atmosphere, indicating that higher oxygen does not trigger an earlier metabolic profile switch **(Fig. 5F)**. Notably, only embryos cultured under 80% oxygen displayed the full metabolomic signature characteristic of the E11.5 metabolic switch, whereas embryos maintained under lower oxygen tension showed only a partial transition in metabolic profile **(Fig. 5F)**. Altogether, these data demonstrate that ex utero embryos establish spatially organized metabolite regions in the absence of maternal and placental environments and undergo a conserved, tissue-wide midgestational metabolic transition. Although this transition is partially modulated by oxygen delivery, it cannot be prematurely induced by increased oxygen availability, indicating that it is developmentally timed rather than triggered by oxygen.

### Characterization of glucose metabolism and redox metabolic switches during ex utero embryogenesis

Mammalian midgestation is characterized by rapid embryonic and placental growth, increased oxygen availability, and a concomitant rise in the contribution of maternally derived glucose to the embryonic tricarboxylic acid (TCA) cycle^2^. To determine whether the E10.5-E11.5 increase in glucose oxidation is recapitulated in embryos cultured ex utero, we leveraged [U-¹³C]glucose isotope tracing. First, we evaluated glucose utilization and mitochondrial metabolic state during ex utero development by ¹³C-glucose isotope tracing across mid-gestation. Consistent with previous results^4^, labelling of TCA-cycle intermediates in utero increased progressively from E10.5 (**Extended data Fig. 17A-D**). Remarkably, embryos cultured ex utero from E6.5 or E7.5 also exhibited a progressive increase in ¹³C incorporation as culture progressed, although this rise emerged starting at approximately E9.5 (**Fig. 6A, Extended data Fig. 17**), suggesting that the shift toward glucose oxidation may occur earlier in culture than in utero. These results suggest that embryos developing ex utero transit to oxidative phosphorylation-based metabolism during organogenesis, but they do so earlier than in utero conditions, likely due to higher oxygen levels in culture than in vivo.

**Figure 6.**
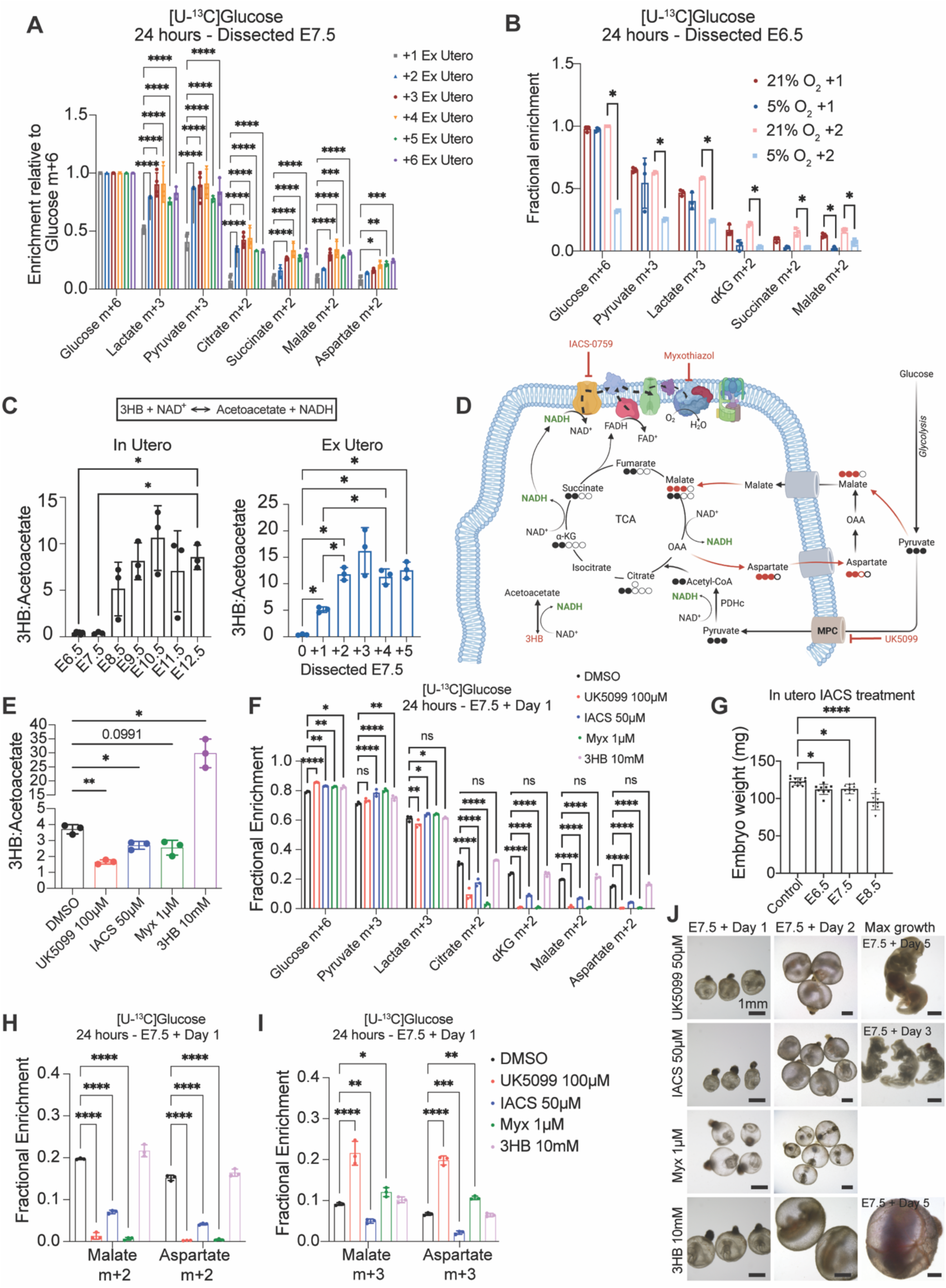
Isotope tracing characterization of glucose metabolism and redox metabolic switches in ex utero-grown embryos. **A,** Incorporation of [U-^13^C]-glucose-derived carbon into TCA-cycle intermediates across developmental time in embryos grown ex utero from E6.5. Values are normalized to Glucose m+6. **B,** [U-^13^C]-Glucose tracing for 24 hours on embryos cultured under 5% or 21% O₂ during gastrulation (E6.5-E8.5). Tracing was started at E6.5 (+1) or at E7.5 (+2). Values are normalized to Glucose m+6. **C,** Increase in the ratio of 3-hydroxybutyrate to acetoacetate, indicator of the mitochondrial NAD⁺/NADH redox state, in both in utero and ex utero embryos at E8.5. **D,** Schematic representation of the electron transport chain in the mitochondria, displaying the specific inhibition of MPC, Complex I and Complex II by small molecules inhibitors, and the role of 3HB in NAD^+^/NADH balance. **E**, Decreased 3HB:acetoacetate ratio in ex utero cultured embryos treated at E7.5 with the MPC inhibitor UK5099, mitochondrial Complex I inhibitor IACS-10759, or the Complex III inhibitor Myxothiazol after 24 hours. In contrast, 3HB increases 3HB:acetoacetate ratio. **F,** [U-^13^C]-glucose tracing in embryos exposed to small molecule inhibitors showing a decrease in TCA-cycle intermediates after exposure to UK5099, IACS-10759, and Myxothiazol, while 3HB does not cause a significant change compared to vehicle controls (DMSO). **G,** Changes in embryo weight measured at E12.5 after in utero treatment with IACS at E6.5, E7.5 and E8.5. **H-I**, Compensatory malate-aspartate shuttle metabolism in ETC inhibitor-treated embryos. Perturbing redox balance in gastrulating embryos showed reduced malate/aspartate m+2 enrichment (**H**), in parallel to increased malate m+3 and aspartate m+3 (**I**). **J,** Bright field images of embryos treated with UK5099, IACS, Myxothiazol and 3HB from E7.5 to E8.5 and kept in culture for up to 5 days or until the maximum developmental growth. All data represent mean ± S.D. of a minimum of 3 biological replicates. * = p<0.05, ** = p<0.01; *** = p<0.001; **** = p<0.0001.

To investigate metabolic regulation at cellular resolution, we inferred pathway-level metabolic activity from single-cell transcriptomic data using the flux-balance framework COMPASS^64^ (**Extended data Fig. 18A**). Embryos cultured ex utero to E11.5-E12.5 exhibited globally increased inferred metabolic activity relative to in utero controls, including in central carbon metabolism including amino acid pathways, the TCA cycle, glycolysis, and oxidative phosphorylation (**Extended data Fig. 18B-E**). Cell-type–resolved analysis revealed a balanced response in embryos explanted at E7.5, with some lineages showing increased and others reduced activity (**Extended data Fig. 19**). In contrast, embryos cultured from E6.5 displayed elevated metabolic activity across nearly all cell types (**Extended data Fig. 20**), indicating that earlier removal from the uterine environment may alter embryonic metabolism more profoundly. Increased metabolic activity was already evident during gastrulation (E6.5-E8.5) ex utero (**Extended data Fig. 21**). Gene ontology analysis independently confirmed enrichment of metabolic processes, including pyruvate metabolism, carbohydrate catabolism, and hypoxia response pathways (**Extended data Fig. 22**). Overall, these findings indicate that while ex utero embryos retain the core metabolic programs characteristic of in utero development, early culture, particularly from pre-gastrulation stages, induces a context-dependent upregulation of metabolic gene expression, probably reflecting adaptation to altered oxygen and nutrient availability.

Given the dynamic changes in oxygen availability that accompany early embryogenesis both in utero and in culture, we next harnessed the ex utero platform to interrogate and perturb mitochondrial metabolic transitions during gastrulation. Oxygen concentrations range from 1% to 5% (pO_2_ 0.5–30 mmHg) in the uterine environment^65^, and low oxygen tension has been shown to support proper mouse embryo development in rotating culture systems as well as in vitro embryo models^4,22,66–68^. In contrast, embryos cultured ex utero from gastrulation under static hypoxic conditions fail to develop beyond two days^4,5^, requiring atmospheric oxygen levels (21%) to transit through gastrulation and organogenesis. To directly assess how oxygen tension shapes metabolic activity during this window, we performed ¹³C-glucose tracing from E6.5 to E8.5 under either 5% or 21% O₂. During the first 24 hours of culture, embryos maintained at 5% O₂ exhibited similar glucose oxidation to those cultured at 21% O₂, as indicated by higher incorporation of glucose-derived carbon into glycolysis and TCA-cycle intermediates (**Fig. 6B**). By E8.5, however, embryos exposed to hypoxia displayed a pronounced reduction in ¹³C-glucose uptake and labeling across all measured metabolic intermediates, including both oxidative and non-oxidative pathways (**Fig. 6B**). These findings may provide a metabolic explanation for the failure of embryos to sustain growth beyond early gastrulation under in vitro hypoxic conditions and might indicate a critical oxygen-sensitive metabolic shift during this interval.

The activity of the TCA cycle is linked to the production of mitochondrial NADH that fuels the electron transport chain (ETC). To assess mitochondrial redox dynamics during early organogenesis, we quantified the 3-hydroxybutyrate (3HB)/acetoacetate ratio, as a proxy for the mitochondrial NAD^+^/NADH balance. We observed a significant increase in this ratio from E7.5-E8.5 in both in utero and ex utero embryos driven by an increase in 3HB levels across gestation (**Fig. 6C**). This increased 3HB production implies an increase in NADH production in the mitochondria during this transition. To test the functional relevance of this transition and the capacity for metabolic flexibility during gastrulation, we applied a panel of small-molecule inhibitors targeting different components of mitochondrial metabolism (**Fig. 6D**) previously implicated in stem cell differentiation and development^69–74^. Blocking mitochondrial pyruvate import with the MPC inhibitor UK5099, inhibiting Complex I with IACS-010759 to limit NADH to NAD^+^ conversion, or disrupting ETC Complex III with Myxothiazol each reduced the 3HB/acetoacetate ratio, suppressed ^13^C-glucose incorporation into TCA-cycle intermediates and lead to evident morphological aberrations after 24 hours of exposure at E7.5-E8.5 (**Fig. 6E-J**). Conversely, 3HB supplementation further increased NADH abundance, as indicated by an increase in 3HB:Acetoacetate, had no impact of ^13^C glucose contribution to TCA intermediates, and generated no evident morphological defects in gastrulating embryos (**Fig. 6E, F, J**). We validated the effect of perturbing the NADH redox shift by treating pregnant dams with 10 mg/kg IACS at E6.5-E8.5 in utero, analyzing the effect on developing embryos at E12.5. We observed a significant reduction of embryo weight, especially when treatment was done at E8.5 (**Fig. 6G**). Interestingly, embryos treated with UK5099 or Myxothiazol showed evidence of malic enzyme activity and altered malate-aspartate shuttle metabolism, evidenced by an increased malate m+3 and aspartate m+3 (**Figure 6H, I**) and consistent with pyruvate carboxylase activity, suggesting a compensatory attempt to increase NADH in the mitochondria using alternative metabolic pathways. Despite their metabolic flexibility, embryos failed to restore mitochondrial redox balance and embryonic development failed to progress properly when NADH balance was perturbed, even after removal of the inhibitors at E8.5 followed by ex utero culture (**Figure 6J**). These findings reveal that successful transition from gastrulation to early organogenesis requires the ability to sustain appropriate mitochondrial NADH levels and TCA-cycle flux.

## Discussion

In this work, we established robust and reproducible embryo ex utero culture systems, capable of supporting mouse embryogenesis from gastrulation to the end of organogenesis, thus extending the limits of maternal-independent embryo growth, and allowing direct experimental access to the mammalian embryo through the whole window of mammalian gastrulation and organogenesis. By combining optimized gas regulation systems capable of maintaining a hypoxic-hyperoxic environment, and dynamic nutrient supplementation, our culture platform supports proper morphogenesis of all major tissues, while preserving transcriptional and metabolic programs that closely match in utero development independently of maternal inputs. The ability to support mouse embryogenesis ex utero for long-term provides an unprecedented opportunity to mechanistically dissect, perturb, and ultimately engineer, embryonic development in mammals. We further demonstrated the amenability of the ex utero technologies described herein for interrogation of embryonic metabolism.

Our whole-embryo ex utero metabolomic and transcriptomic profiling revealed that cultured embryos recapitulate the characteristic E10.5-E11.5 metabolic transition previously described in vivo, including coordinated shifts in purine metabolism, amino acid flux, TCA cycle activity, and mitochondrial redox balance. The preservation of this transition ex utero in most embryonic tissues revealed that it represents a body-wide intrinsic developmental program, regulated by cell-autonomous or embryo-intrinsic mechanisms rather than by placental signaling or maternal nutrient supply. Besides, differences between serum sources further demonstrate the effects of nutrient composition shaping embryonic metabolic landscapes. Our ability to modify environmental components in live developing embryos ex utero revealed that some features of the midgestation metabolic shift are sensitive to extrinsic conditions, whereas others are highly robust. For example, high oxygen culture conditions gradually activate the TCA cycle one day earlier than in vivo, however, early increase in oxygen availability at E9.5 did not trigger most other hallmarks of the metabolic transition, indicating that certain metabolic modules during organogenesis remain plastic while others appear resistant to extrinsic perturbation. Notably, modulation of oxygen availability during the E10.5-E11.5 window indicated that increased oxygen levels are necessary for the metabolic transition observed at this stage. Embryos cultured under reduced oxygen concentrations failed to fully undergo the characteristic metabolic switch, despite progressing with minimal morphological abnormalities until ∼E11. Because sustained development beyond this stage is incompatible with low oxygen tension, it remains unclear whether the attenuated metabolic transition reflects a direct effect of oxygen availability or a secondary consequence of impaired developmental progression. These observations suggest that the midgestation metabolic transition is indispensable for proper embryo development beyond E10.5, although the developmental basis for the timing and nature of this shift remain to be elucidated. Importantly, we identified an early redox shift at the end of gastrulation (E7.5-E8.5) both in utero and ex utero, characterized by a rise in the 3-hydroxybutyrate/acetoacetate ratio, thus reflecting increased mitochondrial NADH production, suggesting oxidative metabolism is activated in the embryo upon gastrulation. Pharmacological inhibition of this redox transition revealed that perturbing it halts developmental progression, underscoring the importance of mitochondrial dynamics as a regulator of embryogenesis. Collectively, these findings illustrate the amenability of ex utero culture to enable mechanistic dissection of embryo metabolism with a precision not achievable in vivo, where embryo metabolism is inseparably entangled with maternal physiology.

Together, these results establish ex utero culture as a powerful experimental system for dissecting the regulatory logic of mammalian embryogenesis and reveal a greater degree of developmental autonomy during mammalian embryogenesis than previously considered. The integration of stem cell-derived embryo models, synthetic biology and ex utero culture systems promises to reshape our understanding of mammalian development and expand the therapeutic possibilities of these technologies.

## Acknowledgments

This work was funded by the Howard Hughes Medical Institute, University of Texas Southwestern, Pascal and Ilana Mantoux, the Helen and Martin Kimmel Institute for Stem Cell Research, the Flight Attendant Medical Research Institute (FAMRI), the Institute for Artificial Intelligence in Weizmann Institute, the MBZUAI-WIS Program for Collaborative Research in AI, the Wolfson Foundation, ICRF-Research Professorship, Institute for Environmental Sustainability (IES) at the Weizmann Institute, the Center for Research on the Development of Innovative Neurotechnologies, the Belle S. and Irving Meller Center for Biology of Aging, Dr. Barry Sherman Institute for Medicinal Chemistry, the Zantker Charitable Foundation, the Estate of Zvia Zeroni, ERC-COG-2022 #101089297—ExUteroEmbryogenesis, Israel Science Foundation-Breakthrough Research Grant (1098/25), the 2025 Nakasone Award by the Human Frontiers Science Program (HFSPO), Minerva Stiftung. The authors thank vivarium staff Brooke Groff and Crystall Lopez from the Howard Hughes Medical Institute Janelia Research Campus for mouse colony care and timed-pregnant setups. The authors acknowledge the Howard Hughes Medical Institute at Janelia Research Campus Molecular Genomics Shared Resource Core Facility (RRID:SCR_026832), and the Quantitative Genomics Core Facility (RRID:SCR_022694).

## Author contributions

D.L. established the ex utero culture protocol, adapted gas regulation systems to the rotating culture, performed embryology, immunostainings, imaging, toxicology assays, and assisted on metabolomic experiments. S.G. performed all single cell transcriptomics bioinformatic analyses and assisted on manuscript writing. W.X. performed metabolomic LCMS analysis and in utero assays. A.K. assisted on assemble, design and measurements of electronic gas regulator modules coupled to embryo incubators. F.R. assisted on embryo culture and immunostainings. W.A., D.N., and N.K. assisted on metabolomic LCMS analysis. A.W. engineered optimized embryo culture incubators. C.C. performed SmartSPIM light-sheet microscopy. S.G. assisted on toxicology assays. L.D., A.W., P.T., M.W., H.Z., T.D., and L.L. performed and supervised sample preparation, processing and analysis for mass spectrometry imaging. M.C. performed cryosectioning. I.C. and S.S. performed sci-seq3 library preparation and sequencing. R.S. assembled and designed Arad Technologies gas regulators. E.G.C assisted on immunostainings and manuscript writing. B.O., A.Y., G.G., M.Y-C. and S.V. collected human adult blood serum and assisted in culture condition testing. L.W., A.Y., K.M., C.R., and K.D. performed library preparation and sequencing for spatial transcriptomics. S.W. and M.E.F-R. recruited donors and performed human cord blood serum extraction during cesarean sections at UTSW. A.H-C., A.V., and G.L. reproduced ex utero culture techniques in their independent laboratory. N.N. assisted on bioinformatic analysis. I.M., N.K. and R.R.K., collected cord blood and calibrated human cord serum production. I.E-M. assisted on embryo clearing methods. A.S. conceived the idea for this project, performed and supervised execution of LC-MS metabolomics and glucose tracing experiments, adequate data analysis, and wrote the manuscript. J.H.H. conceived the idea for this project, supervised executions of experiments, adequate analysis of data and wrote the manuscript. A.A-C. conceived the idea for this project, established the ex utero culture protocol and gas regulation systems/incubators, designed and conducted most embryology, sequencing, imaging and metabolism experiments, supervised executions of experiments, adequate data analysis, and wrote the manuscript.

