## Supplementary figures for "Reconstituting Mouse Embryogenesis Ex Utero from Gastrulation to Fetal Development Reveals Maternally Independent Metabolic Programs"

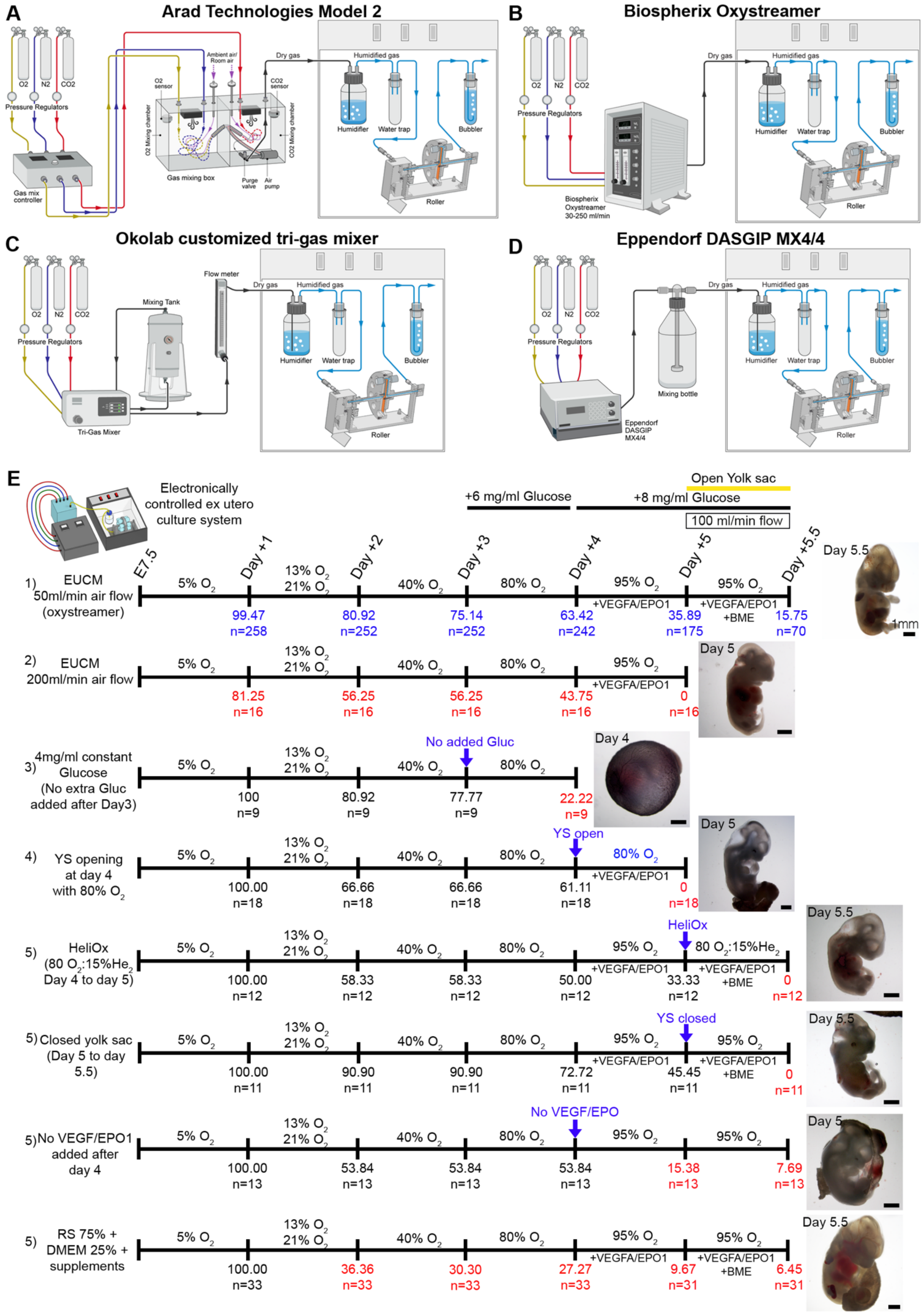
A
Arad Technologies Model 2
B
Biospherix Oxystreamer
C
Okolab customized tri-gas mixer
D
Eppendorf DASGIP MX4/4
O2
N2
CO2
Pressure Regulators
Gas mix controller
Gas mixing box
Dry gas
Humidified gas
Humidifier
Water trap
Bubbler
Roller
Biospherix Oxystreamer 30-250 ml/min
Mixing Tank
Flow meter
Tri-Gas Mixer
Mixing bottle
Eppendorf DASGIP MX4/4
E
Electronically controlled ex utero culture system
+6 mg/ml Glucose
+8 mg/ml Glucose
Open Yolk sac
100 ml/min flow
E7.5
Day +1
Day +2
Day +3
Day +4
Day +5
Day +5.5
1) EUCM 50ml/min air flow (oxystreamer)
5% O2
13% O2 21% O2
40% O2
80% O2
95% O2
+VEGFA/EPO1
+BME
99.47 n=258
80.92 n=252
75.14 n=252
63.42 n=242
35.89 n=175
15.75 n=70
Day 5.5
1mm
2) EUCM 200ml/min air flow
81.25 n=16
56.25 n=16
56.25 n=16
43.75 n=16
0 n=16
Day 5
3) 4mg/ml constant Glucose (No extra Gluc added after Day3)
No added Gluc
100 n=9
80.92 n=9
77.77 n=9
22.22 n=9
Day 4
4) YS opening at day 4 with 80% O2
YS open
80% O2
100.00 n=18
66.66 n=18
66.66 n=18
61.11 n=18
0 n=18
Day 5
5) HeliOx (80 O2:15%He2 Day 4 to day 5)
HeliOx
80 O2:15%He2
100.00 n=12
58.33 n=12
58.33 n=12
50.00 n=12
33.33 n=12
0 n=12
Day 5.5
5) Closed yolk sac (Day 5 to day 5.5)
YS closed
100.00 n=11
90.90 n=11
90.90 n=11
72.72 n=11
45.45 n=11
0 n=11
Day 5.5
5) No VEGF/EPO1 added after day 4
No VEGF/EPO
100.00 n=13
53.84 n=13
53.84 n=13
53.84 n=13
15.38 n=13
7.69 n=13
Day 5
5) RS 75% + DMEM 25% + supplements
100.00 n=33
36.36 n=33
30.30 n=33
27.27 n=33
9.67 n=31
6.45 n=31
Day 5.5

**Extended data Fig. 1. Establishment and optimization of electronic gas regulation modules adapted for embryo culture in rotating bottles. A-D,** Schematic diagrams displaying the designed configuration of the four gas regulation systems coupled to the roller culture incubator: **A**, Arad Technologies Model 2; **B**, Biospherix OxyStreamer; **C**, Okolab customized tri-gas mixer; **D**, Eppendorf DASGIP MX4/4. In each configuration, oxygen, nitrogen, and carbon dioxide are mixed and regulated to achieve precise gas composition and flow or pressure control. Gases are then delivered into a humidification bottle within a 37 °C incubator. Humidified gas flows into a water trap and subsequently enters the hollowed rotating drum. Culture bottles are directly plugged onto the drum, which distributes humidified gas into each individual bottle. Excess gas exits the hollowed drum through an outlet connected to a water bubbler that provides a visual indicator of proper flow. **E**, Percentage of normally developed embryos under distinct conditions, including variations in gas flow rate, glucose concentration, oxygen percentage, media supplementation, etc. Values in red indicate the conditions affecting efficiency of embryo survival. “n” denotes the number of embryos assessed per condition at each timepoint. Embryos dissected, fixed or transferred to alternative conditions were subtracted from subsequent totals. Representative bright field images corresponding to selected conditions are shown (right panels).

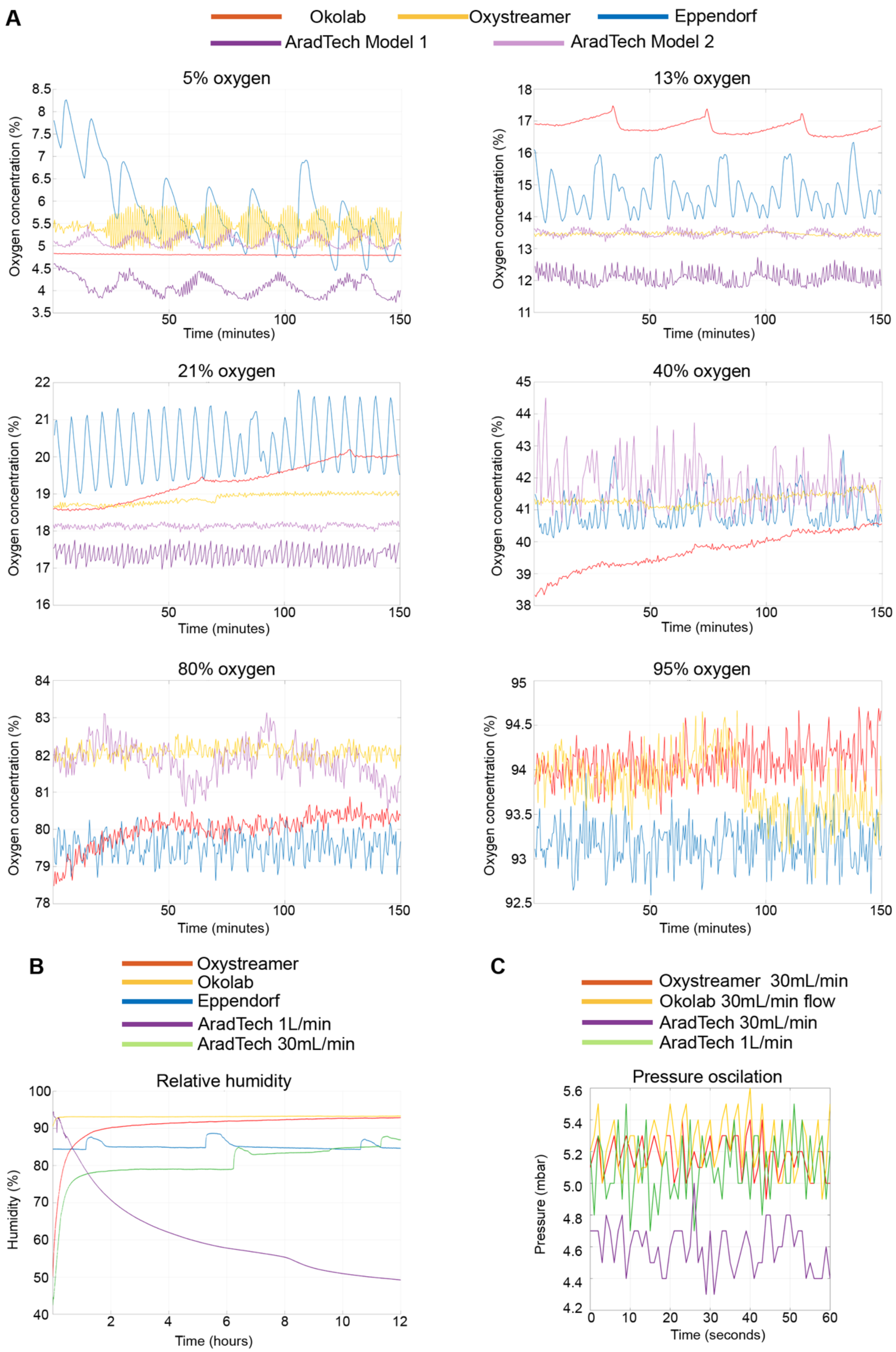
A
Okolab
Oxystreamer
Eppendorf
AradTech Model 1
AradTech Model 2
5% oxygen
13% oxygen
21% oxygen
40% oxygen
80% oxygen
95% oxygen
Oxygen concentration (%)
Time (minutes)
B
Oxystreamer
Okolab
Eppendorf
AradTech 1L/min
AradTech 30mL/min
Relative humidity
Humidity (%)
Time (hours)
C
Oxystreamer 30mL/min
Okolab 30mL/min flow
AradTech 30mL/min
AradTech 1L/min
Pressure oscilation
Pressure (mbar)
Time (seconds)

31

**Extended data Fig. 2. Measurement of oxygen stability, humidity levels, and internal drum pressure across electronic gas-regulation systems. A**, Oxygen concentrations measured on each electronic gas-regulation module when set to 5%, 13%, 21%, 40%, 80%, or 95% $O_2$ over a 150-minute interval. The values for each gas regulation module are represented in colored lines, as indicated in the legend. Colored lines represent real-time oxygen readings for the Arad Technologies Model 1, Arad Technologies Model 2, Biospherix OxyStreamer, Okolab tri-gas mixer, and Eppendorf DASGIP systems, as indicated. **B**, Relative humidity values measured for each system over a 12-hour period. The Arad Technologies unit was assessed at both 1 L/min and 30 mL/min flow rates; all other systems were measured at 50 mL/min. **C**, Differential pressure oscillation values (mbar per second) recorded inside the rotating drum for each gas-regulation system. Arad Technologies Model 2 system was evaluated at two flow rates (1 L/min and 30 mL/min), whereas other modules were tested at 50 mL/min.

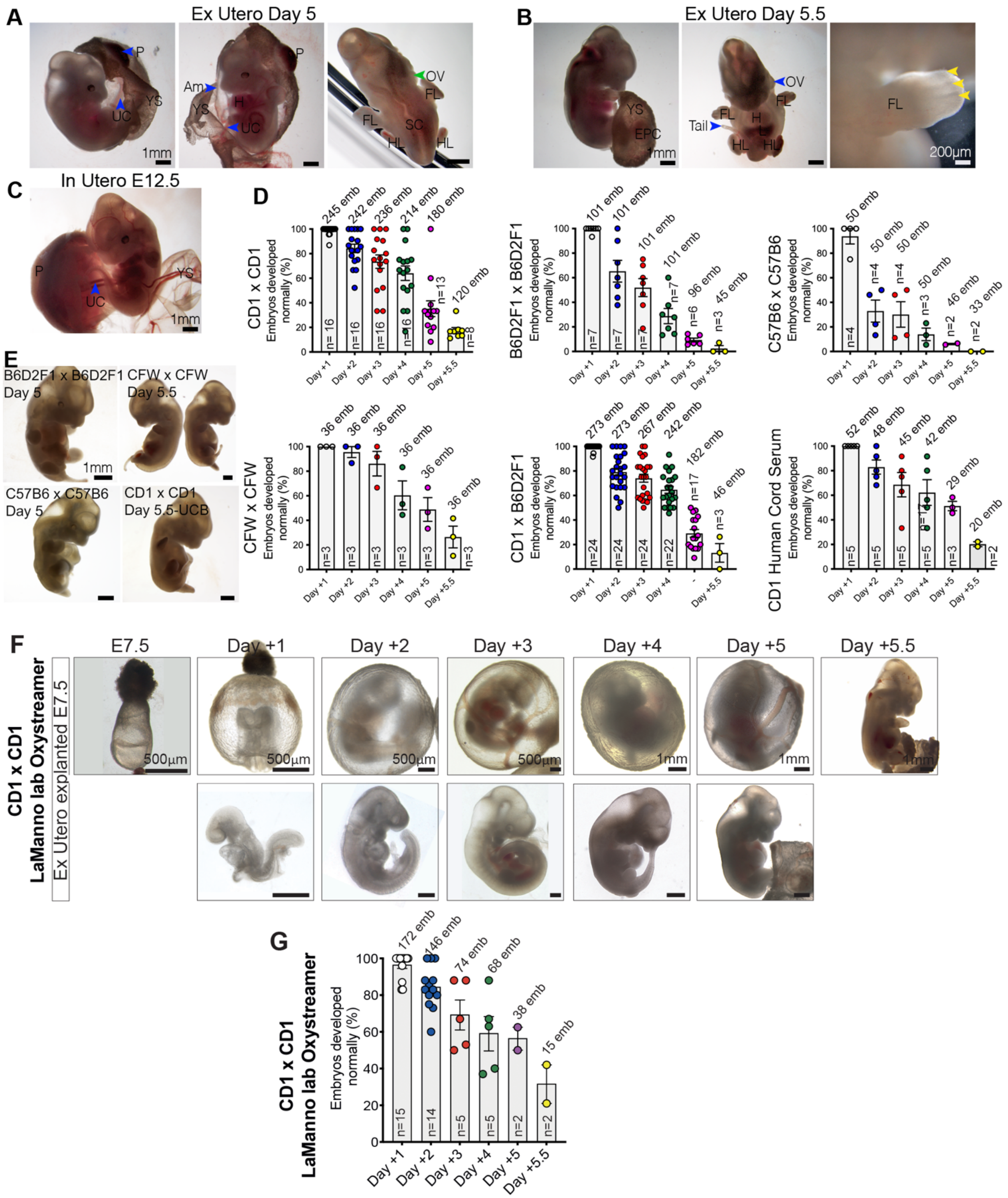


**Extended data Fig. 3. Ex utero development across mouse strains. A**, Representative bright-field images of CD1 embryos cultured ex utero from E7.5 to day +5 and dissected out of the yolk sac, preserving the connection of the umbilical cord to the placenta. **B**, Images of CD1 embryos cultured ex utero for 5.5 days (exteriorized from the yolk sac) starting at E7.5. Right panel is a high-magnification image of the same embryo displaying digit primordia in the forelimb (yellow arrows). **C**, In utero CD1 embryo at E12.5 dissected out of the yolk sac while preserving the connection of the umbilical cord to the placenta, shown for comparison. **D**, Quantification of normally developed embryos across multiple mouse strains and human serum supplementation. CD1×CD1, B6D2F1×B6D2F1, CFW×CFW, and CD1×B6D2F1 were cultured with human adult serum (HAS). C57BL/6×C57BL/6 was supplemented with umbilical cord blood serum (HCS). Human umbilical cord blood serum (HCS) was also tested as a supplement in CD1×CD1 embryos without evident differences observed compared to HAS. **E,** Representative day +5 or day +5.5 embryos from each strain combination shown in (**D**), illustrating the range of morphological outcomes and strain-dependent differences in ex utero growth. **F**, Bright-field images showing the developmental progression of CD1 embryos cultured in an independent laboratory at EPFL using the OxyStreamer system, from E7.5 through Day +5.5. For each timepoint, embryos are shown within the yolk sac (top panels) and after yolk sac dissection (bottom panels). **G**, Quantification of normally developed CD1×CD1 embryos grown using the OxyStreamer platform in an independent laboratory at EPFL. “emb” denotes the total number of embryos evaluated; “n” indicates the number of independent experiments. Data represent mean ± s.e.m. Scale bars are indicated on each image. Abbreviations: Am, amnion; FL, forelimb; H, heart; HL; hindlimb; UC, umbilical cord; OV, otic vesicle; P, placenta; SC, spinal cord; YS, yolk sac.

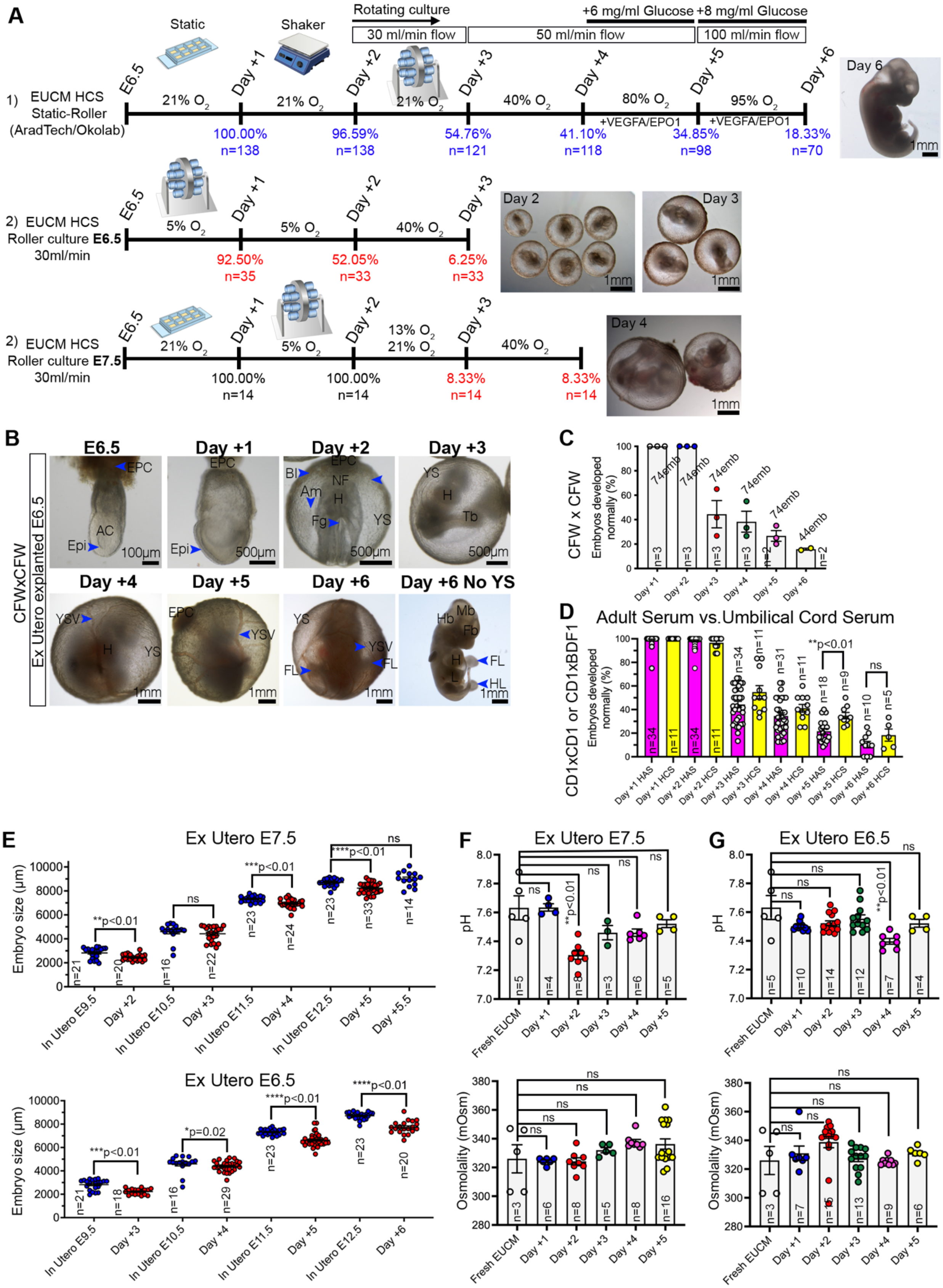
A
Static
Shaker
Rotating culture
30 ml/min flow
50 ml/min flow
100 ml/min flow
+6 mg/ml Glucose
+8 mg/ml Glucose
1) EUCM HCS Static-Roller (AradTech/Okolab)
E6.5
Day +1
Day +2
Day +3
Day +4
Day +5
Day +6
21% $O_2$
21% $O_2$
21% $O_2$
40% $O_2$
80% $O_2$
95% $O_2$
+VEGFA/EPO1
+VEGFA/EPO1
100.00% n=138
96.59% n=138
54.76% n=121
41.10% n=118
34.85% n=98
18.33% n=70
Day 6
1mm
2) EUCM HCS Roller culture E6.5 30ml/min
5% $O_2$
5% $O_2$
40% $O_2$
92.50% n=35
52.05% n=33
6.25% n=33
Day 2
Day 3
2) EUCM HCS Roller culture E7.5 30ml/min
21% $O_2$
5% $O_2$
13% $O_2$
21% $O_2$
40% $O_2$
100.00% n=14
100.00% n=14
8.33% n=14
8.33% n=14
Day 4
B
CFWxCFW Ex Utero explanted E6.5
E6.5
Day +1
Day +2
Day +3
Day +4
Day +5
Day +6
Day +6 No YS
EPC
AC
Epi
BI
Am
NF
H
Fg
YS
Tb
YSV
FL
Hb
Mb
Fb
L
HL
100μm
500μm
1mm
C
CFW x CFW
Embryos developed normally (%)
74emb
44emb
D
Adult Serum vs.Umbilical Cord Serum
CD1xCD1 or CD1xBDF1
**p<0.01
ns
Day +1 HAS
Day +1 HCS
Day +2 HAS
Day +2 HCS
Day +3 HAS
Day +3 HCS
Day +4 HAS
Day +4 HCS
Day +5 HAS
Day +5 HCS
Day +6 HAS
Day +6 HCS
E
Ex Utero E7.5
Embryo size (μm)
**p<0.01
***p<0.01
****p<0.01
In Utero E9.5
In Utero E10.5
In Utero E11.5
In Utero E12.5
Day +5.5
Ex Utero E6.5
***p<0.01
*p=0.02
****p<0.01
****p<0.01
F
Ex Utero E7.5
pH
**p<0.01
Fresh EUCM
Osmolality (mOsm)
G
Ex Utero E6.5

**Extended data Fig. 4. Extending and characterizing mouse embryo ex utero culture systems.** **A,** Schematic protocol indicating the percentage of E6.5 embryos developed properly per day when transferred to the rotating culture at E6.5, E7.5 and E8.5. The media composition, static or roller culture, and oxygen concentrations are specified for each protocol. Representative bright field images of embryos cultured are shown on the right side of the respective protocol. "n" denotes the number of embryos evaluated per condition. Embryos dissected, fixed or transferred to other conditions are subtracted from the total. Numbers in blue indicate the protocol yielding the highest efficiency of embryo survival that was subsequently used throughout the study. Values in red indicate the conditions affecting efficiency of embryo survival. **B**, Representative images of CFW×CFW embryos cultured ex utero from E6.5 through six days of development. Removal of the yolk sac (day +6, right) highlights morphology of the embryo proper. **C,** Quantification of properly developed CFW×CFW embryos across days of culture. "emb" denotes total number of embryos evaluated; "n" reflects number of independent experiments. **D,** Comparison of properly developed embryos when grown in EUCM supplemented with either in adult human serum or human umbilical cord serum (CD1×CD1 or CD1×B6D2F1 embryos). Adult serum supports early development but shows significantly reduced performance at later stages, whereas cord serum maintains higher developmental efficiency. Dots represent independent experiments, with "n" values corresponding to the number of experiments assessed. **E,** Embryo size measurements for ex utero embryos cultured at E7.5 (top) or E6.5 (bottom), compared with in utero controls collected at equivalent developmental stages. Length of the antero-posterior axis was measured for E6.5 to E8.5 and the crown-rump length was measured for later stages. Ex utero embryos closely match in utero size until late stages, with deviations quantified by statistical analysis. Dots represent individual embryos. "n" values correspond to individual embryos measured. **F-G,** Measurements of pH (top) and osmolarity (bottom) are shown for embryos cultured from E7.5 (**F**) or E6.5 (**G**) through 5 days. Values of fresh EUCM are shown for comparison. Statistical comparisons shown where relevant. Data are mean ± s.e.m. P values are indicated. Ns, not significant. Abbreviations: AC, amniotic cavity; Am, Amnion; BI, Blood islands; Epi, Epiblast; EPC, ectoplacental cone; Fg, foregut pocket; FL, forelimb; H, heart; HL; hindlimb; L, Liver; NF, Neural folds; Tb, Tail bud; YS, yolk sac; YSV, yolk sac vessel.

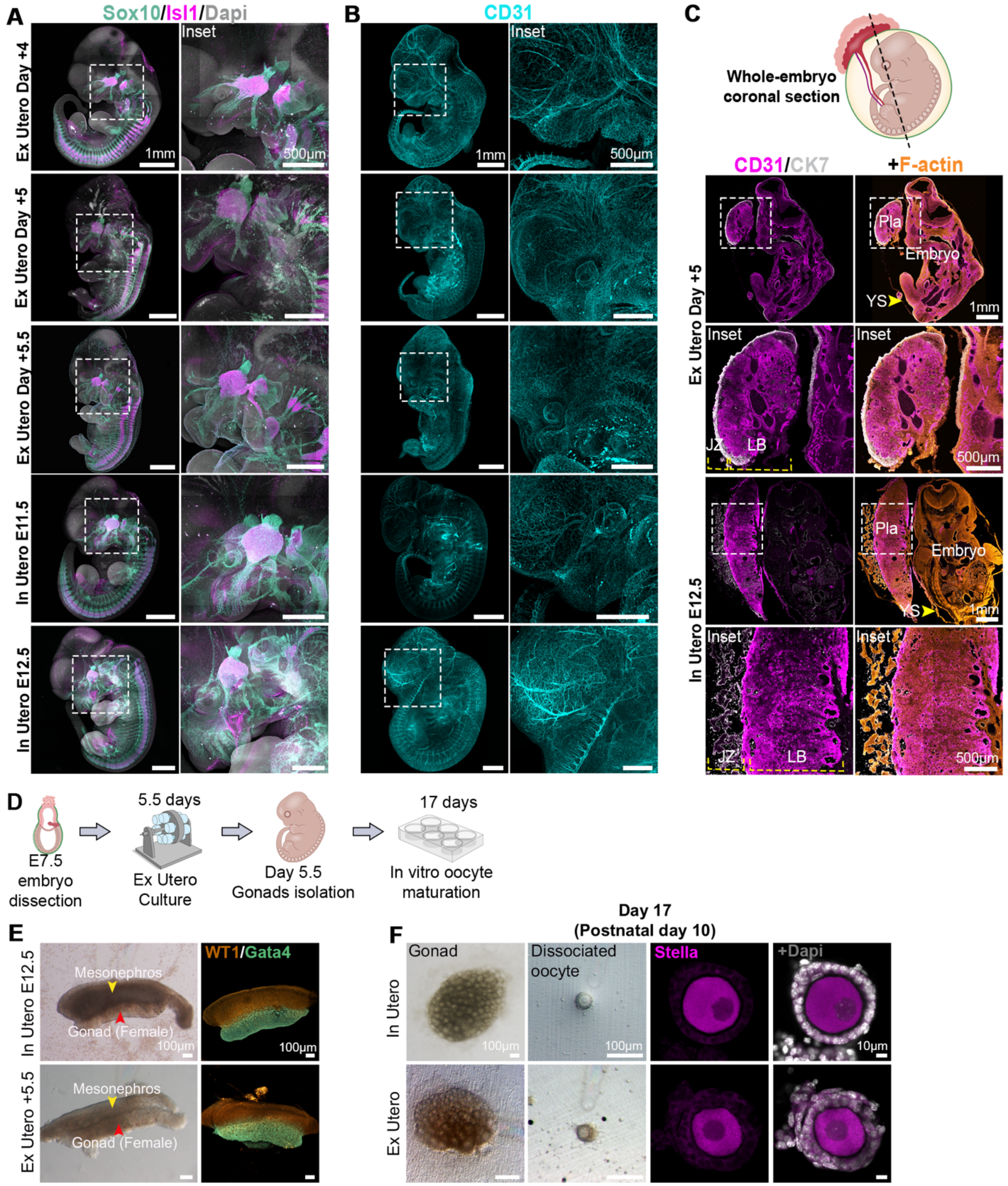


**Extended data Fig. 5. Spatiotemporal expression of lineage-specific markers and extraembryonic tissues in ex utero-grown embryos. A,** Whole-mount immunostaining images of embryos developed ex utero (day +4 to day +5.5) and time-matched in utero controls (E11.5–E12.5) stained for SOX10 and ISL1. Insets show magnified views of the boxed regions. **B**, Maximum intensity projections of ex utero and in utero embryos labeled for CD31 to identify the embryonic vasculature. Insets are enlargements of the dashed boxes. **C**, Representative sagittal mid-section immunohistochemistry of whole embryos with associated yolk sac (YS) and placental (Pla) tissues. Sections are stained for CD31 (magenta), cytokeratin 7 (CK7; yellow), and F-actin (phalloidin; orange) at the indicated stages. **D**. Schematic overview of the in vitro oocyte maturation protocol from ex utero embryos. Embryos dissected at E7.5 were cultured ex utero for 5.5 days, after which gonads were isolated and subjected to a 17-day in vitro maturation protocol. **E**, Representative bright-field and immunostaining images of female gonads from in utero and ex utero embryos. WT1 and GATA4 staining demarcate the mesonephros and embryonic gonad. **F**, Isolated gonads were grown in vitro to day 17, dissociated into individual follicles, and immunostained for the canonical oocyte marker Stella. White, DAPI. Images represent a minimum of 3 biological replicates. Scale bars are indicated on each panel. Abbreviations: JZ, junctional zone; LB, labyrinth; Pla, placenta.

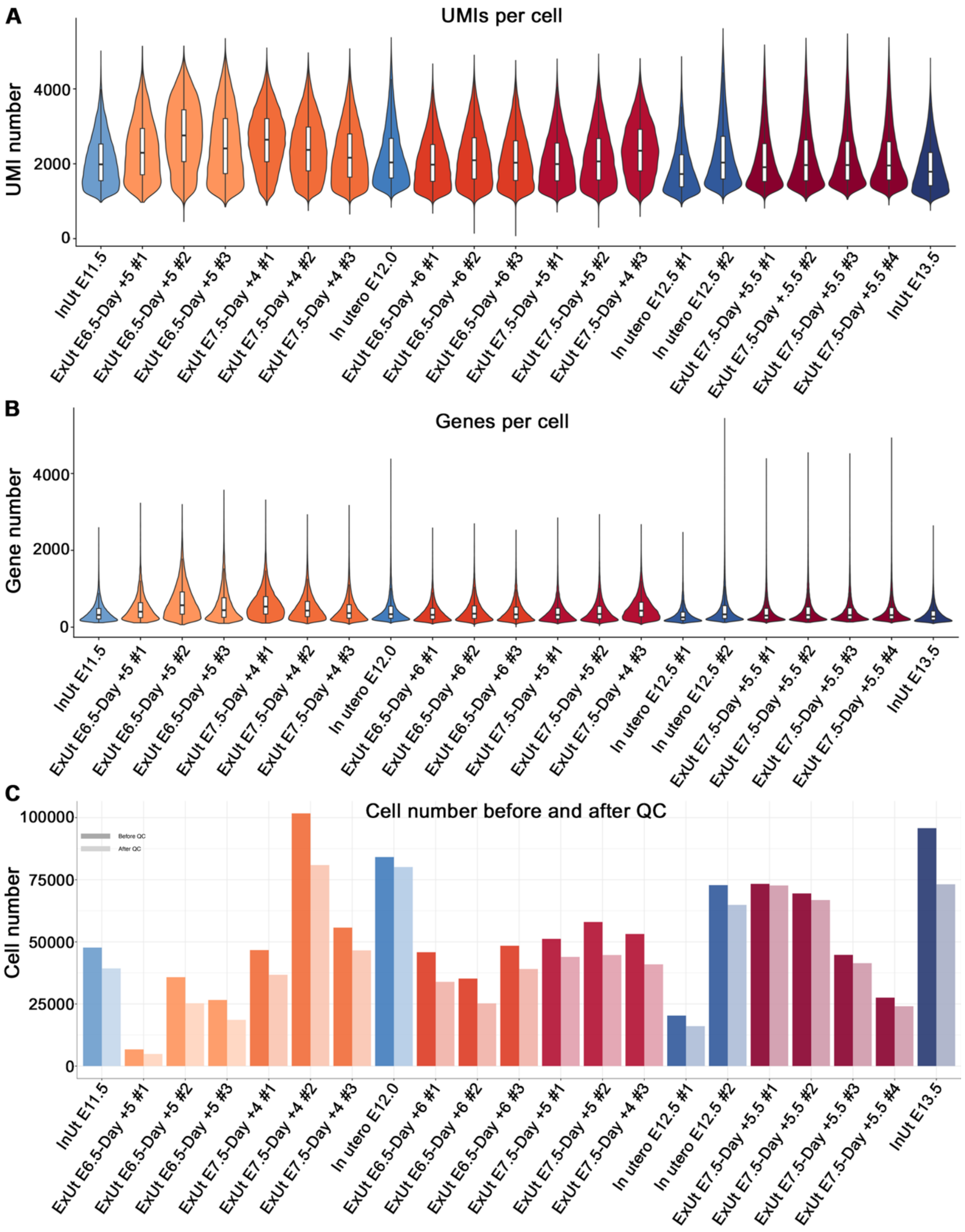


**Extended data Fig. 6. Quality control metrics for single-cell combinatorial indexing RNA-sequencing datasets. A,** Distribution of unique molecular identifiers (UMIs) per cell across all samples, showing comparable sequencing depth between ex utero (EU) and in utero (IU) embryos. **B,** Comparison of the number of detected genes per cell across samples. **C,** Total number of cells recovered before and after quality control filtering per sample used for subsequent analysis. ExUt E6.5 denotes embryos cultured from E6.5, while E7.5 ExUt designates embryos grown ex utero from E7.5.

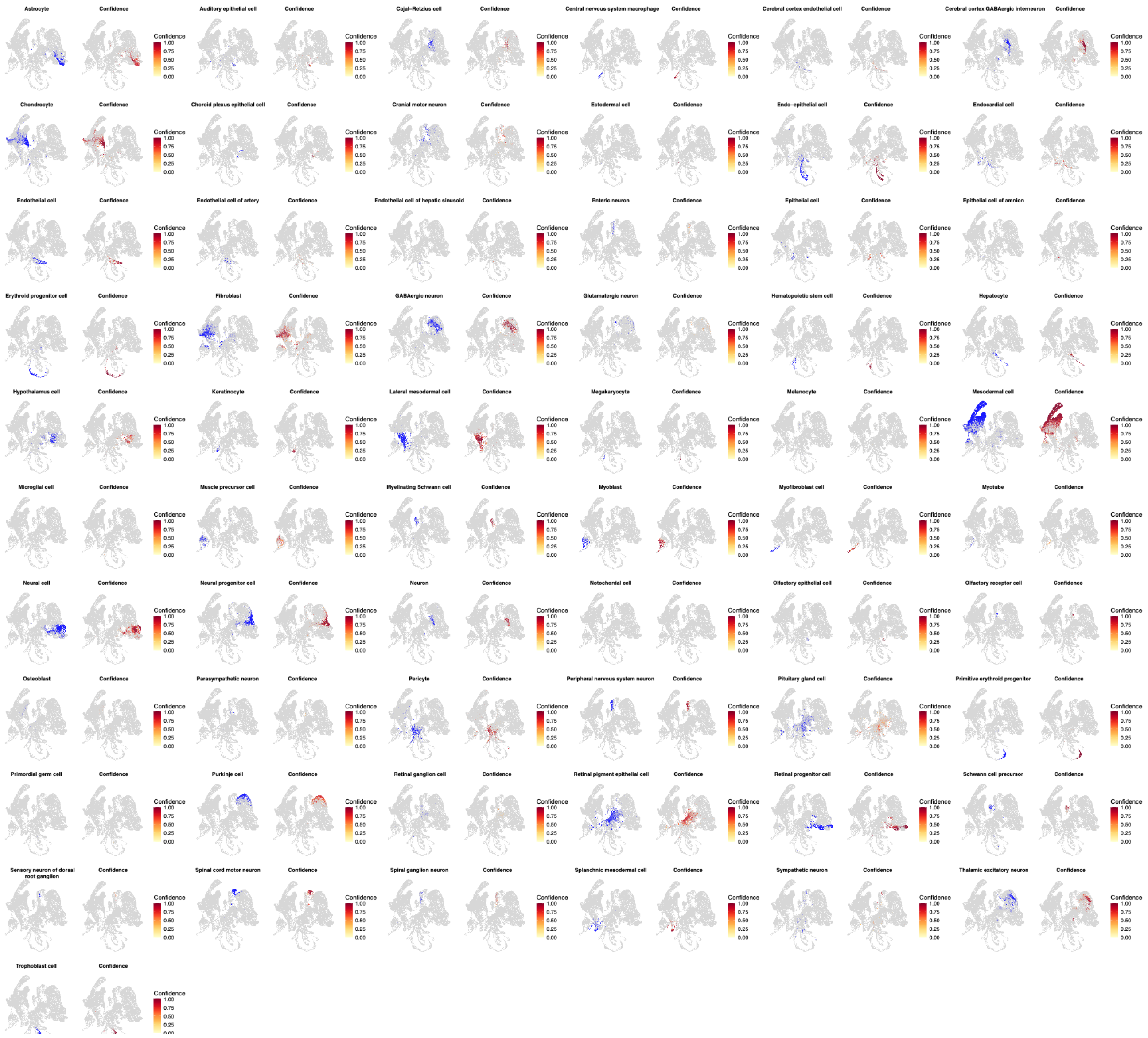


**Extended data Fig. 7. Cell type annotation confidence across all samples.** UMAP projections showing confidence scores for cell type predictions across all identified cell types. Each panel represents a specific cell type, with color intensity indicating prediction confidence (0-1 scale). Gray cells indicate low or no confidence for the respective cell type. Left panels show annotated cell location in UMAP, right panels display the prediction confidence score. Cell type annotations were transferred from a comprehensive mouse developmental reference dataset containing 11.4 million cells[58], using 1.5 million E11.5 - E12.5 cells as reference via scANVI transfer learning.

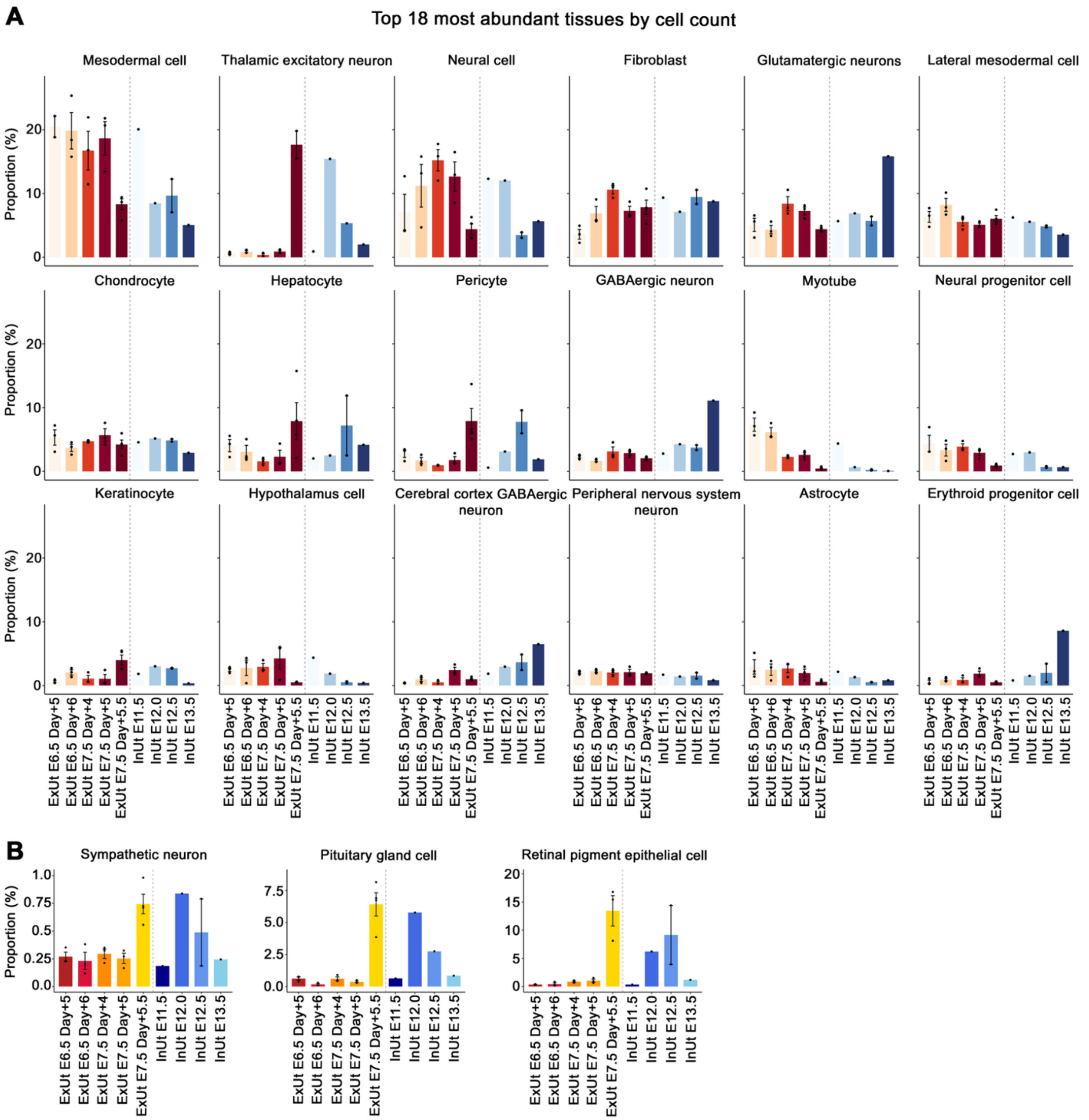


**Extended data Fig. 8. Cell type proportions across developmental stages and culture conditions. A,** Bar plots depicting the proportional abundance of the 18 most represented cell types across samples. Each panel represents one cell type, with bars colored by experimental condition and developmental endpoint. Left (orange/red bars): ex utero samples (embryos starting at E6.5 or E7.5 cultured ex utero for the indicated time). Right (blue bars): in utero control samples (E11.5, E12.0, E12.5, E13.5). **B,** Bar plots depicting the proportional abundance of three specific cell types: sympathetic neurons, pituitary gland cells, and retinal pigment epithelial cells. Y-axis indicates cell count proportion. Dots represent individual biological replicates. Error bars show standard deviation across replicates. EU, Ex Utero; IU, In Utero.

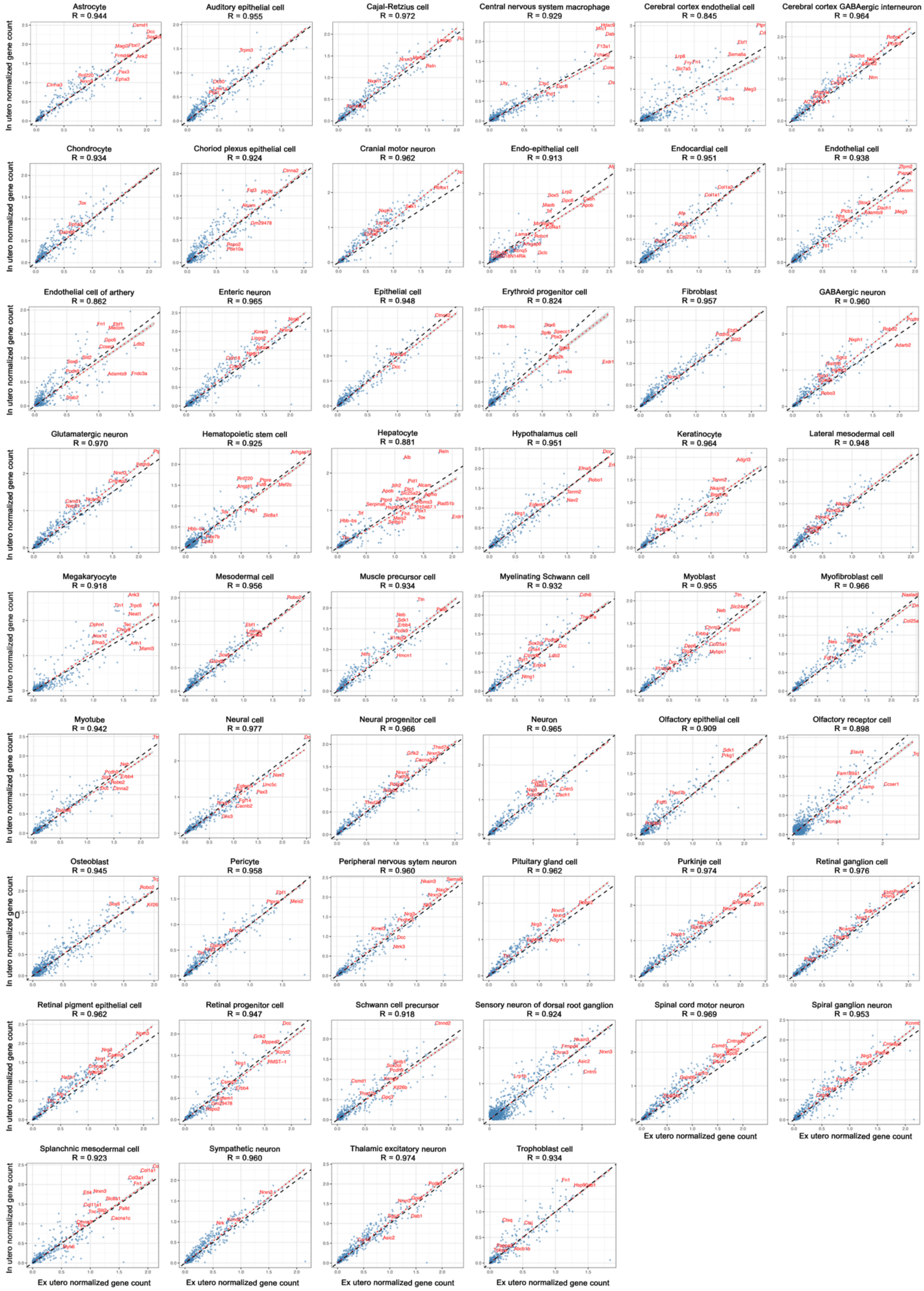

Astrocyte
R = 0.944
Auditory epithelial cell
R = 0.955
Cajal-Retzius cell
R = 0.972
Central nervous system macrophage
R = 0.929
Cerebral cortex endothelial cell
R = 0.845
Cerebral cortex GABAergic interneuron
R = 0.964
Chondrocyte
R = 0.934
Choriod plexus epithelial cell
R = 0.924
Cranial motor neuron
R = 0.962
Endo-epithelial cell
R = 0.913
Endocardial cell
R = 0.951
Endothelial cell
R = 0.938
Endothelial cell of arthery
R = 0.862
Enteric neuron
R = 0.965
Epithelial cell
R = 0.948
Erythroid progenitor cell
R = 0.824
Fibroblast
R = 0.957
GABAergic neuron
R = 0.960
Glutamatergic neuron
R = 0.970
Hematopoietic stem cell
R = 0.925
Hepatocyte
R = 0.881
Hypothalamus cell
R = 0.951
Keratinocyte
R = 0.964
Lateral mesodermal cell
R = 0.948
Megakaryocyte
R = 0.918
Mesodermal cell
R = 0.956
Muscle precursor cell
R = 0.934
Myelinating Schwann cell
R = 0.932
Myoblast
R = 0.955
Myofibroblast cell
R = 0.966
Myotube
R = 0.942
Neural cell
R = 0.977
Neural progenitor cell
R = 0.966
Neuron
R = 0.965
Olfactory epithelial cell
R = 0.909
Olfactory receptor cell
R = 0.898
Osteoblast
R = 0.945
Pericyte
R = 0.958
Peripheral nervous sytem neuron
R = 0.960
Pituitary gland cell
R = 0.962
Purkinje cell
R = 0.974
Retinal ganglion cell
R = 0.976
Retinal pigment epithelial cell
R = 0.962
Retinal progenitor cell
R = 0.947
Schwann cell precursor
R = 0.918
Sensory neuron of dorsal root ganglion
R = 0.924
Spinal cord motor neuron
R = 0.969
Spiral ganglion neuron
R = 0.953
Splanchnic mesodermal cell
R = 0.923
Sympathetic neuron
R = 0.960
Thalamic excitatory neuron
R = 0.974
Trophoblast cell
R = 0.934
In utero normalized gene count
Ex utero normalized gene count

**Extended data Fig. 9. Transcriptomic correlations between ex utero and in utero cell types (E7.5-E12.5).** Scatter plots showing gene correlations between matched cell type pairs from ex utero E7.5 + Day 5.5 (x-axis) and in utero E12.5 (y-axis) embryos. Each panel represents one cell type. Most variable genes per cluster are indicated in red. The diagonal dashed line represents perfect correlation (y = x). Spearman correlation coefficients (R) and $R^2$ values are displayed on each plot. Only cell types with ≥50 cells per condition are included. Analysis based on mean expression of top 2,000 highly variable genes in pseudobulk profiles.

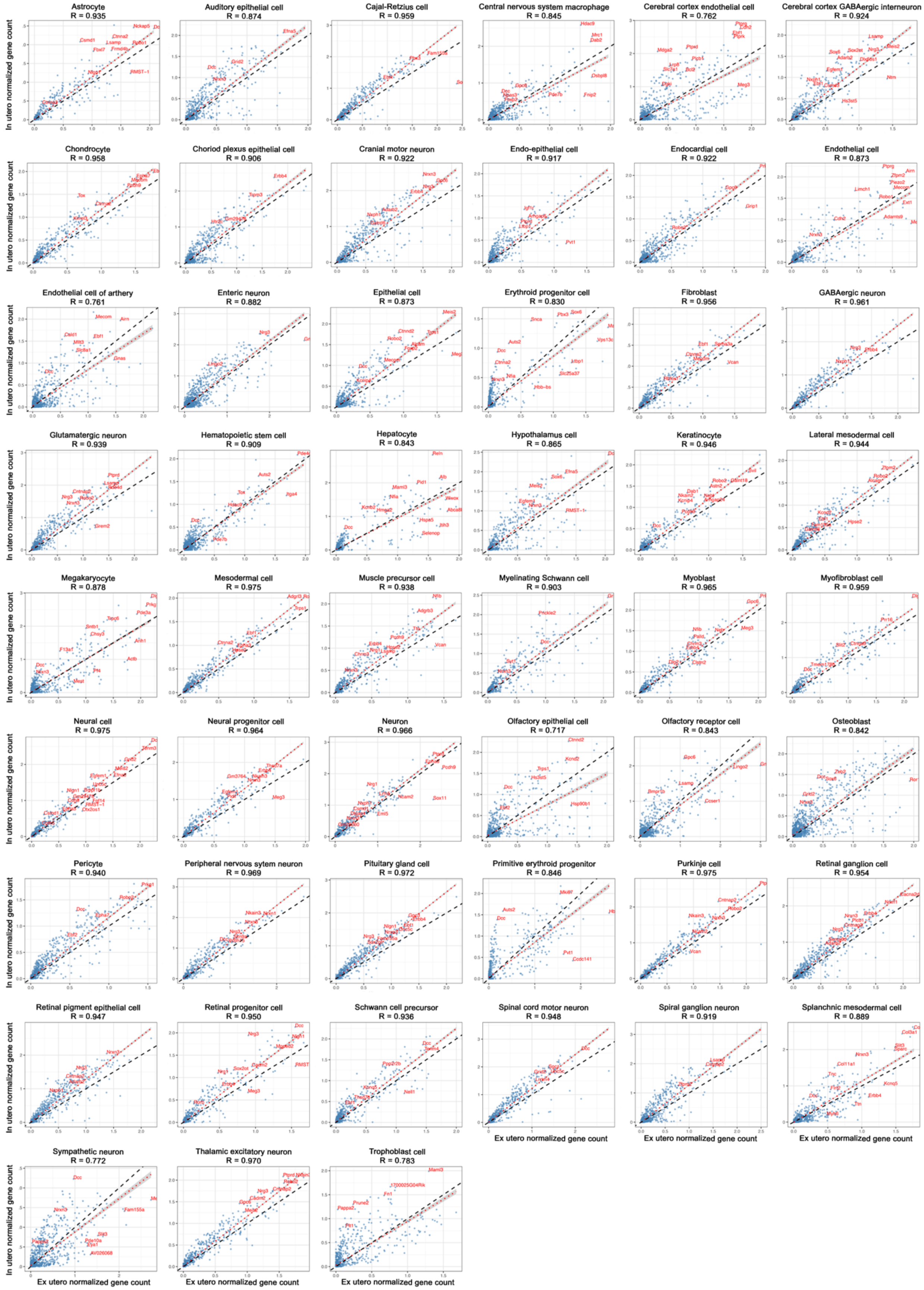

Astrocyte R = 0.935
Auditory epithelial cell R = 0.874
Cajal-Retzius cell R = 0.959
Central nervous system macrophage R = 0.845
Cerebral cortex endothelial cell R = 0.762
Cerebral cortex GABAergic interneuron R = 0.924
Chondrocyte R = 0.958
Choriod plexus epithelial cell R = 0.906
Cranial motor neuron R = 0.922
Endo-epithelial cell R = 0.917
Endocardial cell R = 0.922
Endothelial cell R = 0.873
Endothelial cell of arthery R = 0.761
Enteric neuron R = 0.882
Epithelial cell R = 0.873
Erythroid progenitor cell R = 0.830
Fibroblast R = 0.956
GABAergic neuron R = 0.961
Glutamatergic neuron R = 0.939
Hematopoietic stem cell R = 0.909
Hepatocyte R = 0.843
Hypothalamus cell R = 0.865
Keratinocyte R = 0.946
Lateral mesodermal cell R = 0.944
Megakaryocyte R = 0.878
Mesodermal cell R = 0.975
Muscle precursor cell R = 0.938
Myelinating Schwann cell R = 0.903
Myoblast R = 0.965
Myofibroblast cell R = 0.959
Neural cell R = 0.975
Neural progenitor cell R = 0.964
Neuron R = 0.966
Olfactory epithelial cell R = 0.717
Olfactory receptor cell R = 0.843
Osteoblast R = 0.842
Pericyte R = 0.940
Peripheral nervous sytem neuron R = 0.969
Pituitary gland cell R = 0.972
Primitive erythroid progenitor R = 0.846
Purkinje cell R = 0.975
Retinal ganglion cell R = 0.954
Retinal pigment epithelial cell R = 0.947
Retinal progenitor cell R = 0.950
Schwann cell precursor R = 0.936
Spinal cord motor neuron R = 0.948
Spiral ganglion neuron R = 0.919
Splanchnic mesodermal cell R = 0.889
Sympathetic neuron R = 0.772
Thalamic excitatory neuron R = 0.970
Trophoblast cell R = 0.783
In utero normalized gene count
Ex utero normalized gene count

**Extended data Fig. 10. Transcriptomic correlations between ex utero and in utero cell types (E6.5-E12.0).** Scatter plots showing gene correlations between time-matched cell type pairs from ex utero (E6.5 + Day 6) (x-axis) and in utero (E12.0) (y-axis) embryos. Most variable genes per cluster are named and shown as red dots. The diagonal dashed line represents perfect correlation (y = x). Spearman correlation coefficients (R) and $R^2$ values are displayed on each plot. Only cell types with ≥50 cells per condition are included. Analysis based on mean expression of top 2,000 highly variable genes in pseudobulk profiles.

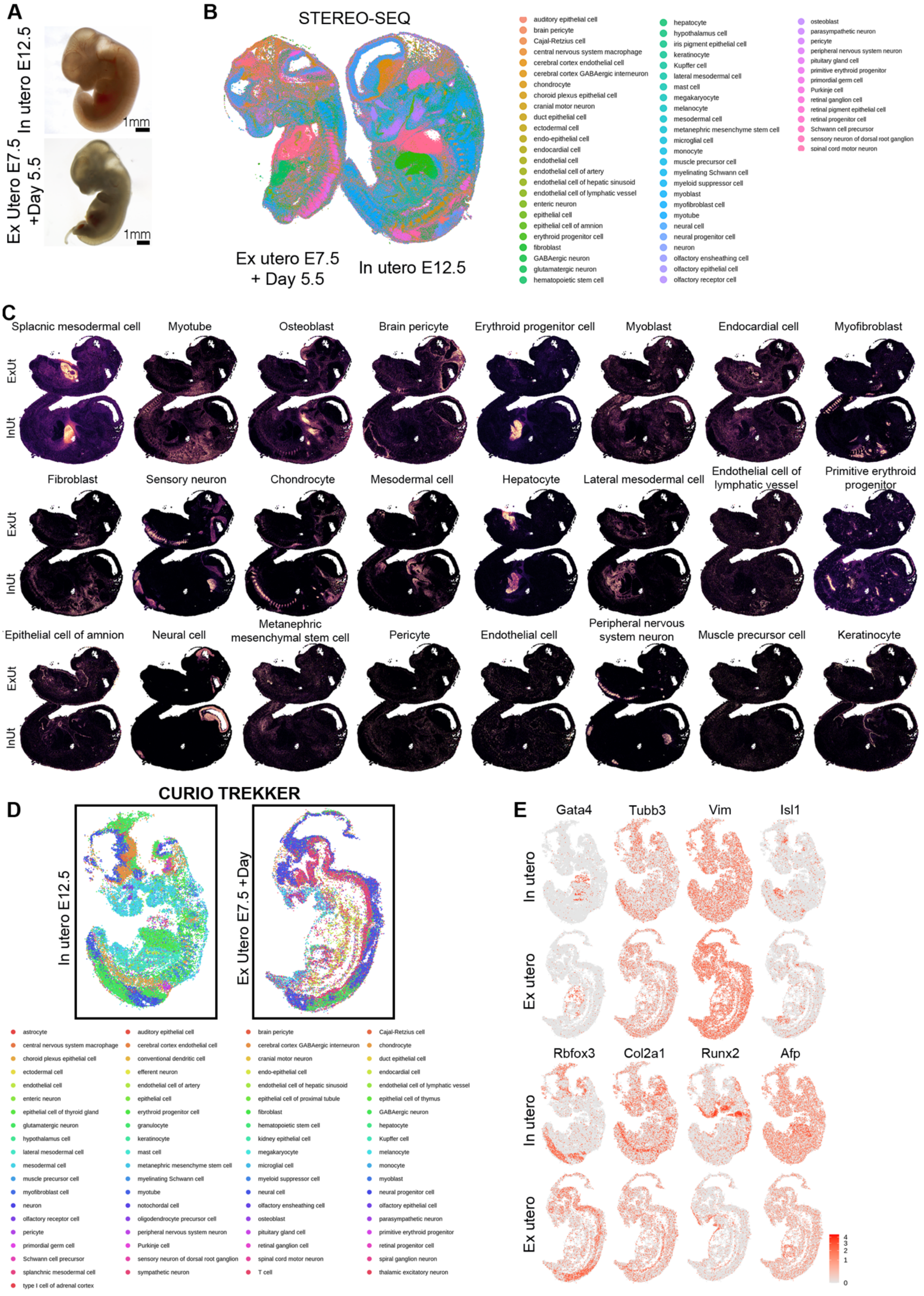
A
In utero E12.5
1mm
Ex Utero E7.5 +Day 5.5
1mm
B
STEREO-SEQ
Ex utero E7.5 + Day 5.5
In utero E12.5
auditory epithelial cell
brain pericyte
Cajal-Retzius cell
central nervous system macrophage
cerebral cortex endothelial cell
cerebral cortex GABAergic interneuron
chondrocyte
choroid plexus epithelial cell
cranial motor neuron
duct epithelial cell
ectodermal cell
endo-epithelial cell
endocardial cell
endothelial cell
endothelial cell of artery
endothelial cell of hepatic sinusoid
endothelial cell of lymphatic vessel
enteric neuron
epithelial cell
epithelial cell of amnion
erythroid progenitor cell
fibroblast
GABAergic neuron
glutamatergic neuron
hematopoietic stem cell
hepatocyte
hypothalamus cell
iris pigment epithelial cell
keratinocyte
Kupffer cell
lateral mesodermal cell
mast cell
megakaryocyte
melanocyte
mesodermal cell
metanephric mesenchyme stem cell
microglial cell
monocyte
muscle precursor cell
myelinating Schwann cell
myeloid suppressor cell
myoblast
myofibroblast cell
myotube
neural cell
neural progenitor cell
neuron
olfactory ensheathing cell
olfactory epithelial cell
olfactory receptor cell
osteoblast
parasympathetic neuron
pericyte
peripheral nervous system neuron
pituitary gland cell
primitive erythroid progenitor
primordial germ cell
Purkinje cell
retinal ganglion cell
retinal pigment epithelial cell
retinal progenitor cell
Schwann cell precursor
sensory neuron of dorsal root ganglion
spinal cord motor neuron
C
Splacnic mesodermal cell
Myotube
Osteoblast
Brain pericyte
Erythroid progenitor cell
Myoblast
Endocardial cell
Myofibroblast
ExUt
InUt
Fibroblast
Sensory neuron
Chondrocyte
Mesodermal cell
Hepatocyte
Lateral mesodermal cell
Endothelial cell of lymphatic vessel
Primitive erythroid progenitor
Epithelial cell of amnion
Neural cell
Metanephric mesenchymal stem cell
Pericyte
Endothelial cell
Peripheral nervous system neuron
Muscle precursor cell
Keratinocyte
D
CURIO TREKKER
In utero E12.5
Ex Utero E7.5 +Day
astrocyte
auditory epithelial cell
brain pericyte
Cajal-Retzius cell
central nervous system macrophage
cerebral cortex endothelial cell
cerebral cortex GABAergic interneuron
chondrocyte
choroid plexus epithelial cell
conventional dendritic cell
cranial motor neuron
duct epithelial cell
ectodermal cell
efferent neuron
endo-epithelial cell
endocardial cell
endothelial cell
endothelial cell of artery
endothelial cell of hepatic sinusoid
endothelial cell of lymphatic vessel
enteric neuron
epithelial cell
epithelial cell of proximal tubule
epithelial cell of thymus
epithelial cell of thyroid gland
erythroid progenitor cell
fibroblast
GABAergic neuron
glutamatergic neuron
granulocyte
hematopoietic stem cell
hepatocyte
hypothalamus cell
keratinocyte
kidney epithelial cell
Kupffer cell
lateral mesodermal cell
mast cell
megakaryocyte
melanocyte
mesodermal cell
metanephric mesenchyme stem cell
microglial cell
monocyte
muscle precursor cell
myelinating Schwann cell
myeloid suppressor cell
myoblast
myofibroblast cell
myotube
neural cell
neural progenitor cell
neuron
notochordal cell
olfactory ensheathing cell
olfactory epithelial cell
olfactory receptor cell
oligodendrocyte precursor cell
osteoblast
parasympathetic neuron
pericyte
peripheral nervous system neuron
pituitary gland cell
primitive erythroid progenitor
primordial germ cell
Purkinje cell
retinal ganglion cell
retinal progenitor cell
Schwann cell precursor
sensory neuron of dorsal root ganglion
spinal cord motor neuron
spiral ganglion neuron
splanchnic mesodermal cell
sympathetic neuron
T cell
thalamic excitatory neuron
type I cell of adrenal cortex
E
Gata4
Tubb3
Vim
Isl1
In utero
Ex utero
Rbfox3
Col2a1
Runx2
Afp
4
3
2
1
0

**Extended data Fig. 11. Spatial transcriptomic analysis of ex utero embryogenesis. A**, Representative bright-field images of an E12.5 embryo developed in utero and an embryo explanted at E7.5 and grown ex utero for 5.5 days. **B**, STEREO-seq spatial transcriptomic maps for in utero (E12.5) and ex utero embryos (E7.5 + Day 5.5). Dots are colored according to annotated cell types. **C,** Spatial location of top 24 most represented cell lineages identified by STEREO-seq in ex utero (E7.5 day 5.5) and in utero embryos (E12.5). **D**, Curio Trekker spatial transcriptomic maps for in utero (E12.5) and ex utero embryos (E7.5 + Day 5.5). Dots are colored according to annotated cell types. In utero and ex utero sections represent different anatomical regions in the embryo across the sagittal axis. **E,** Spatial expression patterns of representative lineage-specific genes. Gene markers highlight cardiac (Gata4), neuronal (Tubb3, Isl1), mesenchymal (Vim, Col2a1), osteogenic (Runx2), endodermal (Afp), and sensory neuron (Rbfox3) lineages. The spatial localization of these lineages observed in utero is reproduced in the ex utero samples. Colors indicate normalized gene expression levels. Scale bars, 1 mm.

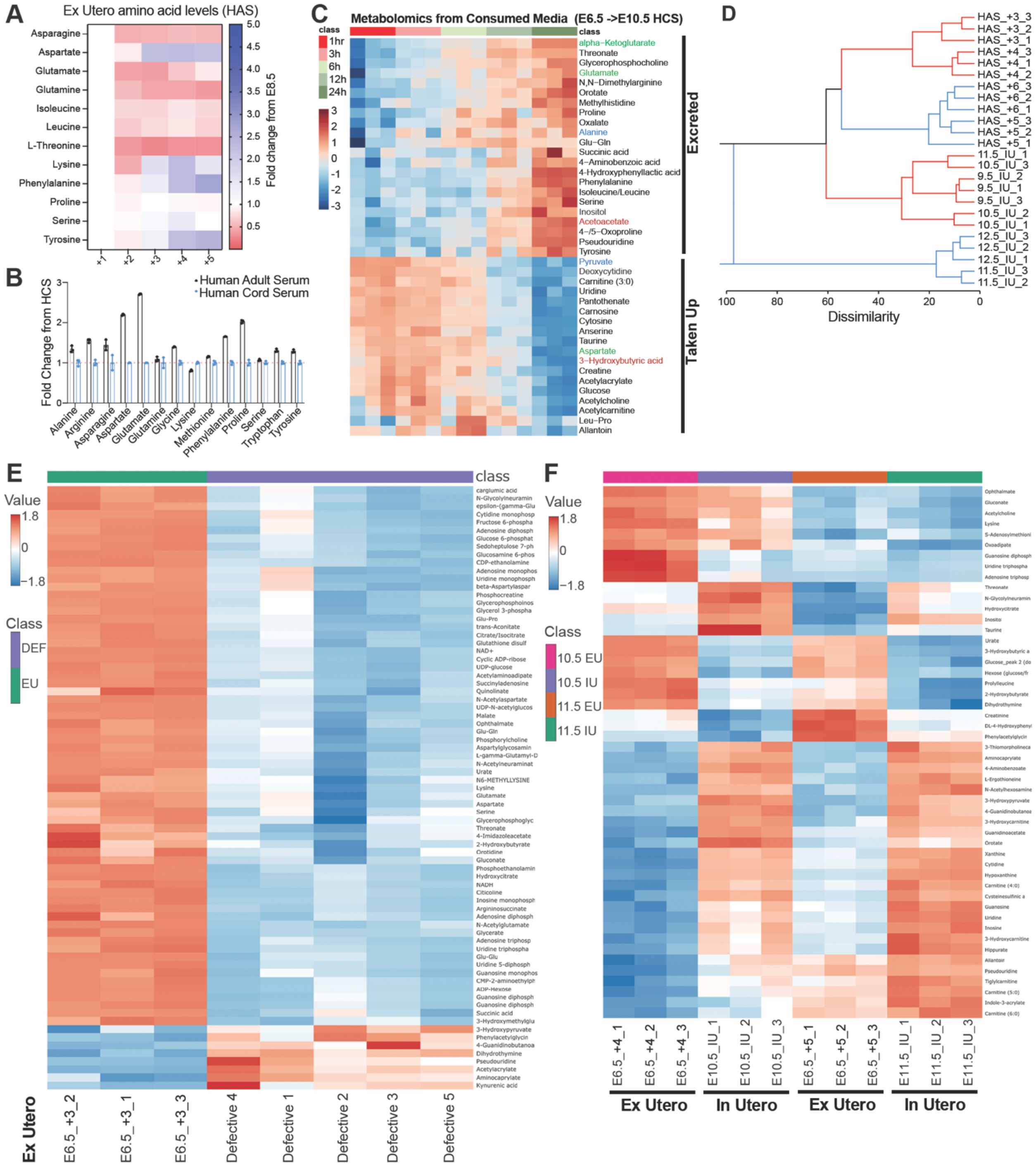


**Extended data Fig. 12. Metabolomic analysis of ex utero embryos and consumed media. A**, Metabolomic analysis of amino acid changes in ex utero cultured embryos from E7.5 though five culture days in EUCM supplemented with HAS. Values are fold change normalized to culture day 1 (E8.5). **B**, Differential amino acid composition of human adult serum (HAS) relative to human cord serum (HCS), shown as fold change normalized to HCS. Data represent mean ± S.D. **C**, Heatmap of top 40 differential metabolites in consumed EUCM through 24 hours of culture from E10.5 to E11.5 (culture started at E6.5)

demonstrating progressive excretion or uptake in ex utero embryos (normalized to culture fresh EUCM). **D,** Hierarchical clustering of global metabolomic signatures in ex utero embryos cultured from E6.5 under HAS supplementation across developmental stages (E9.5–E12.5). HAS-grown embryos cluster primarily by serum source but still display the characteristic metabolic transition at E10.5/E11.5. **E,** Heatmap of the top 75 differential metabolites distinguishing normally developing ex utero embryos from developmentally defective embryos after three days of culture from E6.5. **F,** Heatmap of the top 50 differential metabolites present in E10.5 (E6.5 +4) and E11.5 (E6.5 +5) embryos cultured ex utero (EU) compared with stage-matched in utero controls (IU). Data represent a minimum of 3 independent biological replicates.

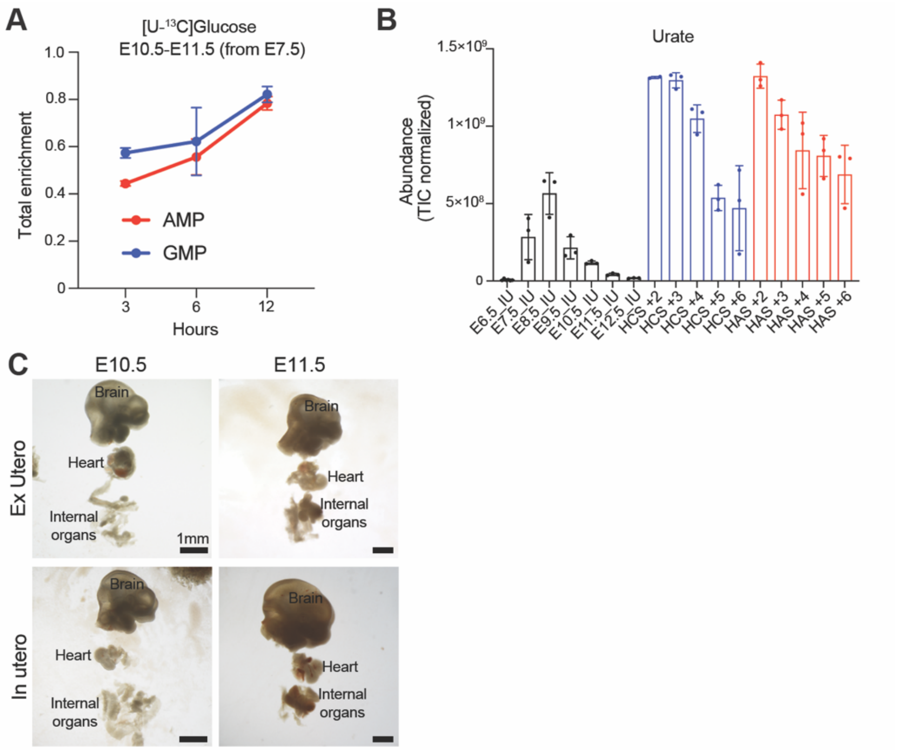


**Extended data Fig. 13. Characterization of metabolic features during the E10.5 metabolic transition in ex utero embryogenesis. A,** Accumulation of purine derivatives in ex utero embryos during the E10.5 metabolic transition. Total enrichment of [U-$^{13}$C]-Glucose-derived carbon (1 - unlabeled fraction) in AMP and GMP in a 12-hour interval during the E10.5-E11.5 transition in ex utero embryos (culture initiated at E7.5). **B**, Relative abundance and accumulation of the purine breakdown product urate through development in ex utero cultured embryos compared to in utero time-matched controls. Embryos were explanted at E6.5 and cultured with HCS or HCS for 2-6 days. Data represent mean ± S.D. n ≤ 3. **C**, Representative images of dissected organs (heart, brain and internal organs) obtained from E10.5 and E11.5 embryos grown ex utero (cultured from E7.5) and in utero controls. Images are representative at least 4 independent samples. Scale bars, 1mm.

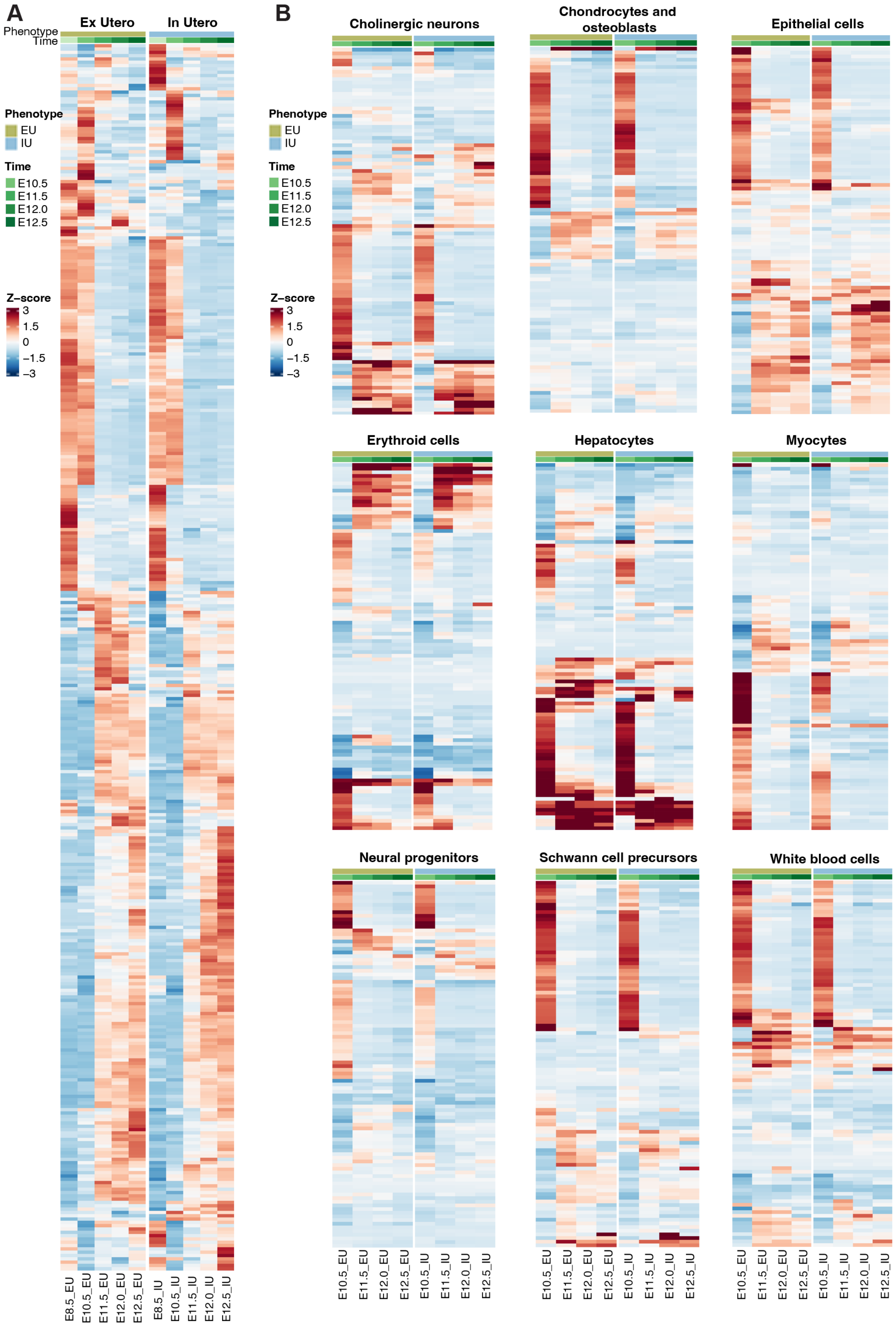
A
Ex Utero
In Utero
Phenotype
Time
Phenotype
EU
IU
Time
E10.5
E11.5
E12.0
E12.5
Z-score
3
1.5
0
−1.5
−3
E8.5_EU
E10.5_EU
E11.5_EU
E12.0_EU
E12.5_EU
E8.5_IU
E10.5_IU
E11.5_IU
E12.0_IU
E12.5_IU
B
Cholinergic neurons
Chondrocytes and osteoblasts
Epithelial cells
Erythroid cells
Hepatocytes
Myocytes
Neural progenitors
Schwann cell precursors
White blood cells
E10.5_EU
E11.5_EU
E12.0_EU
E12.5_EU
E10.5_IU
E11.5_IU
E12.0_IU
E12.5_IU

**Extended data Fig. 14. Metabolic enzyme gene expression profiles demonstrate a global metabolic transition at mid-gestation during mouse embryogenesis ex utero and in utero. A**, Heatmap showing normalized expression values of 370 metabolic enzyme genes identified by KEGG pathway mapping. Analyzed genes encode enzymes that catalyze reactions involving metabolites detected in the metabolomics dataset described above. All ex utero and in utero samples were considered, regardless of the culture start day. Rows indicate metabolism-related genes identified by KEGG pathway mapping. Hierarchical clustering of genes was performed using Euclidean distance and complete linkage. Color scale represents normalized z-score expression values. EU, Ex Utero; IU, In Utero. Heatmap incorporates datasets from E8.5, E10.5, E11.5, E12.0, and E12.5; E8.5 and E10.5 datasets were obtained from a prior study[4]. **B**, Tissue-specific heatmaps showing normalized expression values of top 100 metabolic enzyme genes ($|r| \geq 0.7$) identified by KEGG pathway mapping. Labeling as described for panel A. Heatmap incorporates datasets from E10.5, E11.5, E12.0, and E12.5 due to differences in lineages present at early stages.

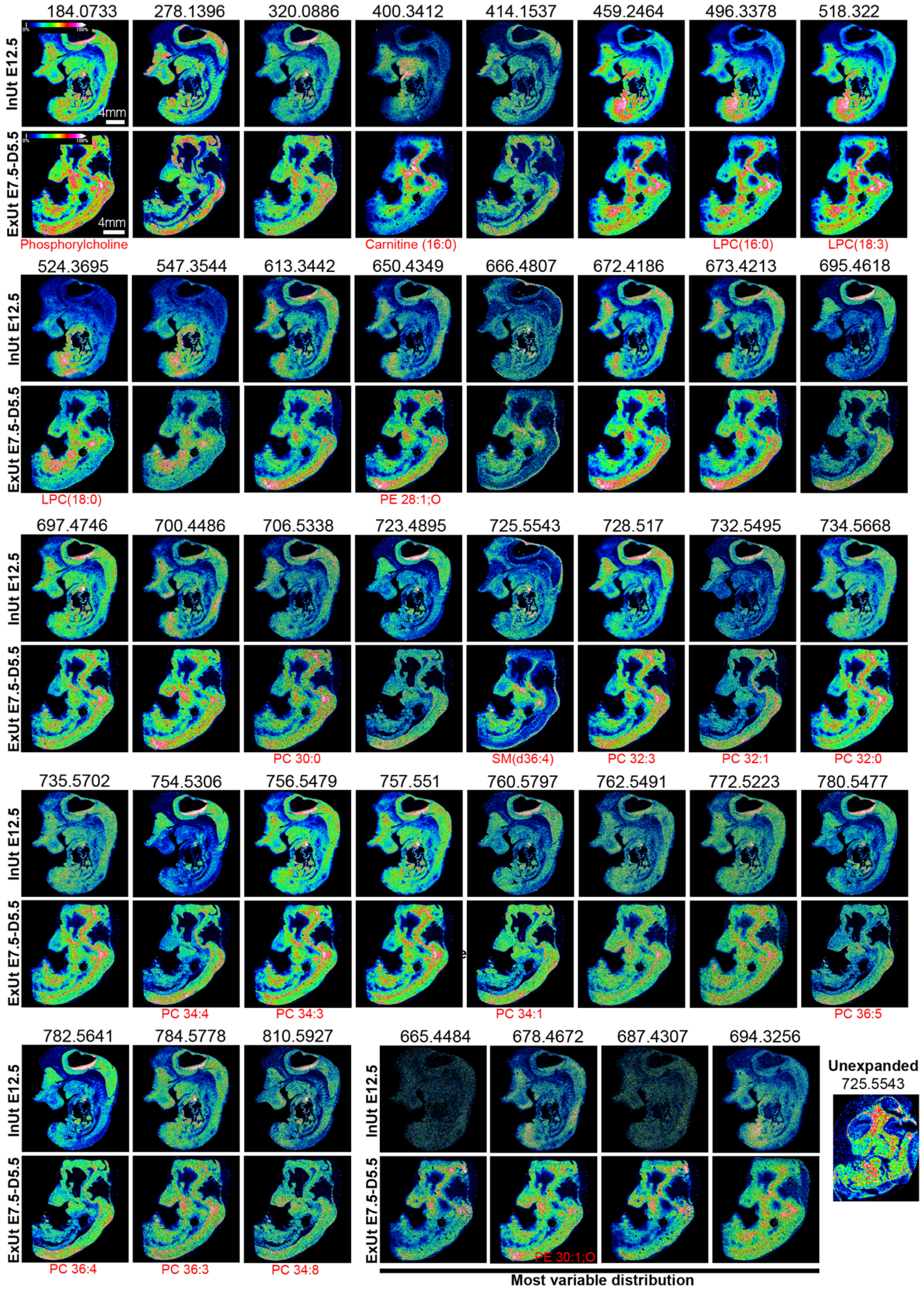
184.0733
278.1396
320.0886
400.3412
414.1537
459.2464
496.3378
518.322
InUt E12.5
ExUt E7.5-D5.5
4mm
Phosphorylcholine
Carnitine (16:0)
LPC(16:0)
LPC(18:3)
524.3695
547.3544
613.3442
650.4349
666.4807
672.4186
673.4213
695.4618
LPC(18:0)
PE 28:1;O
697.4746
700.4486
706.5338
723.4895
725.5543
728.517
732.5495
734.5668
PC 30:0
SM(d36:4)
PC 32:3
PC 32:1
PC 32:0
735.5702
754.5306
756.5479
757.551
760.5797
762.5491
772.5223
780.5477
PC 34:4
PC 34:3
PC 34:1
PC 36:5
782.5641
784.5778
810.5927
PC 36:4
PC 36:3
PC 34:8
665.4484
678.4672
687.4307
694.3256
PE 30:1;O
Most variable distribution
Unexpanded
725.5543

**Extended data Fig. 15. Tissue expansion mass spectrometry imaging of ex utero cultured mouse embryos. A**, Representative mass spectrometry images of ~3.5 fold expanded (TEMI) embryos grown ex utero for 5.5 days starting from E7.5, compared to E12.5 mouse in utero control embryos. Thirty-nine metabolites/lipids with *m/z* values ranging from 184.07 to 810.59 are shown. Four molecules with the most differential distribution between ex utero and in utero conditions are displayed at the bottom-right. All the MSI images were obtained using a 50-μm laser raster scanning with a mass error tolerance of 10 ppm. Putative metabolite identities for a subset of molecules are shown in red below the corresponding image. Scale bars, 4 mm. Color bar indicates the ion intensity scale ranging from 0 to 100%. n = 2 ex utero and 2 in utero.

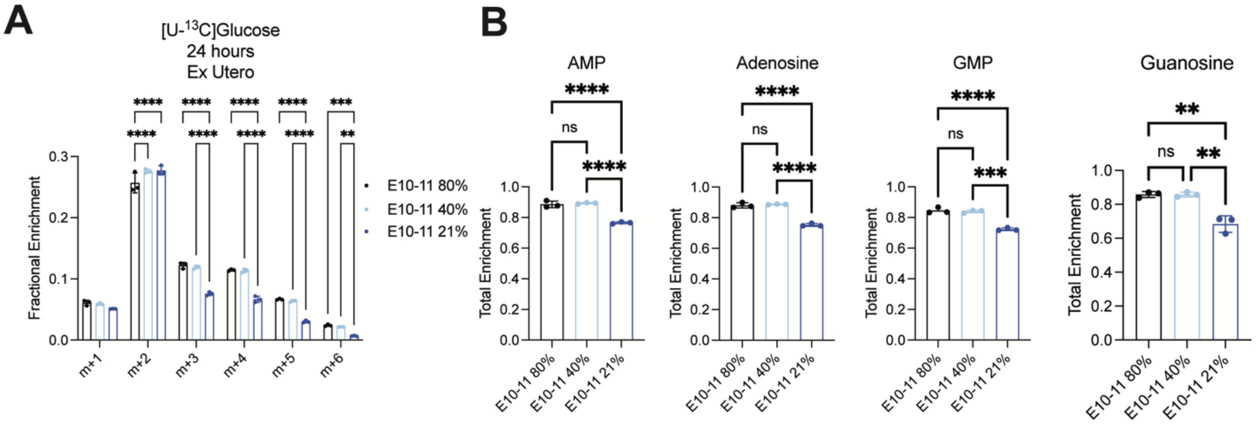


**Extended data Fig. 16. Oxygen availability and mitochondrial glucose oxidation limits purine synthesis. A.** $^{13}C$ fractional enrichment into citrate isotopologues after 24 hour [U-$^{13}C$]glucose tracing in embryos cultured ex utero under 21%, 40% or 80% $O_2$ from E10.5 to E11.5 Data are mean ± S.D. of 3 biological replicates. * = p<0.05, ** = p<0.01; *** = p<0.001; **** = p<0.0001. **B.** Total $^{13}C$ enrichment in purine metabolites after 24 hour [U-$^{13}C$]glucose tracing in embryos cultured ex utero under 21%, 40% or 80% $O_2$ from E10.5 to E11.5.

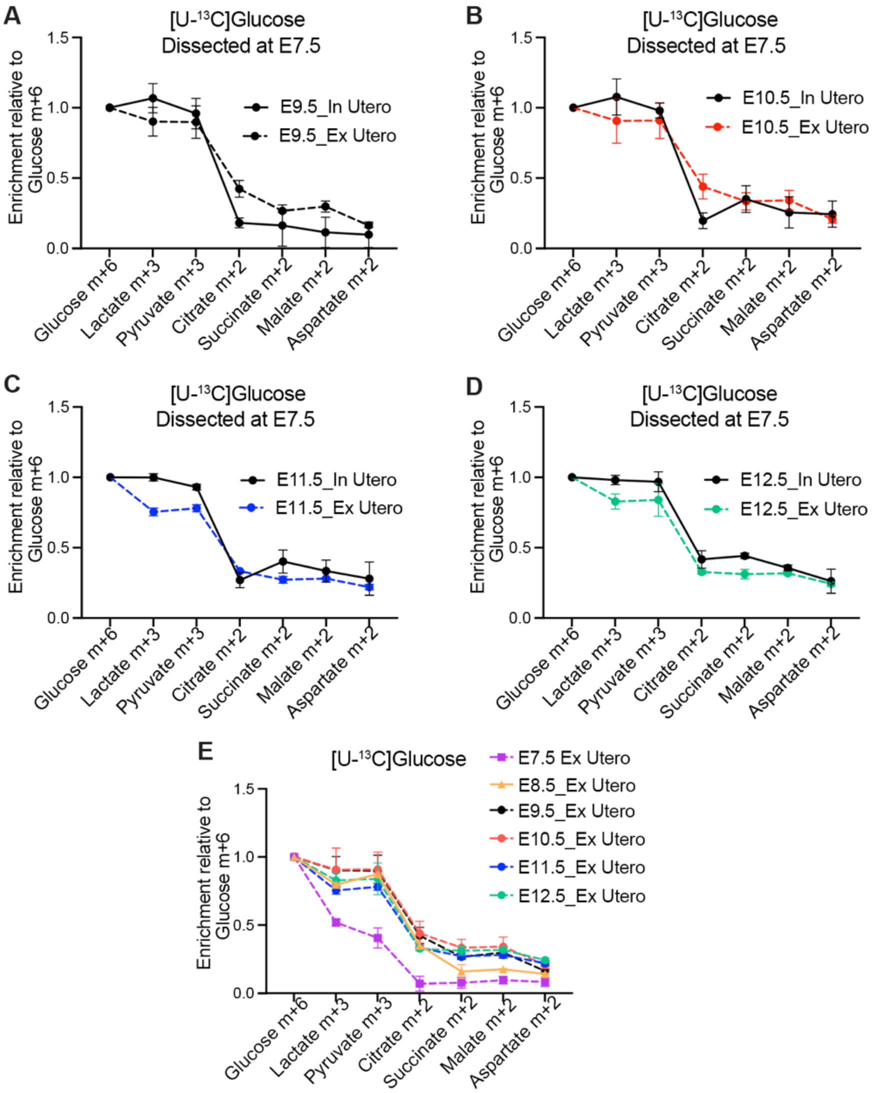


**Extended data Fig. 17. $^{13}$C-glucose tracing reveals preserved mitochondrial glucose oxidation in ex utero–grown embryos across midgestation. A–D,** [U-$^{13}$C] Glucose-derived carbon isotopic enrichment in glycolytic and TCA cycle-associated metabolites in embryos grown ex utero from E7.5 as compared to in utero embryos. Plots show fractional labeling (normalized to glucose m+6). Each panel compares stage-matched in utero (solid lines) and ex utero (dashed lines) embryos: in utero E9.5 vs ex utero day +2 (**A**), E10.5 vs day +3 (**B**), E11.5 vs day +4 (**C**), and E12.5 vs day +5 (**D**). **E**, Incorporation of [U-$^{13}$C]-labeled glucose into TCA-cycle intermediates in embryos grown ex utero from E6.5 through five days of culture. Values are normalized to Glucose m+6. Data are mean ± S.D. from 3 independent biological replicates.

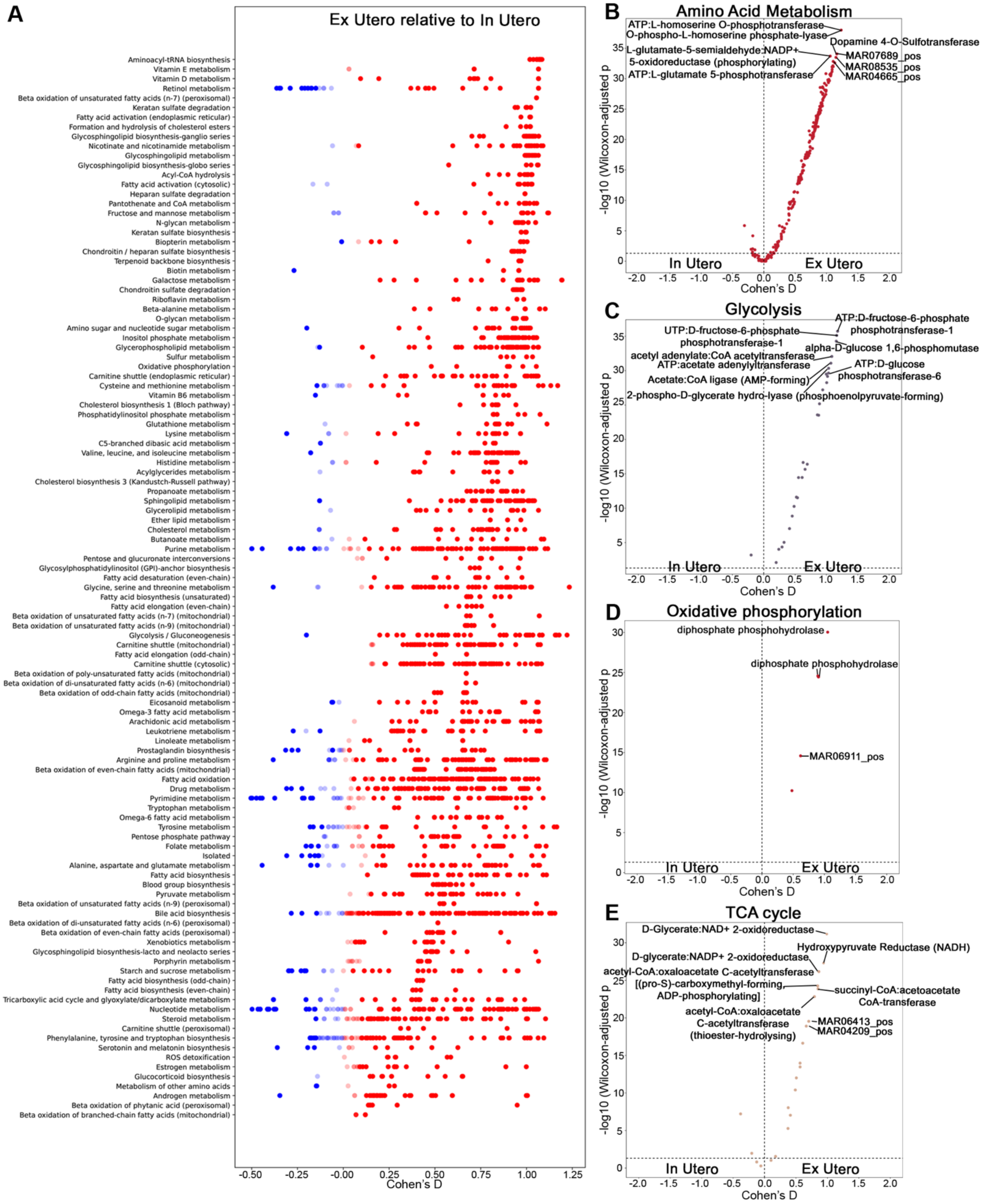


**Extended data Fig. 18. COMPASS metabolic pathway activities inferred from single-nucleus transcriptomics. A,** COMPASS-inferred metabolic pathway activities from single-nucleus transcriptomics. Scatter plot shows differential activity of metabolic reactions between ex utero and in utero conditions across all analyzed cell types and samples. Each row represents a metabolic pathway; each dot represents a metabolic reaction catalyzed by a given gene. Dot position on x-axis indicates Cohen's d (effect size), with positive values (red dots) indicating ex utero-favored pathways and negative values (blue dots) indicating in utero-favored pathways. Only significant reactions are shown. **B-E,** Volcano plots showing COMPASS-score differential activity of individual metabolic reactions within selected pathway categories: **B**, amino acid metabolism; **C**, glycolysis; **D**, oxidative phosphorylation; **E**, TCA cycle. X-axis shows Cohen's d, y-axis shows -log10(adjusted p-value). Each point represents one metabolic reaction. Key reactions are labeled. Horizontal dashed line indicates significance threshold (adjusted p-value = 0.05).

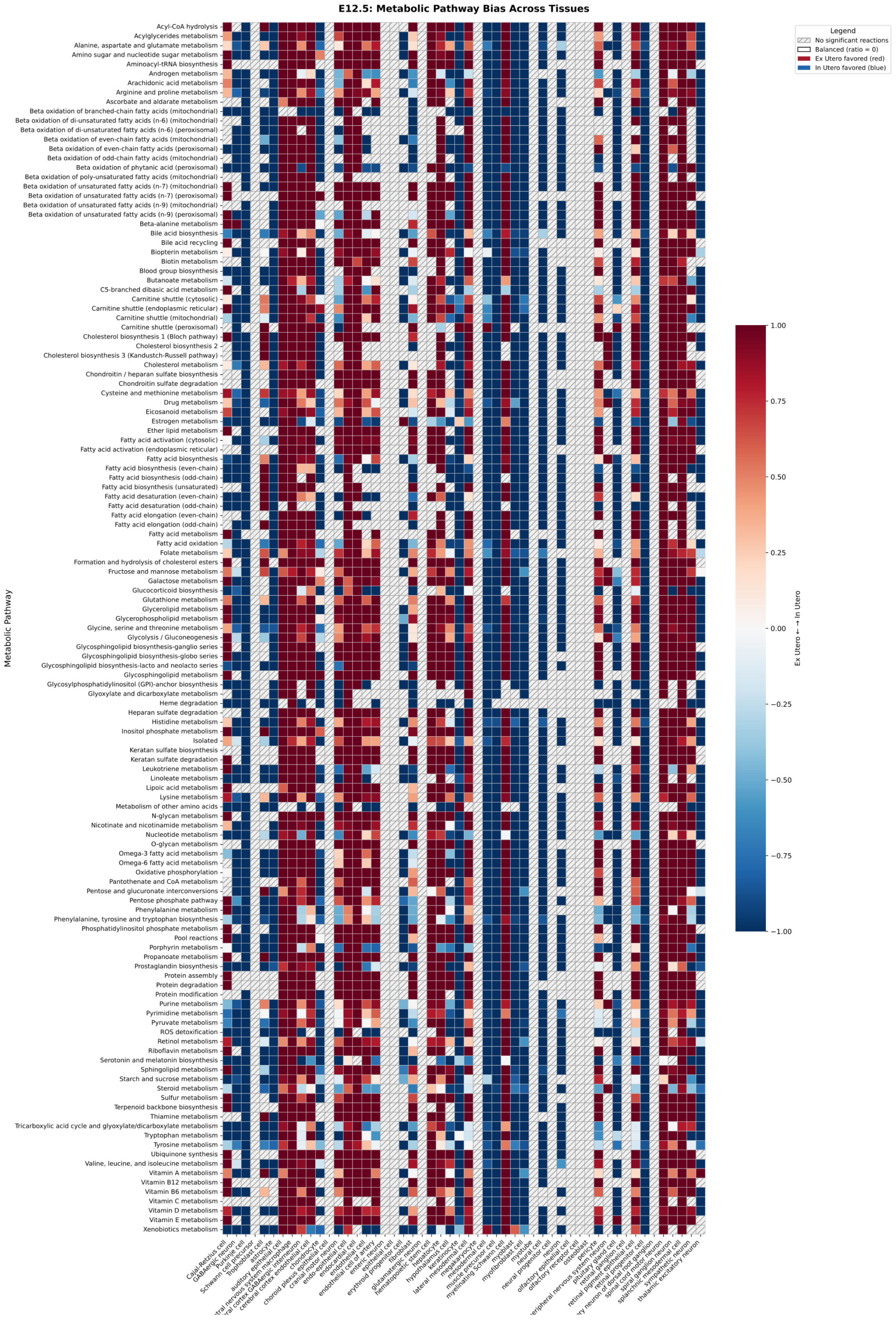
E12.5: Metabolic Pathway Bias Across Tissues
Legend
No significant reactions
Balanced (ratio = 0)
Ex Utero favored (red)
In Utero favored (blue)
Metabolic Pathway
Acyl-CoA hydrolysis
Acylglycerides metabolism
Alanine, aspartate and glutamate metabolism
Amino sugar and nucleotide sugar metabolism
Aminoacyl-tRNA biosynthesis
Androgen metabolism
Arachidonic acid metabolism
Arginine and proline metabolism
Ascorbate and aldarate metabolism
Beta oxidation of branched-chain fatty acids (mitochondrial)
Beta oxidation of di-unsaturated fatty acids (n-6) (mitochondrial)
Beta oxidation of di-unsaturated fatty acids (n-6) (peroxisomal)
Beta oxidation of even-chain fatty acids (mitochondrial)
Beta oxidation of even-chain fatty acids (peroxisomal)
Beta oxidation of odd-chain fatty acids (mitochondrial)
Beta oxidation of phytanic acid (peroxisomal)
Beta oxidation of poly-unsaturated fatty acids (mitochondrial)
Beta oxidation of unsaturated fatty acids (n-7) (mitochondrial)
Beta oxidation of unsaturated fatty acids (n-7) (peroxisomal)
Beta oxidation of unsaturated fatty acids (n-9) (mitochondrial)
Beta oxidation of unsaturated fatty acids (n-9) (peroxisomal)
Beta-alanine metabolism
Bile acid biosynthesis
Bile acid recycling
Biopterin metabolism
Biotin metabolism
Blood group biosynthesis
Butanoate metabolism
C5-branched dibasic acid metabolism
Carnitine shuttle (cytosolic)
Carnitine shuttle (endoplasmic reticular)
Carnitine shuttle (mitochondrial)
Carnitine shuttle (peroxisomal)
Cholesterol biosynthesis 1 (Bloch pathway)
Cholesterol biosynthesis 2
Cholesterol biosynthesis 3 (Kandustch-Russell pathway)
Cholesterol metabolism
Chondroitin / heparan sulfate biosynthesis
Chondroitin sulfate degradation
Cysteine and methionine metabolism
Drug metabolism
Eicosanoid metabolism
Estrogen metabolism
Ether lipid metabolism
Fatty acid activation (cytosolic)
Fatty acid activation (endoplasmic reticular)
Fatty acid biosynthesis
Fatty acid biosynthesis (even-chain)
Fatty acid biosynthesis (odd-chain)
Fatty acid biosynthesis (unsaturated)
Fatty acid desaturation (even-chain)
Fatty acid desaturation (odd-chain)
Fatty acid elongation (even-chain)
Fatty acid elongation (odd-chain)
Fatty acid metabolism
Fatty acid oxidation
Folate metabolism
Formation and hydrolysis of cholesterol esters
Fructose and mannose metabolism
Galactose metabolism
Glucocorticoid biosynthesis
Glutathione metabolism
Glycerolipid metabolism
Glycerophospholipid metabolism
Glycine, serine and threonine metabolism
Glycolysis / Gluconeogenesis
Glycosphingolipid biosynthesis-ganglio series
Glycosphingolipid biosynthesis-globo series
Glycosphingolipid biosynthesis-lacto and neolacto series
Glycosphingolipid metabolism
Glycosylphosphatidylinositol (GPI)-anchor biosynthesis
Glyoxylate and dicarboxylate metabolism
Heme degradation
Heparan sulfate degradation
Histidine metabolism
Inositol phosphate metabolism
Isolated
Keratan sulfate biosynthesis
Keratan sulfate degradation
Leukotriene metabolism
Linoleate metabolism
Lipoic acid metabolism
Lysine metabolism
Metabolism of other amino acids
N-glycan metabolism
Nicotinate and nicotinamide metabolism
Nucleotide metabolism
O-glycan metabolism
Omega-3 fatty acid metabolism
Omega-6 fatty acid metabolism
Oxidative phosphorylation
Pantothenate and CoA metabolism
Pentose and glucuronate interconversions
Pentose phosphate pathway
Phenylalanine metabolism
Phenylalanine, tyrosine and tryptophan biosynthesis
Phosphatidylinositol phosphate metabolism
Pool reactions
Porphyrin metabolism
Propanoate metabolism
Prostaglandin biosynthesis
Protein assembly
Protein degradation
Protein modification
Purine metabolism
Pyrimidine metabolism
Pyruvate metabolism
ROS detoxification
Retinol metabolism
Riboflavin metabolism
Serotonin and melatonin biosynthesis
Sphingolipid metabolism
Starch and sucrose metabolism
Steroid metabolism
Sulfur metabolism
Terpenoid backbone biosynthesis
Thiamine metabolism
Tricarboxylic acid cycle and glyoxylate/dicarboxylate metabolism
Tryptophan metabolism
Tyrosine metabolism
Ubiquinone synthesis
Valine, leucine, and isoleucine metabolism
Vitamin A metabolism
Vitamin B12 metabolism
Vitamin B6 metabolism
Vitamin C metabolism
Vitamin D metabolism
Vitamin E metabolism
Xenobiotics metabolism
Cajal-Retzius cell
GABAergic neuron
Purkinje cell
Schwann cell precursor
Trophoblast cell
astrocyte
auditory epithelial cell
cerebral cortex endothelial cell
chondrocyte
choroid plexus epithelial cell
cranial motor neuron
endo-epithelial cell
endocardial cell
endothelial cell
endothelial cell of artery
enteric neuron
epithelial cell
erythroid progenitor cell
fibroblast
glutamatergic neuron
hematopoietic stem cell
hepatocyte
hypothalamus cell
keratinocyte
lateral mesodermal cell
megakaryocyte
mesodermal cell
muscle precursor cell
myelinating Schwann cell
myoblast
myofibroblast cell
myotube
neural cell
neural progenitor cell
neuron
olfactory epithelial cell
olfactory receptor cell
osteoblast
pericyte
pituitary gland cell
retinal ganglion cell
retinal pigment epithelial cell
retinal progenitor cell
spinal cord motor neuron
spiral ganglion neuron
splanchnic mesodermal cell
sympathetic neuron
thalamic excitatory neuron
1.00
0.75
0.50
0.25
0.00
−0.25
−0.50
−0.75
−1.00
Ex Utero ← → In Utero

**Extended data Fig. 19. Cell-type-resolved metabolic pathway bias in E7.5 embryos cultured ex utero for 5.5 days.** Heatmap of COMPASS-inferred metabolic pathway activities across annotated cell types in embryos explanted at E7.5 and cultured ex utero for 5.5 days, compared to stage-matched in utero controls. Rows correspond to metabolic pathways and columns to annotated cell types. Red = ex utero-favored pathways, blue = in utero-favored pathways, white = balanced metabolism, gray with hatching = insufficient data.

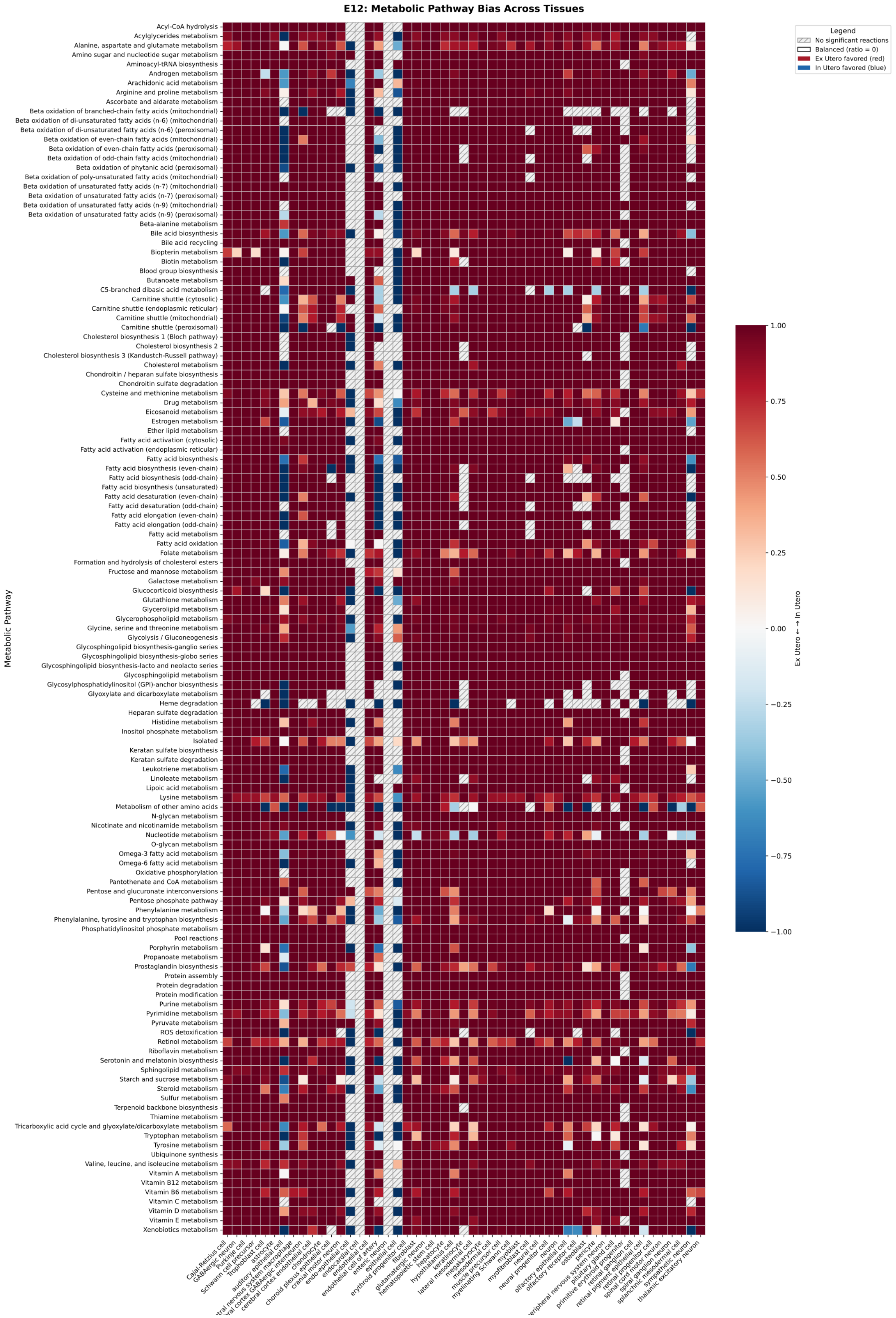
E12: Metabolic Pathway Bias Across Tissues
Metabolic Pathway
Acyl-CoA hydrolysis
Acylglycerides metabolism
Alanine, aspartate and glutamate metabolism
Amino sugar and nucleotide sugar metabolism
Aminoacyl-tRNA biosynthesis
Androgen metabolism
Arachidonic acid metabolism
Arginine and proline metabolism
Ascorbate and aldarate metabolism
Beta oxidation of branched-chain fatty acids (mitochondrial)
Beta oxidation of di-unsaturated fatty acids (n-6) (mitochondrial)
Beta oxidation of di-unsaturated fatty acids (n-6) (peroxisomal)
Beta oxidation of even-chain fatty acids (mitochondrial)
Beta oxidation of even-chain fatty acids (peroxisomal)
Beta oxidation of odd-chain fatty acids (mitochondrial)
Beta oxidation of phytanic acid (peroxisomal)
Beta oxidation of poly-unsaturated fatty acids (mitochondrial)
Beta oxidation of unsaturated fatty acids (n-7) (mitochondrial)
Beta oxidation of unsaturated fatty acids (n-7) (peroxisomal)
Beta oxidation of unsaturated fatty acids (n-9) (mitochondrial)
Beta oxidation of unsaturated fatty acids (n-9) (peroxisomal)
Beta-alanine metabolism
Bile acid biosynthesis
Bile acid recycling
Biopterin metabolism
Biotin metabolism
Blood group biosynthesis
Butanoate metabolism
C5-branched dibasic acid metabolism
Carnitine shuttle (cytosolic)
Carnitine shuttle (endoplasmic reticular)
Carnitine shuttle (mitochondrial)
Carnitine shuttle (peroxisomal)
Cholesterol biosynthesis 1 (Bloch pathway)
Cholesterol biosynthesis 2
Cholesterol biosynthesis 3 (Kandustch-Russell pathway)
Cholesterol metabolism
Chondroitin / heparan sulfate biosynthesis
Chondroitin sulfate degradation
Cysteine and methionine metabolism
Drug metabolism
Eicosanoid metabolism
Estrogen metabolism
Ether lipid metabolism
Fatty acid activation (cytosolic)
Fatty acid activation (endoplasmic reticular)
Fatty acid biosynthesis
Fatty acid biosynthesis (even-chain)
Fatty acid biosynthesis (odd-chain)
Fatty acid biosynthesis (unsaturated)
Fatty acid desaturation (even-chain)
Fatty acid desaturation (odd-chain)
Fatty acid elongation (even-chain)
Fatty acid elongation (odd-chain)
Fatty acid metabolism
Fatty acid oxidation
Folate metabolism
Formation and hydrolysis of cholesterol esters
Fructose and mannose metabolism
Galactose metabolism
Glucocorticoid biosynthesis
Glutathione metabolism
Glycerolipid metabolism
Glycerophospholipid metabolism
Glycine, serine and threonine metabolism
Glycolysis / Gluconeogenesis
Glycosphingolipid biosynthesis-ganglio series
Glycosphingolipid biosynthesis-globo series
Glycosphingolipid biosynthesis-lacto and neolacto series
Glycosphingolipid metabolism
Glycosylphosphatidylinositol (GPI)-anchor biosynthesis
Glyoxylate and dicarboxylate metabolism
Heme degradation
Heparan sulfate degradation
Histidine metabolism
Inositol phosphate metabolism
Isolated
Keratan sulfate biosynthesis
Keratan sulfate degradation
Leukotriene metabolism
Linoleate metabolism
Lipoic acid metabolism
Lysine metabolism
Metabolism of other amino acids
N-glycan metabolism
Nicotinate and nicotinamide metabolism
Nucleotide metabolism
O-glycan metabolism
Omega-3 fatty acid metabolism
Omega-6 fatty acid metabolism
Oxidative phosphorylation
Pantothenate and CoA metabolism
Pentose and glucuronate interconversions
Pentose phosphate pathway
Phenylalanine metabolism
Phenylalanine, tyrosine and tryptophan biosynthesis
Phosphatidylinositol phosphate metabolism
Pool reactions
Porphyrin metabolism
Propanoate metabolism
Prostaglandin biosynthesis
Protein assembly
Protein degradation
Protein modification
Purine metabolism
Pyrimidine metabolism
Pyruvate metabolism
ROS detoxification
Retinol metabolism
Riboflavin metabolism
Serotonin and melatonin biosynthesis
Sphingolipid metabolism
Starch and sucrose metabolism
Steroid metabolism
Sulfur metabolism
Terpenoid backbone biosynthesis
Thiamine metabolism
Tricarboxylic acid cycle and glyoxylate/dicarboxylate metabolism
Tryptophan metabolism
Tyrosine metabolism
Ubiquinone synthesis
Valine, leucine, and isoleucine metabolism
Vitamin A metabolism
Vitamin B12 metabolism
Vitamin B6 metabolism
Vitamin C metabolism
Vitamin D metabolism
Vitamin E metabolism
Xenobiotics metabolism
Cajal-Retzius cell
GABAergic neuron
Purkinje cell
Schwann cell precursor
Trophoblast cell
astrocyte
auditory epithelial cell
tral nervous system macrophage
ral cortex GABAergic interneuron
cerebral cortex endothelial cell
chondrocyte
choroid plexus epithelial cell
cranial motor neuron
endo-epithelial cell
endocardial cell
endothelial cell
endothelial cell of artery
enteric neuron
epithelial cell
erythroid progenitor cell
fibroblast
glutamatergic neuron
hematopoietic stem cell
hepatocyte
hypothalamus cell
keratinocyte
lateral mesodermal cell
megakaryocyte
mesodermal cell
muscle precursor cell
myelinating Schwann cell
myoblast
myofibroblast cell
neural cell
neural progenitor cell
neuron
olfactory epithelial cell
olfactory receptor cell
osteoblast
pericyte
eripheral nervous system neuron
pituitary gland cell
primitive erythroid progenitor
retinal ganglion cell
retinal pigment epithelial cell
retinal progenitor cell
spinal cord motor neuron
spiral ganglion neuron
splanchnic mesodermal cell
sympathetic neuron
thalamic excitatory neuron
Legend
No significant reactions
Balanced (ratio = 0)
Ex Utero favored (red)
In Utero favored (blue)
Ex Utero ← → In Utero
1.00
0.75
0.50
0.25
0.00
−0.25
−0.50
−0.75
−1.00

**Extended data Fig. 20. Metabolic pathway bias across cell types in E6.5 embryos grown ex utero for six days.** COMPASS-based heatmap showing tissue-specific metabolic pathway activities in embryos explanted at E6.5 and cultured ex utero for six days, compared with E12.0 in utero controls. Rows correspond to metabolic pathways and columns to annotated cell types. Red = ex utero-favored pathways, blue = in utero-favored pathways, white = balanced metabolism, gray with hatching = insufficient data.

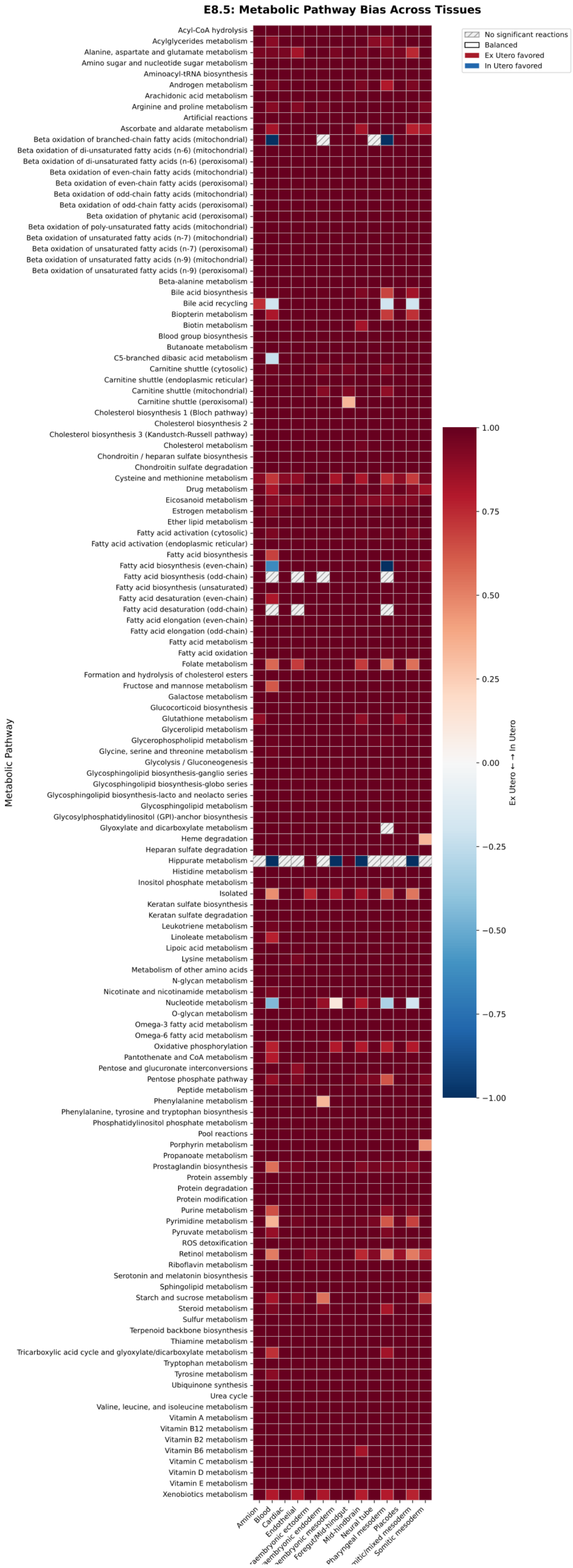
E8.5: Metabolic Pathway Bias Across Tissues
No significant reactions
Balanced
Ex Utero favored
In Utero favored
Metabolic Pathway
Acyl-CoA hydrolysis
Acylglycerides metabolism
Alanine, aspartate and glutamate metabolism
Amino sugar and nucleotide sugar metabolism
Aminoacyl-tRNA biosynthesis
Androgen metabolism
Arachidonic acid metabolism
Arginine and proline metabolism
Artificial reactions
Ascorbate and aldarate metabolism
Beta oxidation of branched-chain fatty acids (mitochondrial)
Beta oxidation of di-unsaturated fatty acids (n-6) (mitochondrial)
Beta oxidation of di-unsaturated fatty acids (n-6) (peroxisomal)
Beta oxidation of even-chain fatty acids (mitochondrial)
Beta oxidation of even-chain fatty acids (peroxisomal)
Beta oxidation of odd-chain fatty acids (mitochondrial)
Beta oxidation of odd-chain fatty acids (peroxisomal)
Beta oxidation of phytanic acid (peroxisomal)
Beta oxidation of poly-unsaturated fatty acids (mitochondrial)
Beta oxidation of unsaturated fatty acids (n-7) (mitochondrial)
Beta oxidation of unsaturated fatty acids (n-7) (peroxisomal)
Beta oxidation of unsaturated fatty acids (n-9) (mitochondrial)
Beta oxidation of unsaturated fatty acids (n-9) (peroxisomal)
Beta-alanine metabolism
Bile acid biosynthesis
Bile acid recycling
Biopterin metabolism
Biotin metabolism
Blood group biosynthesis
Butanoate metabolism
C5-branched dibasic acid metabolism
Carnitine shuttle (cytosolic)
Carnitine shuttle (endoplasmic reticular)
Carnitine shuttle (mitochondrial)
Carnitine shuttle (peroxisomal)
Cholesterol biosynthesis 1 (Bloch pathway)
Cholesterol biosynthesis 2
Cholesterol biosynthesis 3 (Kandustch-Russell pathway)
Cholesterol metabolism
Chondroitin / heparan sulfate biosynthesis
Chondroitin sulfate degradation
Cysteine and methionine metabolism
Drug metabolism
Eicosanoid metabolism
Estrogen metabolism
Ether lipid metabolism
Fatty acid activation (cytosolic)
Fatty acid activation (endoplasmic reticular)
Fatty acid biosynthesis
Fatty acid biosynthesis (even-chain)
Fatty acid biosynthesis (odd-chain)
Fatty acid biosynthesis (unsaturated)
Fatty acid desaturation (even-chain)
Fatty acid desaturation (odd-chain)
Fatty acid elongation (even-chain)
Fatty acid elongation (odd-chain)
Fatty acid metabolism
Fatty acid oxidation
Folate metabolism
Formation and hydrolysis of cholesterol esters
Fructose and mannose metabolism
Galactose metabolism
Glucocorticoid biosynthesis
Glutathione metabolism
Glycerolipid metabolism
Glycerophospholipid metabolism
Glycine, serine and threonine metabolism
Glycolysis / Gluconeogenesis
Glycosphingolipid biosynthesis-ganglio series
Glycosphingolipid biosynthesis-globo series
Glycosphingolipid biosynthesis-lacto and neolacto series
Glycosphingolipid metabolism
Glycosylphosphatidylinositol (GPI)-anchor biosynthesis
Glyoxylate and dicarboxylate metabolism
Heme degradation
Heparan sulfate degradation
Hippurate metabolism
Histidine metabolism
Inositol phosphate metabolism
Isolated
Keratan sulfate biosynthesis
Keratan sulfate degradation
Leukotriene metabolism
Linoleate metabolism
Lipoic acid metabolism
Lysine metabolism
Metabolism of other amino acids
N-glycan metabolism
Nicotinate and nicotinamide metabolism
Nucleotide metabolism
O-glycan metabolism
Omega-3 fatty acid metabolism
Omega-6 fatty acid metabolism
Oxidative phosphorylation
Pantothenate and CoA metabolism
Pentose and glucuronate interconversions
Pentose phosphate pathway
Peptide metabolism
Phenylalanine metabolism
Phenylalanine, tyrosine and tryptophan biosynthesis
Phosphatidylinositol phosphate metabolism
Pool reactions
Porphyrin metabolism
Propanoate metabolism
Prostaglandin biosynthesis
Protein assembly
Protein degradation
Protein modification
Purine metabolism
Pyrimidine metabolism
Pyruvate metabolism
ROS detoxification
Retinol metabolism
Riboflavin metabolism
Serotonin and melatonin biosynthesis
Sphingolipid metabolism
Starch and sucrose metabolism
Steroid metabolism
Sulfur metabolism
Terpenoid backbone biosynthesis
Thiamine metabolism
Tricarboxylic acid cycle and glyoxylate/dicarboxylate metabolism
Tryptophan metabolism
Tyrosine metabolism
Ubiquinone synthesis
Urea cycle
Valine, leucine, and isoleucine metabolism
Vitamin A metabolism
Vitamin B12 metabolism
Vitamin B2 metabolism
Vitamin B6 metabolism
Vitamin C metabolism
Vitamin D metabolism
Vitamin E metabolism
Xenobiotics metabolism
Amnion
Blood
Cardiac
Endothelial
Foregut/Mid-hindgut
Mid-hindbrain
Neural tube
Pharyngeal mesoderm
Placodes
Somitic mesoderm
1.00
0.75
0.50
0.25
0.00
−0.25
−0.50
−0.75
−1.00
Ex Utero ← → In Utero

**Extended data Fig. 21. Early metabolic pathway bias during pre-gastrulation to early organogenesis.** Heatmap showing COMPASS-inferred tissue-specific metabolic pathway activities in embryos cultured ex utero for two days starting at E6.5 and compared to in utero embryos at E8.5. Data were obtained from a previously published study[4]. Rows correspond to metabolic pathways and columns to annotated cell types. Red = ex utero-favored pathways, blue = in utero-favored pathways, white = balanced metabolism, gray with hatching = insufficient data.

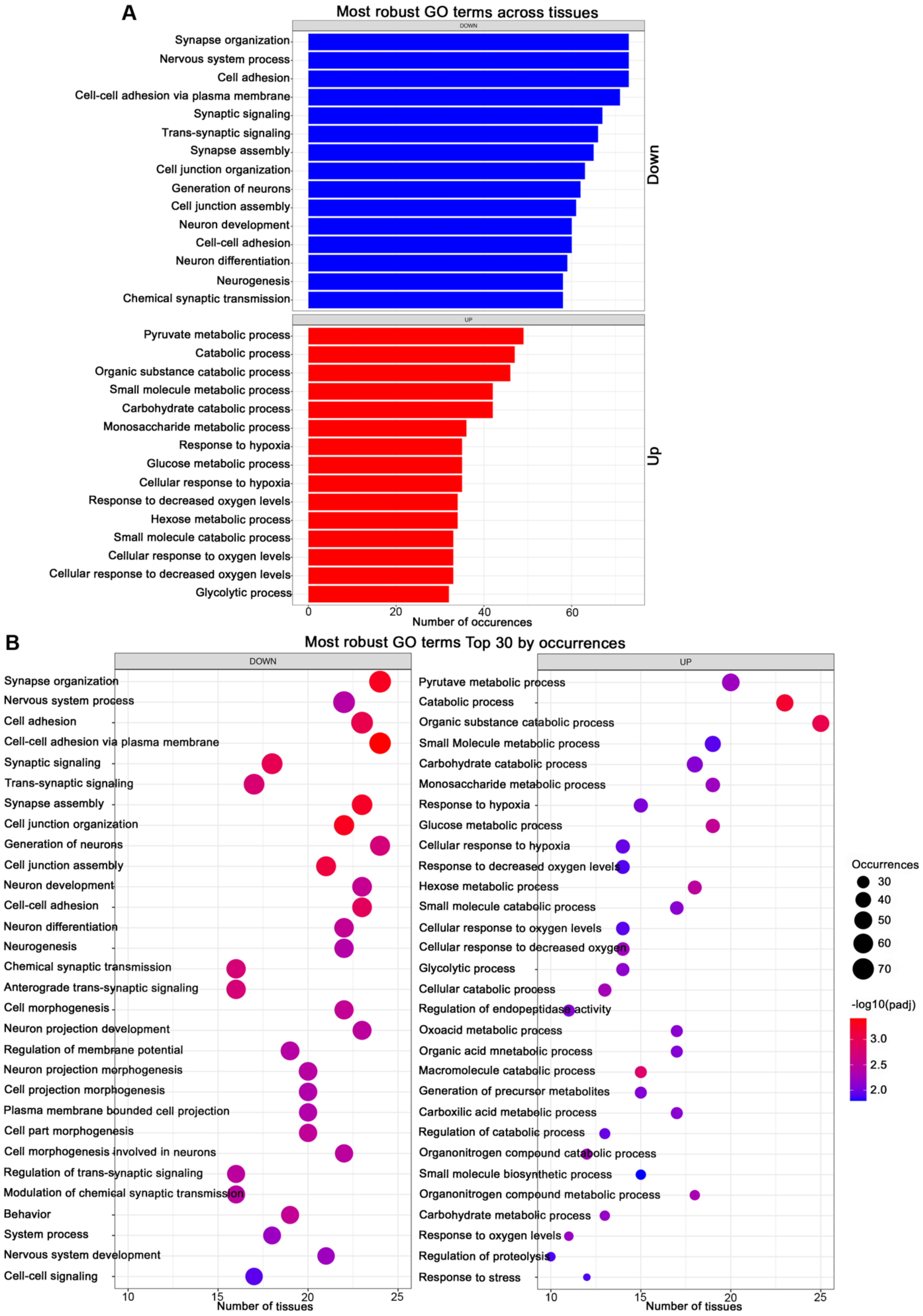
A
Most robust GO terms across tissues
DOWN
Synapse organization
Nervous system process
Cell adhesion
Cell-cell adhesion via plasma membrane
Synaptic signaling
Trans-synaptic signaling
Synapse assembly
Cell junction organization
Generation of neurons
Cell junction assembly
Neuron development
Cell-cell adhesion
Neuron differentiation
Neurogenesis
Chemical synaptic transmission
Down
UP
Pyruvate metabolic process
Catabolic process
Organic substance catabolic process
Small molecule metabolic process
Carbohydrate catabolic process
Monosaccharide metabolic process
Response to hypoxia
Glucose metabolic process
Cellular response to hypoxia
Response to decreased oxygen levels
Hexose metabolic process
Small molecule catabolic process
Cellular response to oxygen levels
Cellular response to decreased oxygen levels
Glycolytic process
Up
0
20
40
60
Number of occurences
B
Most robust GO terms Top 30 by occurrences
DOWN
Synapse organization
Nervous system process
Cell adhesion
Cell-cell adhesion via plasma membrane
Synaptic signaling
Trans-synaptic signaling
Synapse assembly
Cell junction organization
Generation of neurons
Cell junction assembly
Neuron development
Cell-cell adhesion
Neuron differentiation
Neurogenesis
Chemical synaptic transmission
Anterograde trans-synaptic signaling
Cell morphogenesis
Neuron projection development
Regulation of membrane potential
Neuron projection morphogenesis
Cell projection morphogenesis
Plasma membrane bounded cell projection
Cell part morphogenesis
Cell morphogenesis involved in neurons
Regulation of trans-synaptic signaling
Modulation of chemical synaptic transmission
Behavior
System process
Nervous system development
Cell-cell signaling
10
15
20
25
Number of tissues
UP
Pyrutave metabolic process
Catabolic process
Organic substance catabolic process
Small Molecule metabolic process
Carbohydrate catabolic process
Monosaccharide metabolic process
Response to hypoxia
Glucose metabolic process
Cellular response to hypoxia
Response to decreased oxygen levels
Hexose metabolic process
Small molecule catabolic process
Cellular response to oxygen levels
Cellular response to decreased oxygen
Glycolytic process
Cellular catabolic process
Regulation of endopeptidase activity
Oxoacid metabolic process
Organic acid mnetabolic process
Macromolecule catabolic process
Generation of precursor metabolites
Carboxilic acid metabolic process
Regulation of catabolic process
Organonitrogen compound catabolic process
Small molecule biosynthetic process
Organonitrogen compound metabolic process
Carbohydrate metabolic process
Response to oxygen levels
Regulation of proteolysis
Response to stress
10
15
20
25
Number of tissues
Occurrences
30
40
50
60
70
-log10(padj)
3.0
2.5
2.0

**Extended data Fig. 22. Gene Ontology enrichment analysis of differentially expressed genes.** **A,** Bar plot showing the most frequently enriched GO terms (Biological Process) across all analyzed cell types. Blue bars: GO terms enriched in downregulated genes (higher in in utero, labeled "Down"). Red bars: GO terms enriched in upregulated genes (higher in ex utero, labeled "Up"). X-axis indicates the number of cell types (tissues) showing significant enrichment for each GO term. Top enriched processes in downregulated genes include synapse organization, nervous system process, cell adhesion, and synaptic signaling. Top enriched processes in upregulated genes include pyruvate metabolic process, catabolic process, organic substance catabolic process, and various metabolic processes. **B,** Dot plot showing the top 30 most robust GO terms across tissues. Left panel: GO terms for downregulated genes. Right panel: GO terms for upregulated genes. Dot size indicates the number of tissues showing enrichment, dot color indicates -log10(adjusted p-value). This aggregated analysis reveals biological processes consistently affected by ex utero culture conditions across multiple cell types.
